# The within-subject stability of cortical thickness, surface area, and brain volumes across one year

**DOI:** 10.1101/2024.06.01.596956

**Authors:** Meng-Yun Wang, Max Korbmacher, Rune Eikeland, Stener Nerland, Didac Vidal-Pineiro, Karsten Specht

## Abstract

T1-weighted (T1w) imaging is widely used to examine brain structure based on image-derived phenotypes (IDPs) such as cortical thickness, surface area, and brain volumes. The reliability of these IDPs has been extensively explored, mainly focusing on the inter-subject variations, whereas the stability of the within-subject variations has often been overlooked. Additionally, how environmental factors such as time of day and daylight hours impact the structural brain is poorly understood. Therefore, we aimed to address the stability of T1w-derived phenotypes and explore the effecting factors including processing stream, brain region size, time of day, daylight hours, and head movement.

Three subjects in their late 20s, early 30s, and early 40s were scanned repeatedly on the same scanner over one year, from which a densely sampled dataset was acquired with 38, 40, and 25 sessions for subjects 1, 2, and 3, respectively. The temporal stability was evaluated using within-subject percentage change and coefficients of variation (CV). The effects of time of day and daylight hours were assessed by fitting general linear models, aggregating the effect size with meta-analysis, and equivalent analysis.

The longitudinal processing stream generates more stable results than the cross-sectional processing stream, hence the following results are all derived from the longitudinal processing stream. First and foremost, most IDPs demonstrated percentage changes within 5% and CVs within 2% for almost all brain regions, indicating high stability. Interestingly, we found that the brain region size negatively correlates with the CVs. Specifically, several small brain regions, including the temporal pole, frontal pole, pericalcarine, entorhinal cortex, and accumbens area, showed low stability. In addition, cortical thickness change was strongly and positively correlated with that of volume change while being negatively correlated with change in surface area, illustrating their distinct roles in brain anatomy. Moreover, the time of day could be ignored when evaluating the total surface area and total cortical brain volume but not the average cortical thickness and total subcortical brain volume. Furthermore, daylight hours could be left out when evaluating IDPs since there was no appreciable effect of daylight hours on the IDPs stability. Lastly, apparent head motion causes cortical thickness and volume underestimated and surface area overestimated.

## 1 Introduction

Magnetic resonance imaging (MRI) has become widely utilized to study brain structure, due to its ability to capture in vivo human brain images (Lerch et al., 2017). Commonly used T1-weighted (T1w) image-derived phenotypes (IDPs), including cortical thickness, surface area, and cortical as well as subcortical brain volumes, can be estimated with FreeSurfer, which is the most commonly used package for surface-based analysis (Dale, Fischl, & Sereno, 1999; Fischl, Sereno, & Dale, 1999). These IDPs have been ubiquitously used to investigate longitudinal brain changes (Bethlehem et al., 2022; Lemaitre et al., 2012; Sowell et al., 2003; Sowell et al., 2004; Vijayakumar et al., 2016), associations with other phenotypes (Chen et al., 2012; L. T. Elliott et al., 2018; Genon, Eickhoff, & Kharabian, 2022; Grasby et al., 2020; Holz et al., 2023; Wu, Li, Eickhoff, Scheinost, & Genon, 2023), and brain abnormalities in a variety of clinical conditions (Thompson et al., 2020).

However, before its grand application, the reliability and stability of the measurements should be articulated (Brandmaier et al., 2018). Reliability represents the contribution of the between-subject variance in terms of the total variance (sum of between- and within-subject variances) (Brandmaier et al., 2018; Zuo, Xu, & Milham, 2019) whereas stability refers to the precision of the within-subject estimations (Brandmaier et al., 2018). The reliability of the MRI measurements, which can be quantified with the intra-class correlation coefficient (ICC), has been extensively explored (Brandmaier et al., 2018; M. L. Elliott et al., 2020; Noble, Scheinost, & Constable, 2021; Noble et al., 2017; Parsons, Brandmaier, Lindenberger, & Kievit, 2024; Vidal-Piñeiro et al., 2024; Zuo et al., 2019). However, the stability of the MRI measurements has been largely ignored because of the scarcity of datasets with sufficient data points for each subject, which often leads to the presumption of taking the within-subject variance as a random error or noise (Brandmaier et al., 2018; Zuo et al., 2019).

Since the insufficiency of IDPs stability studies, it is still largely unknown which factor could sway the IDPs stability. Previous studies suggest that, among all the factors, different FreeSurfer versions (Gronenschild et al., 2012), operating systems (Gronenschild et al., 2012), scanners (Han et al., 2006; Iscan et al., 2015), processing streams (Reuter, Schmansky, Rosas, & Fischl, 2012; Vidal-Piñeiro et al., 2024), time of day (Karch et al., 2019; Nakamura et al., 2015; Trefler et al., 2016), daylight hours (Di, Woelfer, Kühn, Zhang, & Biswal, 2022), and head movement (Alexander-Bloch et al., 2016; Reuter et al., 2015) could affect the reliability or stability of the IDPs. The factor of the operating system and scanner can be controlled by a within-subject design, and the FreeSurfer version can be controlled by using the same version to process all the datasets. Thereafter, the remaining factors compassing processing stream, time of day, daylight hours, and head movement will be primarily investigated in this study.

There are two processing streams in FreeSurfer to calculate IDPs, namely the cross-sectional and longitudinal processing streams (Reuter et al., 2012). The cross-sectional stream processes each scan independently while the longitudinal stream creates an unbiased within-subject template space and estimates IDPs based on all available data for each participant (Reuter et al., 2012). However, the impact of the two streams on the within-subject stability level has not been fully uncovered. The stability of those two streams, therefore, will be directly compared in this study.

Although time-of-day effects have been documented in functional brain organization studies (Orban, Kong, Li, Chee, & Yeo, 2020), the influence on brain structure is still elusive. Some studies have reported associations between time of day and cortical measures (Karch et al., 2019; Nakamura et al., 2015; Trefler et al., 2016). Specifically, the cortical thickness and brain volume are larger in the morning compared to the afternoon (Karch et al., 2019; Nakamura et al., 2015; Trefler et al., 2016). However, two of the studies (Nakamura et al., 2015; Trefler et al., 2016) employed a cross-sectional design where between-subject variation could interfere with the reported time-of-day effect. For example, in one study, each subject only had two data points in their primary dataset for morning and afternoon sessions (Trefler et al., 2016), and the other study just pooled the data from other datasets and separated them based on the collection time (Nakamura et al., 2015). Although one is within-subject design (Karch et al., 2019), they did not use the longitudinal stream as recommended (Reuter et al., 2012), which could hinder the robustness and reliability of the results. More critically, the data distributions of the morning and afternoon sessions within subjects were not illustrated. Therefore, in this study, within-subject design together with the longitudinal processing stream will be adopted meanwhile the data distribution will be provided.

Besides the time-of-day effect, the season of the year has been suggested to affect functional brain organization (M. Y. Wang, Korbmacher, Eikeland, & Specht, 2023b; Zhang, Shokri-Kojori, & Volkow, 2023). Specifically, daylight hours comes out as a potential contributor (Di et al., 2022) to the functional brain organization (M. Y. Wang et al., 2023b). For example, the salience network, one of the resting-state brain networks, is enlarged when daylight hours is short compared to when it is long (M. Y. Wang et al., 2023b). To the best of our knowledge, there is still no study investigating the daylight-hour effect on brain structure. Located right within and beneath the Arctic Circle, the daylight hours in Norway varies substantially throughout the year. Specifically, the daylight hours in the city of Bergen can be as short as around 2 hours of daylight and as long as around 16 hours of daylight, which makes it suitable for exploring the daylight-hour effect.

Low brain image quality can introduce bias which can lead to inaccurate and even erroneous results (Alexander-Bloch et al., 2016; Ducharme et al., 2016; Reuter et al., 2015). Brain images with obvious artifacts, such as field of view lacking parts of the brain, ringing artifacts, and image distortions are routinely discarded after visual inspection and are not used in further analysis. However, images that have passed the visual inspection can still influence the downstream processing. For example, the head movement can reduce gray matter volume and cortical thickness estimations (Alexander-Bloch et al., 2016; Reuter et al., 2015) and even micro-movement can influence the longitudinal brain change evaluations (Alexander-Bloch et al., 2016). However, a systematic overview of how within-subject head movement affects the estimation of IDPs is still lacking.

Additionally, the influence of different brain region sizes on IDPs stability is still unknown. It has become common practice to report brain imaging results in a common brain atlas (Desikan et al., 2006; Yeo et al., 2011) to aggregate or compare findings across different studies. Among these atlases, the Desikan-Killiany (DK) atlas (Desikan et al., 2006) has become the most used atlas in the T1w brain imaging studies (Bethlehem et al., 2022; Chen et al., 2012; L. T. Elliott et al., 2018; Genon et al., 2022; Grasby et al., 2020; Lemaitre et al., 2012; Vijayakumar et al., 2016). The DK atlas was developed based on curvature-based information such as the sulcal/gyral representations and contains 34 cortical brain regions in each hemisphere (Desikan et al., 2006). In this study, therefore, IDPs were extracted from those 68 brain regions. Additionally, 7 subcortical regions (thalamus, pallidum, amygdala, hippocampus, putamen, accumbens area, and caudate nucleus) based on the *aseg* atlas were used for the subcortical brain volume analysis. The sizes of these 68 brain regions are, however, distinct from each other indicated by their volumes or surface areas. Therefore, the brain region size effect on the stability will also be explored.

The main aims of this study were twofold. First, to characterize the stability of IDPs, including cortical thickness, surface area, and brain volumes. Second, to investigate the influence of potential confounding variables on the stability of IDPs, specifically the effects of processing streams, brain region size, time of day, daylight hours, and head movement.

## 2 Methods

### 2.1 Participants

The ethics approval was granted by the Regional Committees for Medical and Health Research Ethics (REK-vest). Informed consent was obtained from all participants for being included in the study. Here, we collected a densely sampled brain imaging dataset derived from the BBSC (Bergen Breakfast Scanning Club) project (M. Y. Wang, Korbmacher, Eikeland, & Specht, 2022, 2023a). In this project, three subjects (**Table 1**) were repeatedly scanned in the same scanner located at the same place spanning over one year, which makes it suitable to evaluate IDPs stability (M. Y. Wang, Korbmacher, Eikeland, Craven, & Specht, 2024; M. Y. Wang et al., 2022). The BBSC project aims to illustrate the individual precision brain networks and to examine the stability of both functional and structural MRI measurements over one year (M. Y. Wang et al., 2022). For this purpose, the participants were scanned twice a week between February 2021 and February 2022 with two breaks (Jun. to Oct. 2021, and Jan. 2022) where functional and structural MRI data (M. Y. Wang et al., 2024; M. Y. Wang et al., 2023a) were collected. In total, there are 38, 40, and 25 T1w MRI sessions for subjects 1, 2, and 3, respectively. The exact dates of these sessions can be found in **Table S1.**

**Table 1.** Basic demographic information of subjects.

|  | Sub1 | Sub2 | Sub3 |
| --- | --- | --- | --- |
| **Gender** | Male | Male | Male |
| **Age at baseline** | 31 | 27 | 40 |
| **Laterality** | Right | Right | Right |
| **Regular caffeine consumption?** | No | Yes | Yes |
| **Regular nicotine consumption?** | No | No | No |

All three participants speak at least two languages (their native languages, and English). Notably, the first participant has also been acquiring an additional language since January 2021 and remained COVID-19 free during data collection. The second participant contracted COVID-19 around December 2021, while the third participant had COVID-19 in approximately August 2021.

### 2.2 Data collection

Data collection was embedded in the functional protocol of the BBSC project (M. Y. Wang et al., 2022, 2023a), which lasted around 25 minutes in total. The procedure of the data collection in the functional protocol is seven mins T1w MRI, 5 mins MR spectroscopy (MRS) data, and 12 mins rs-fMRI, where the data were collected with a 3T MRI scanner (GE Discovery MR750) with a 32-channel head coil at the Haukeland University Hospital in Bergen, Norway. The other modalities in the BBSC project have been described previously (Korbmacher et al., 2023; M. Y. Wang et al., 2024; M. Y. Wang et al., 2023b).

Seven-minute structural T1w images were acquired using a 3D Fast Spoiled Gradient-Recalled Echo (FSPGR) sequence with the following parameters: 188 contiguous slices acquired, with repetition time (TR) = 6.88 ms, echo time (TE) = 2.95 ms, flip angle (FA) = 12°, slice thickness = 1 mm, in-plane resolution = 1 mm × 1 mm, and field of view (FOV) = 256 mm, with an isotropic voxel size of 1 mm^3^.

### 2.3 Data processing

Cortical thickness and surface area, as well as the cortical and subcortical volumes were computed with FreeSurfer v7.2 (freesurfer-darwin-macOS-7.2.0-20210713-aa8f76b) (Dale et al., 1999; Fischl et al., 1999). First, each session was processed with the cross-sectional *recon-all* stream. Then, the results went through the longitudinal stream (Reuter et al., 2012), where an unbiased within-subject template was created by using a robust, inverse consistent registration (Reuter, Rosas, & Fischl, 2010). For more details on data processing, see the **Supplementary Materials**.

Moreover, the T1w images were quality assessed with MRIQC 23.1.0 (Esteban et al., 2017), where several image quality metrics (IQMs) were generated. Head movement was evaluated using coefficient of joint variation (CJV) values, which was proposed as a measure for INU correction algorithms. Higher CJV indicates head motion or INU artifacts, whereas lower values imply better image quality (Ganzetti, Wenderoth, & Mantini, 2016). From the data distribution of CJV, it is noticeable that there were two sessions (sessions 1 and 7) with excessive values in sub3. Accordingly, those two sessions were excluded from evaluating cortical IDPs but not subcortical ones, since the impact was negligible on subcortical brain volume (**Figs. S11-S12**). The IQMs were described in the image quality checking part of the **Supplementary Materials**.

### 2.4 Data Analysis

#### 2.4.1 Coefficient of variation

For each subject, stability was quantified with the coefficient of variation (CV), defined as the ratio of an estimate of variability and a mean, and it estimates the dispersion of the different measurement values to the average values (Brandmaier et al., 2018; M. Y. Wang et al., 2024). The CV values are always positive with smaller values indicating higher stability (Brandmaier et al., 2018; M. Y. Wang et al., 2024). We computed CVs as follows:

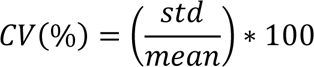

where *std* denotes the standard deviation of the measurements taken across sessions for the same subject, while *mean* refers to the average of the within-subject measurements.

CVs facilitate the comparison of variability across varying units or scales (Brandmaier et al., 2018). It has also been instrumental in assessing the stability of measurements over time, where it is especially suitable for the within-subject design (Brandmaier et al., 2018; M. Y. Wang et al., 2024). Lower CVs indicate less variability relative to the mean, signifying higher stability or precision. Conversely, higher CVs indicate more variability relative to the mean, suggesting lower stability or precision (Brandmaier et al., 2018; M. Y. Wang et al., 2024).

#### 2.4.2 Percentage change

To account for discrepancies among different brain regions, the percentage changes instead of raw values of each IDP were used to illustrate the discrepancy (e.g., **Figs 2-5, 8**), which were calculated as the deviation from the mean divided by the mean itself. Furthermore, to explore which brain regions covaried, correlation matrices were computed within and between each phenotype.

#### 2.4.3 Time-of-day & daylight length effects

Time-of-day was classified based on the data collection time, labeling sessions before noon as ‘morning’ and those after as ‘afternoon’. The distribution of sessions was as follows: in sub1, there were 22 morning and 16 afternoon sessions; in sub2, 23 morning and 16 afternoon sessions; and in sub3, 14 morning and 9 afternoon sessions. Daylight hour data, ranging from 120 to 948 minutes, was obtained from the weather station in the Florida area of Bergen maintained by the University of Bergen (https://veret.gfi.uib.no/). The time-of-day indices and daylight hours were described in **Table S1**.

Regarding the time-of-day and daylight-hour effects, to increase the statistical power, the effect size calculated within each subject will be aggregated using a meta-analysis (Viechtbauer, 2010). In addition, to explore whether the effect size is too trivial to be considered regardless of its significance from meta-analysis, equivalent tests will be implemented (Lakens, 2017; Lakens, Scheel, & Isager, 2018). The rationale of the equivalent test is straightforward: if the observed effect size of one factor is significantly different from a predefined effect size (i.e., 0.5), it then can be deduced that this effect size is too trivial (< 0.5), and the factor can be left out in future experiment designs (Lakens, 2017; Lakens et al., 2018; Vidal-Pineiro et al., 2021). Otherwise, it cannot be ruled out that there is no actual effect of that factor (Lakens, 2017; Lakens et al., 2018).

Accordingly, the following steps were implemented to evaluate the time-of-day effect and the daylight-hour effect on the IDPs. Here, for the sake of simplicity of the general linear model, the IDPs in this analysis only include the average cortical thickness, total surface area, and total cortical as well as subcortical brain volumes, instead of looking at the 68 regions separately.

First, the general linear model was fitted within each subject to examine the statistical significance of each subject, where the normalized values of each IDP were designated as dependent variables and time of day or daylight hours was considered as the independent variables:

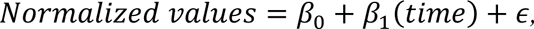

where *β_0_* is the intercept; *β_1_* is the fixed effect coefficients for each predictor; *time* represents time of day or daylight hours, and *ɛ* is the residual error.

Second, the beta values and standard deviation errors (SE) derived from the linear model were fed into the *rma* function in the package Metafor (v4.6.0) (Viechtbauer, 2010) to aggregate estimates of the strength of the effects.

Third, the equivalence test was computed with the package TOSTER (V0.8.3) (Lakens, 2017; Lakens et al., 2018) based on the meta-analysis results to examine the practical significance meaning whether the effect was too trivial to ignore (Lakens, 2017; Lakens et al., 2018; Vidal-Pineiro et al., 2021). Here, we set the equivalence bounds for median effect size (d) of - 0.50 and 0.50 at a type I error (alpha) of 0.05.

#### 2.4.4 Head movement effect

To assess the influence of head movement on the stability of IDPs, paired *t*-tests were executed between the CV values of sessions before and after quality checking. Specifically, data from sub3 were used since two sessions were excluded because of the presence of excessive head movements.

#### 2.4.5 Brain region size

To investigate the influence of different brain region sizes exerting on stability, the brain region sizes indicated by their volume and CVs were normalized within each subject and each hemisphere, leading to 204 (34*2*3) for thickness, surface area as well as cortical volume, and 42 (7*2*3) data points for subcortical volume. Then, Pearson’s correlation between normalized brain region size and CVs was calculated.

Statistical analyses and visualizations were done with R 4.3.2 (R Core Team, 2022), and visualization was performed with ggplot2 (v3.5.1) (Wickham, 2009), ggseg (v1.6.6) (Mowinckel & Vidal-Piñeiro, 2020), and patchwork (v1.2.0).

## 3 Results

### 3.1 Longitudinal vs cross-sectional streams

The CVs of the longitudinal and cross-sectional processing streams on cortical brain volumes from FreeSurfer were described in **Table 2**. Meanwhile, the percentage changes against the average of different brain tissues for each subject were illustrated in **Fig. 1**.

**Fig. 1.**
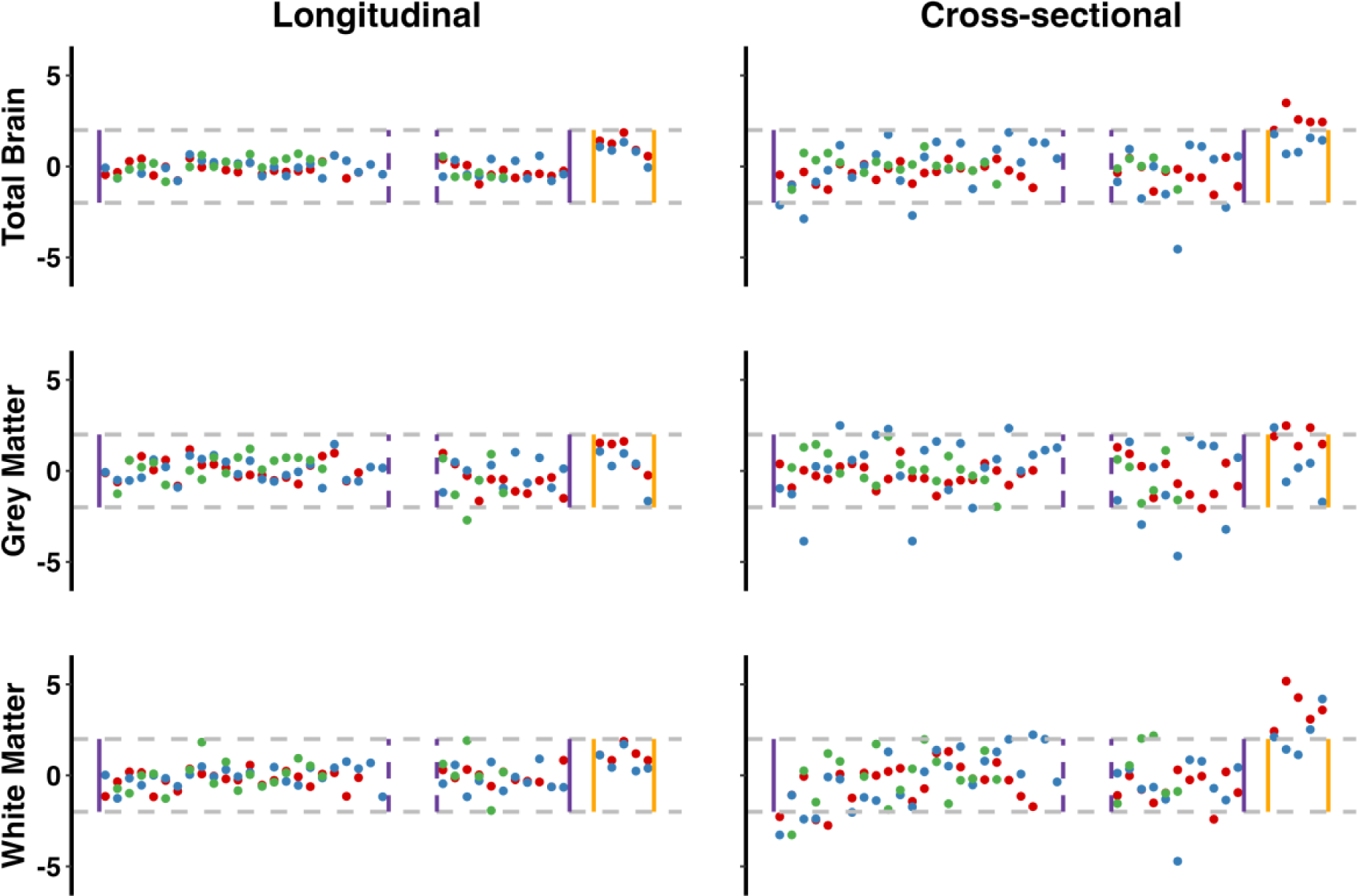
Percentage changes of different brain volumes from different processing streams. Percentage changes along the data collections generated from the longitudinal and cross-sectional processing streams. The x-axis represents the time sequence of data collection while the y-axis depicts the percentage change. The purple solid vertical lines denote that the data were collected in the same year whereas the yellow solid vertical lines represent the data were collected the next year. In addition, the gap between the two dashed purple lines represents the summer break while the gap between the purple and yellow solid lines indicates the winter break. The gray horizontal dashed lines denote 2% percentage changes. The red color represents subject 1, the blue depicts subject 2, and the green represents subject 3.

**Table 2.** The longitudinal stream generates smaller CVs.

|  | Sub1 |  | Sub2 |  | Sub3 |  |
| --- | --- | --- | --- | --- | --- | --- |
|  | Longitudinal | Cross-sectional | Longitudinal | Cross-sectional | Longitudinal | Cross-sectional |
| **Total Brain** | 0.61 | 1.16 | 0.54 | 1.48 | 0.47 | 0.58 |
| **Gray Matter** | 0.81 | 1.06 | 0.71 | 1.86 | 0.94 | 1.02 |
| **Left - GM** | 0.89 | 1.17 | 0.80 | 1.97 | 1.10 | 1.14 |
| **Right - GM** | 0.91 | 1.13 | 0.74 | 1.84 | 0.90 | 1.03 |
| **White Matter** | 0.66 | 1.79 | 0.65 | 1.74 | 0.89 | 1.42 |
| **Left - WM** | 0.69 | 1.86 | 0.63 | 1.76 | 1.22 | 1.49 |
| **Right - WM** | 0.76 | 1.77 | 0.74 | 1.76 | 0.81 | 1.51 |

All CVs are reported as percentages, where smaller is better. Total Brain: ‘BrainSegVolNotVent’ generated from FreeSurfer.

As shown in **Table 2 and Fig. 1**, the CVs and percentage changes generated by the longitudinal processing stream were smaller than those of the cross-sectional stream, which indicates the longitudinal stream generated more stable results. Therefore, the following analyses were based on the results derived from the longitudinal processing stream.

### 3.2 The stability of IDPs

#### 3.2.1 Cortical thickness

The raw values (**Table S2**) and characteristics (**Fig. S2**) of cortical thickness were described in the **Supplementary Materials**. The CVs of within-subject thickness ranged from 0.67 to 4.40 across all subjects, where most brain regions had CVs below 2. Specifically, the CVs vary from 0.74 to 2.64 for sub1, 0.67 to 3.24 for sub2, and 0.84 to 4.40 for sub3 (**Table 3**). Furthermore, the percentage changes in almost all brain regions across the three subjects were well-constrained within 5% as illustrated in **Fig. 2**. Several brain regions, however, showed larger CVs and percentage changes across all three subjects (**Fig. 2** and **Table 3**), such as the temporal pole, frontal pole, pericalcarine, and entorhinal cortex.

**Fig. 2.**
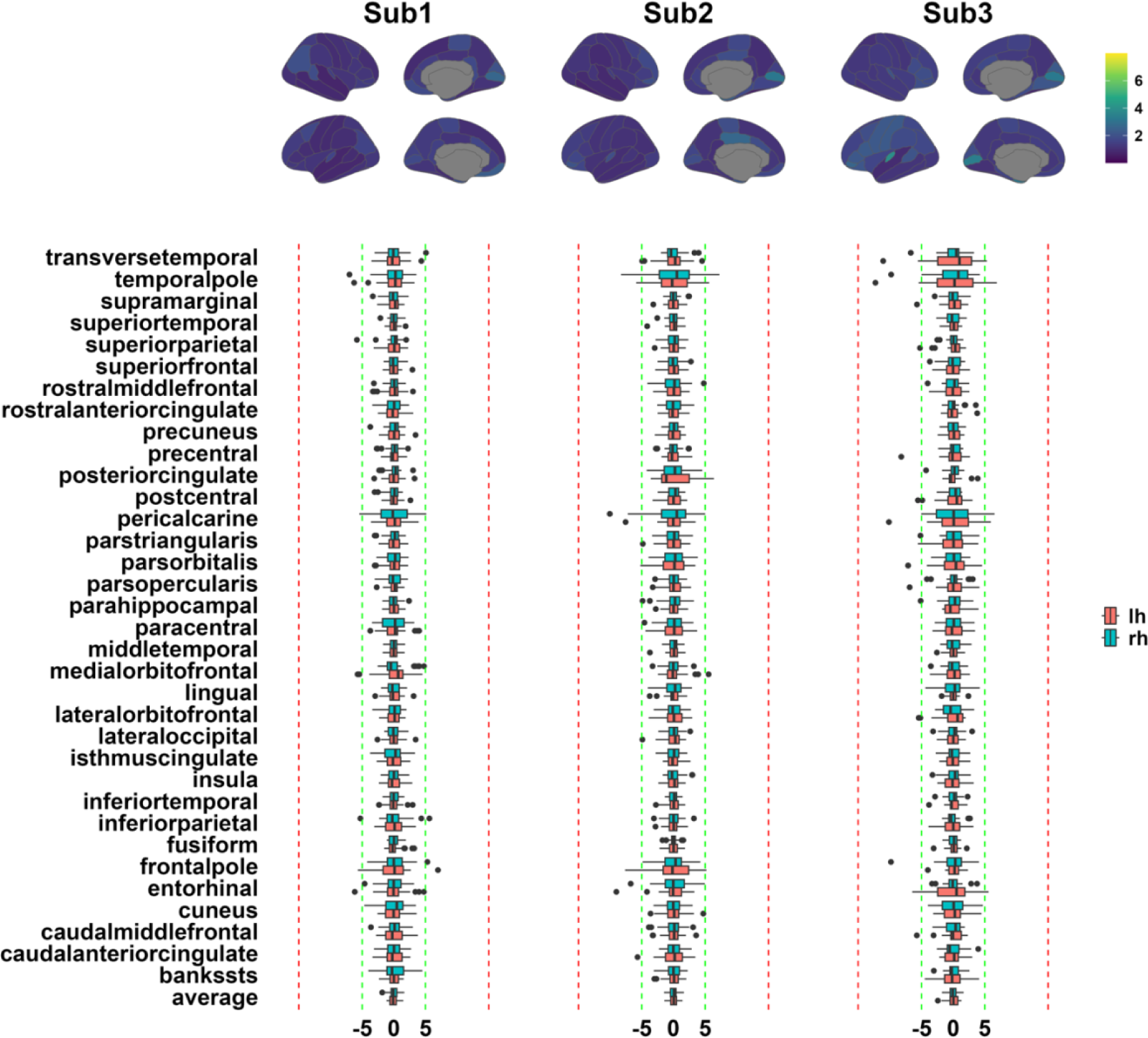
The CVs and percentage changes in the cortical thickness. The upper panel depicts the coefficient of the variation (CV) for each subject. The lower boxplots show percentage change against the average for each brain region and each subject. The green lines are in the range of ±5 % while the red lines depict the range of ±15%. All values in the boxplot are given as percentages.

**Table 3.**
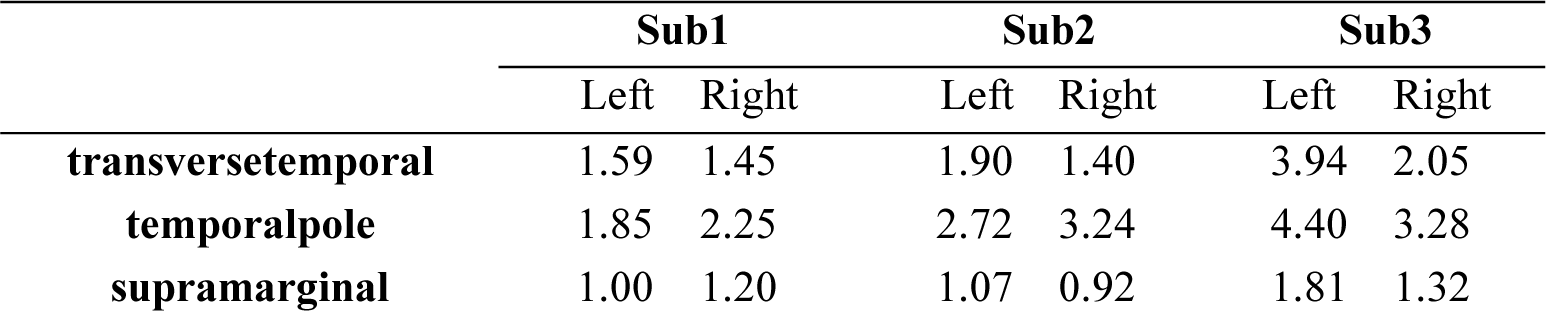
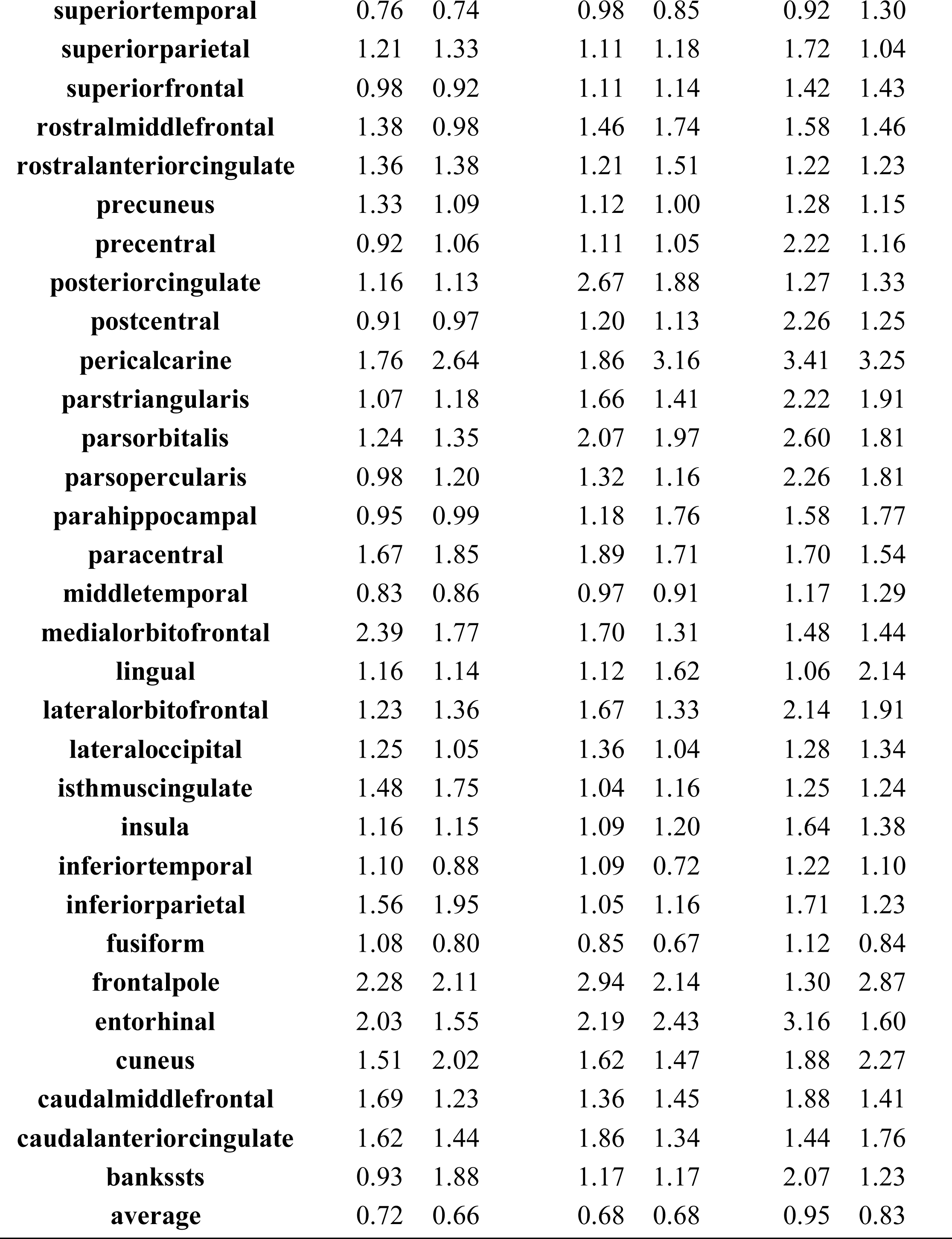
The CVs of the cortical thickness based on the DK atlas.

#### 3.2.2 Surface area

The raw values (**Table S3**) and characteristics (**Fig. S4**) of the surface area were described in the **Supplementary Materials**. The CVs of the within-subject area ranged from 0.41 to 7.25, specifically 0.41 to 4.52 for Sub1, 0.49 to 7.25 for Sub2, and 0.53 to 5.34 for Sub3 (**Table 4**). Subject 2 manifested a generally larger percentage change in the cingulate cortex (**Fig. 3**) and greater CVs (**Table 4**) than other brain regions along the data collections. Aside from the cingulate cortex in sub2, almost all brain regions showed percentage changes within 5% across all three subjects (**Fig. 3**). Notably, several surface areas showed larger CVs and percentage changes across all three subjects, including the temporal pole, frontal pole, and entorhinal cortex (**Fig. 3** and **Table 4**).

**Fig. 3.**
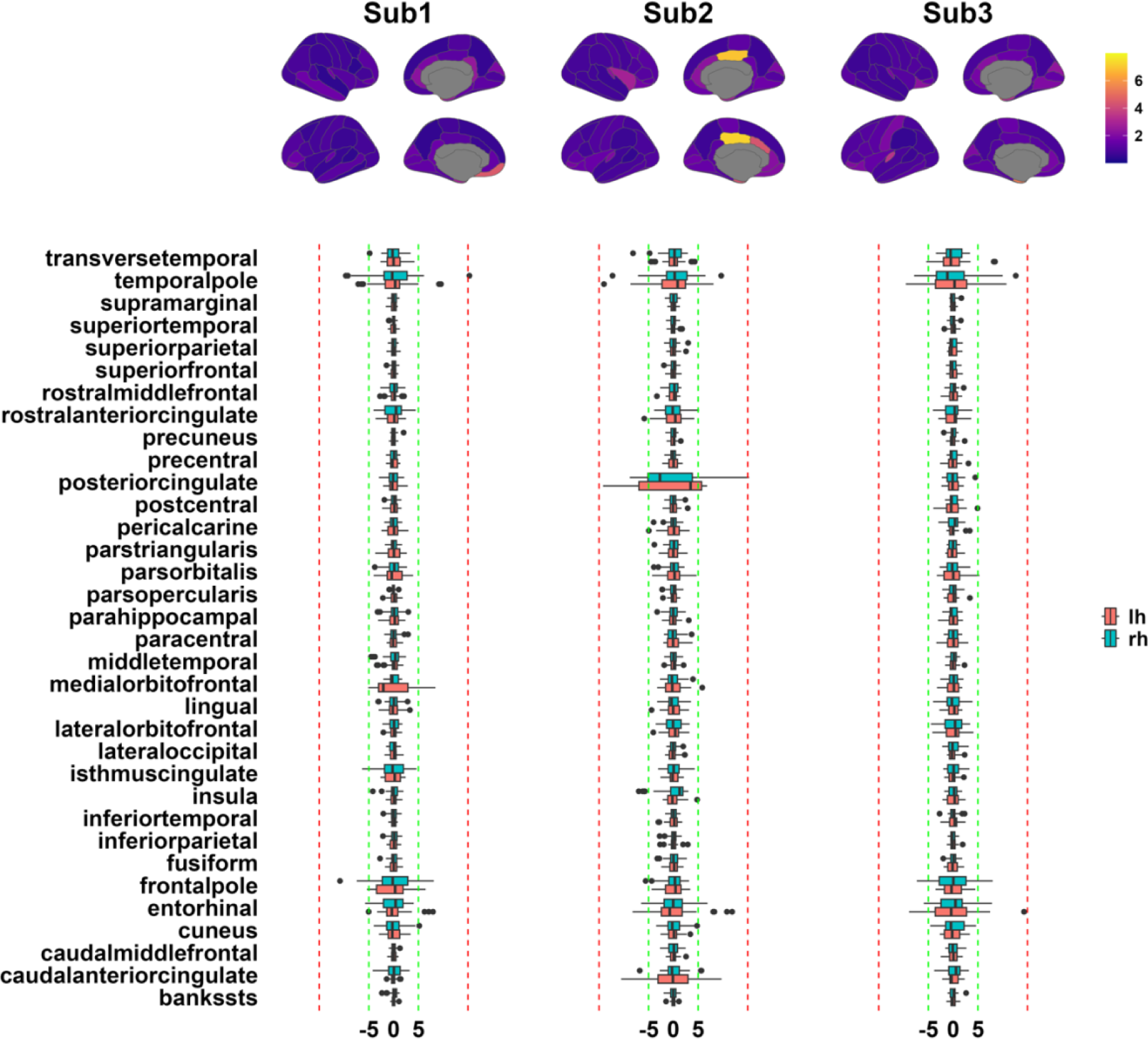
The CVs and percentage changes in surface area. The upper panel depicts the coefficient of variation (CV) for each subject. The lower boxplot shows percentage changes for each brain region and each subject. The green lines are in the range of ±5 % while the red lines depict the range of ±15%. The unit of all values in the boxplot is percentage.

**Table 4.**
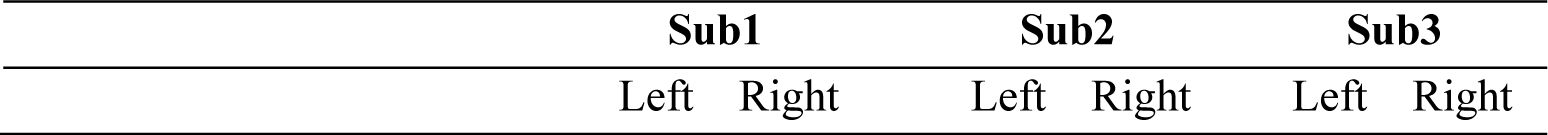
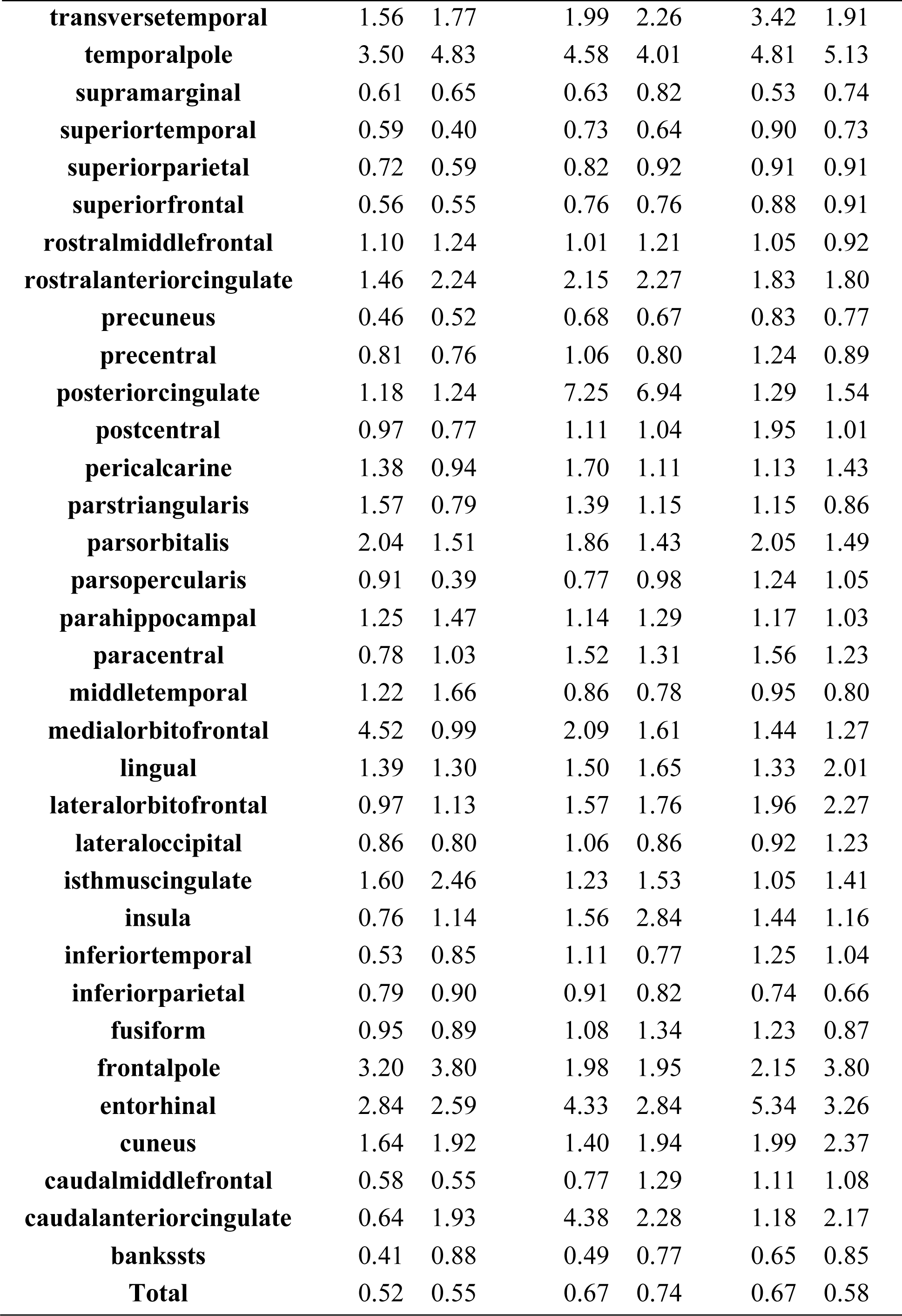
The CVs of the surface area based on the DK atlas.

|  |  |  |  |  |  |  |
| --- | --- | --- | --- | --- | --- | --- |
| **transversetemporal** | 1.56 | 1.77 | 1.99 | 2.26 | 3.42 | 1.91 |
| **temporalpole** | 3.50 | 4.83 | 4.58 | 4.01 | 4.81 | 5.13 |
| **supramarginal** | 0.61 | 0.65 | 0.63 | 0.82 | 0.53 | 0.74 |
| **superiortemporal** | 0.59 | 0.40 | 0.73 | 0.64 | 0.90 | 0.73 |
| **superiorparietal** | 0.72 | 0.59 | 0.82 | 0.92 | 0.91 | 0.91 |
| **superiorfrontal** | 0.56 | 0.55 | 0.76 | 0.76 | 0.88 | 0.91 |
| **rostralmiddlefrontal** | 1.10 | 1.24 | 1.01 | 1.21 | 1.05 | 0.92 |
| **rostralanteriorcingulate** | 1.46 | 2.24 | 2.15 | 2.27 | 1.83 | 1.80 |
| **precuneus** | 0.46 | 0.52 | 0.68 | 0.67 | 0.83 | 0.77 |
| **precentral** | 0.81 | 0.76 | 1.06 | 0.80 | 1.24 | 0.89 |
| **posteriorcingulate** | 1.18 | 1.24 | 7.25 | 6.94 | 1.29 | 1.54 |
| **postcentral** | 0.97 | 0.77 | 1.11 | 1.04 | 1.95 | 1.01 |
| **pericalcarine** | 1.38 | 0.94 | 1.70 | 1.11 | 1.13 | 1.43 |
| **parstriangularis** | 1.57 | 0.79 | 1.39 | 1.15 | 1.15 | 0.86 |
| **parsorbitalis** | 2.04 | 1.51 | 1.86 | 1.43 | 2.05 | 1.49 |
| **parsopercularis** | 0.91 | 0.39 | 0.77 | 0.98 | 1.24 | 1.05 |
| **parahippocampal** | 1.25 | 1.47 | 1.14 | 1.29 | 1.17 | 1.03 |
| **paracentral** | 0.78 | 1.03 | 1.52 | 1.31 | 1.56 | 1.23 |
| **middletemporal** | 1.22 | 1.66 | 0.86 | 0.78 | 0.95 | 0.80 |
| **medialorbitofrontal** | 4.52 | 0.99 | 2.09 | 1.61 | 1.44 | 1.27 |
| **lingual** | 1.39 | 1.30 | 1.50 | 1.65 | 1.33 | 2.01 |
| **lateralorbitofrontal** | 0.97 | 1.13 | 1.57 | 1.76 | 1.96 | 2.27 |
| **lateraloccipital** | 0.86 | 0.80 | 1.06 | 0.86 | 0.92 | 1.23 |
| **isthmuscingulate** | 1.60 | 2.46 | 1.23 | 1.53 | 1.05 | 1.41 |
| **insula** | 0.76 | 1.14 | 1.56 | 2.84 | 1.44 | 1.16 |
| **inferiortemporal** | 0.53 | 0.85 | 1.11 | 0.77 | 1.25 | 1.04 |
| **inferiorparietal** | 0.79 | 0.90 | 0.91 | 0.82 | 0.74 | 0.66 |
| **fusiform** | 0.95 | 0.89 | 1.08 | 1.34 | 1.23 | 0.87 |
| **frontalpole** | 3.20 | 3.80 | 1.98 | 1.95 | 2.15 | 3.80 |
| **entorhinal** | 2.84 | 2.59 | 4.33 | 2.84 | 5.34 | 3.26 |
| **cuneus** | 1.64 | 1.92 | 1.40 | 1.94 | 1.99 | 2.37 |
| **caudalmiddlefrontal** | 0.58 | 0.55 | 0.77 | 1.29 | 1.11 | 1.08 |
| **caudalanteriorcingulate** | 0.64 | 1.93 | 4.38 | 2.28 | 1.18 | 2.17 |
| **bankssts** | 0.41 | 0.88 | 0.49 | 0.77 | 0.65 | 0.85 |
| **Total** | 0.52 | 0.55 | 0.67 | 0.74 | 0.67 | 0.58 |

#### 3.2.3 Cortical brain volume

The raw values (**Tables S4 & S5**) and characteristics (**Fig. S6**) of the cortical brain volume were described in the **Supplementary Materials**. The CVs of the within-subject brain volume range from 0.64 to 4.73 for sub1, 0.72 to 4.87 for sub2, and 0.86 to 4.99 for sub3 (**Table 5**). Furthermore, several brain regions showed larger CVs (**Table 5**) and percentage changes (**Fig. 4**), such as the temporal pole, pericalcarine, and entorhinal cortex. Moreover, aside from the abovementioned regions, the percentage changes are well confined within 5% in all subjects (**Fig. 4**).

**Fig. 4.**
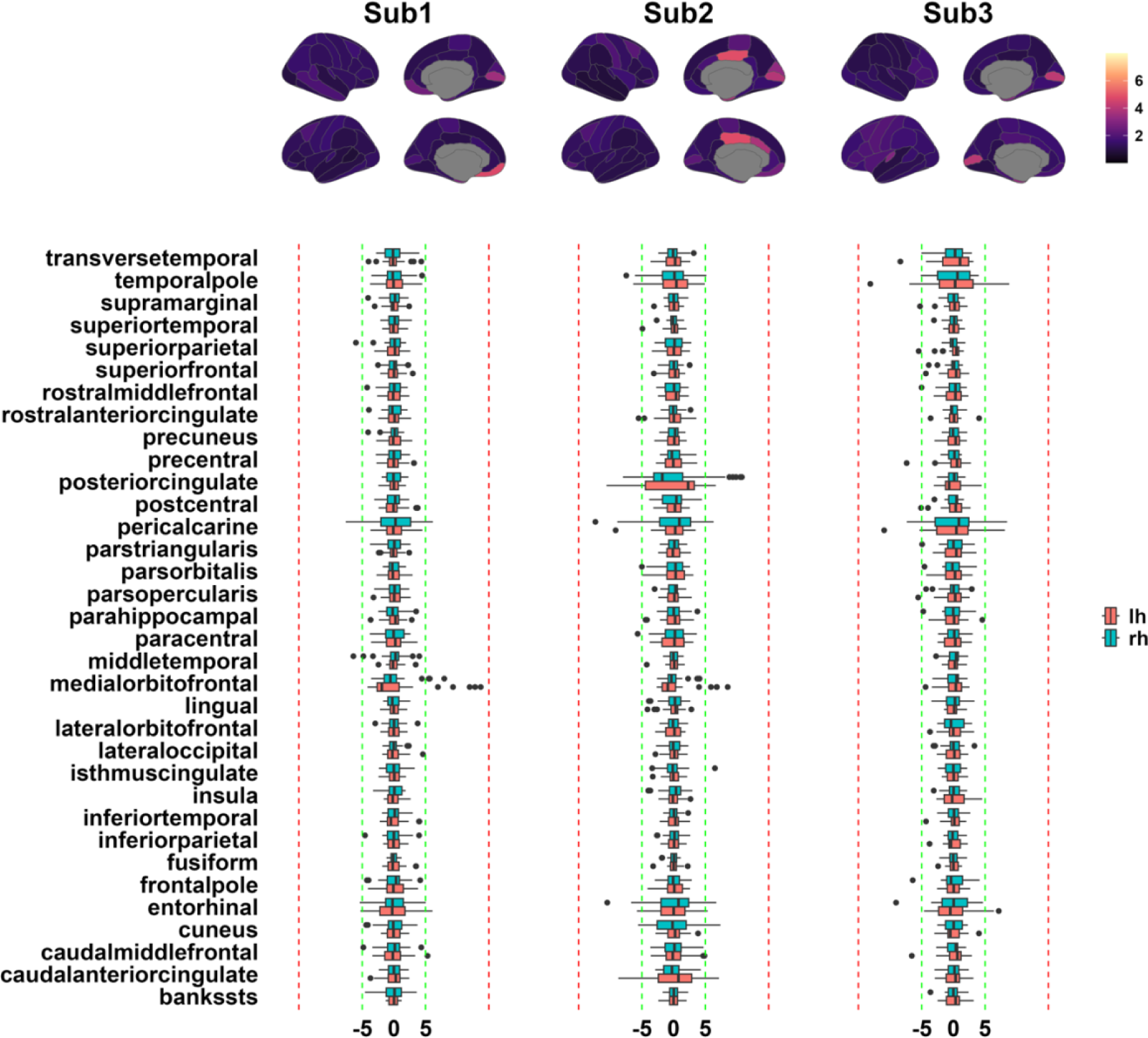
The CVs and percentage changes in cortical brain volume. The upper panel depicts the coefficient of variation (CV) for each subject. The lower boxplot shows the percentage change for each brain region and subject. The green lines are in the range of ±5 % while the red lines depict the range of ±15%. All values in the boxplot are reported as percentages.

**Table 5.**
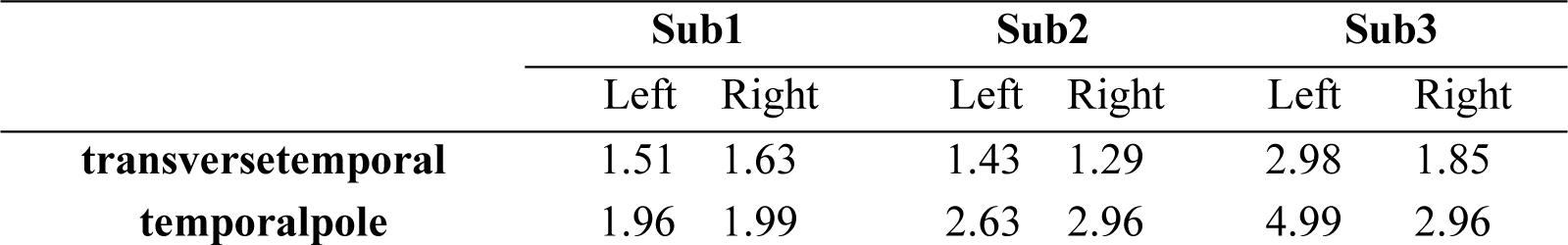
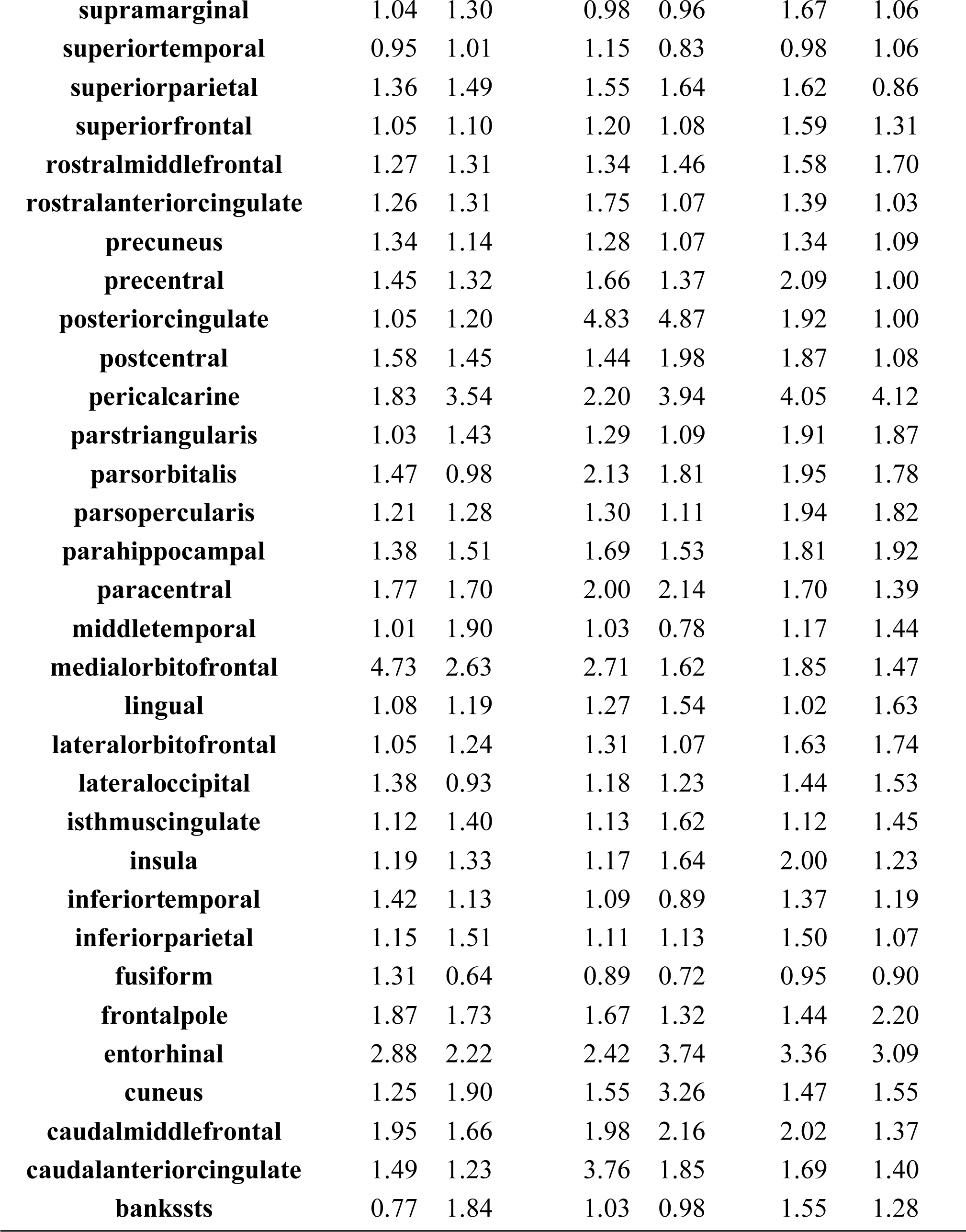
The CVs of the cortical brain volume based on the DK atlas.

|  | Sub1 |  | Sub2 |  | Sub3 |  |
| --- | --- | --- | --- | --- | --- | --- |
|  | Left | Right | Left | Right | Left | Right |
| transversetemporal | 1.51 | 1.63 | 1.43 | 1.29 | 2.98 | 1.85 |
| temporalpole | 1.96 | 1.99 | 2.63 | 2.96 | 4.99 | 2.96 |
| **supramarginal** | 1.04 | 1.30 | 0.98 | 0.96 | 1.67 | 1.06 |
| **superiortemporal** | 0.95 | 1.01 | 1.15 | 0.83 | 0.98 | 1.06 |
| **superiorparietal** | 1.36 | 1.49 | 1.55 | 1.64 | 1.62 | 0.86 |
| **superiorfrontal** | 1.05 | 1.10 | 1.20 | 1.08 | 1.59 | 1.31 |
| **rostralmiddlefrontal** | 1.27 | 1.31 | 1.34 | 1.46 | 1.58 | 1.70 |
| **rostralanteriorcingulate** | 1.26 | 1.31 | 1.75 | 1.07 | 1.39 | 1.03 |
| **precuneus** | 1.34 | 1.14 | 1.28 | 1.07 | 1.34 | 1.09 |
| **precentral** | 1.45 | 1.32 | 1.66 | 1.37 | 2.09 | 1.00 |
| **posteriorcingulate** | 1.05 | 1.20 | 4.83 | 4.87 | 1.92 | 1.00 |
| **postcentral** | 1.58 | 1.45 | 1.44 | 1.98 | 1.87 | 1.08 |
| **pericalcarine** | 1.83 | 3.54 | 2.20 | 3.94 | 4.05 | 4.12 |
| **parstriangularis** | 1.03 | 1.43 | 1.29 | 1.09 | 1.91 | 1.87 |
| **parsorbitalis** | 1.47 | 0.98 | 2.13 | 1.81 | 1.95 | 1.78 |
| **parsopercularis** | 1.21 | 1.28 | 1.30 | 1.11 | 1.94 | 1.82 |
| **parahippocampal** | 1.38 | 1.51 | 1.69 | 1.53 | 1.81 | 1.92 |
| **paracentral** | 1.77 | 1.70 | 2.00 | 2.14 | 1.70 | 1.39 |
| **middletemporal** | 1.01 | 1.90 | 1.03 | 0.78 | 1.17 | 1.44 |
| **medialorbitofrontal** | 4.73 | 2.63 | 2.71 | 1.62 | 1.85 | 1.47 |
| **lingual** | 1.08 | 1.19 | 1.27 | 1.54 | 1.02 | 1.63 |
| **lateralorbitofrontal** | 1.05 | 1.24 | 1.31 | 1.07 | 1.63 | 1.74 |
| **lateraloccipital** | 1.38 | 0.93 | 1.18 | 1.23 | 1.44 | 1.53 |
| **isthmuscingulate** | 1.12 | 1.40 | 1.13 | 1.62 | 1.12 | 1.45 |
| **insula** | 1.19 | 1.33 | 1.17 | 1.64 | 2.00 | 1.23 |
| **inferiortemporal** | 1.42 | 1.13 | 1.09 | 0.89 | 1.37 | 1.19 |
| **inferiorparietal** | 1.15 | 1.51 | 1.11 | 1.13 | 1.50 | 1.07 |
| **fusiform** | 1.31 | 0.64 | 0.89 | 0.72 | 0.95 | 0.90 |
| **frontalpole** | 1.87 | 1.73 | 1.67 | 1.32 | 1.44 | 2.20 |
| **entorhinal** | 2.88 | 2.22 | 2.42 | 3.74 | 3.36 | 3.09 |
| **cuneus** | 1.25 | 1.90 | 1.55 | 3.26 | 1.47 | 1.55 |
| **caudalmiddlefrontal** | 1.95 | 1.66 | 1.98 | 2.16 | 2.02 | 1.37 |
| **caudalanteriorcingulate** | 1.49 | 1.23 | 3.76 | 1.85 | 1.69 | 1.40 |
| **bankssts** | 0.77 | 1.84 | 1.03 | 0.98 | 1.55 | 1.28 |

#### 3.2.4 Subcortical brain volume

The raw values (**Table S6**) and characteristics (**Fig. S9**) of the subcortical brain volumes were described in the **Supplementary Materials**. The CVs of the within-subject subcortex ranged from 0.64 to 6.17 for Sub1, 0.54 to 6.24 for Sub2, and 0.62 to 7.56 for Sub3 (**Table 6**). As depicted in the upper panel of **Fig. 5**, the subcortex including the thalamus, caudate, putamen, pallidum, hippocampus, and amygdala, showed very small CVs (< 2.5%) except the accumbens area (**Table 6**), especially the left side. In addition, **Table S7** depicted the CVs of the subcortices along with the cerebellum, ventricles, brain stem, corpus callosum (CC), and CSF, where it was shown that the corpus callosum showed much higher stability as a whole than its parts such as the anterior, middle, and posterior (**Fig. S11**).

**Fig. 5.**
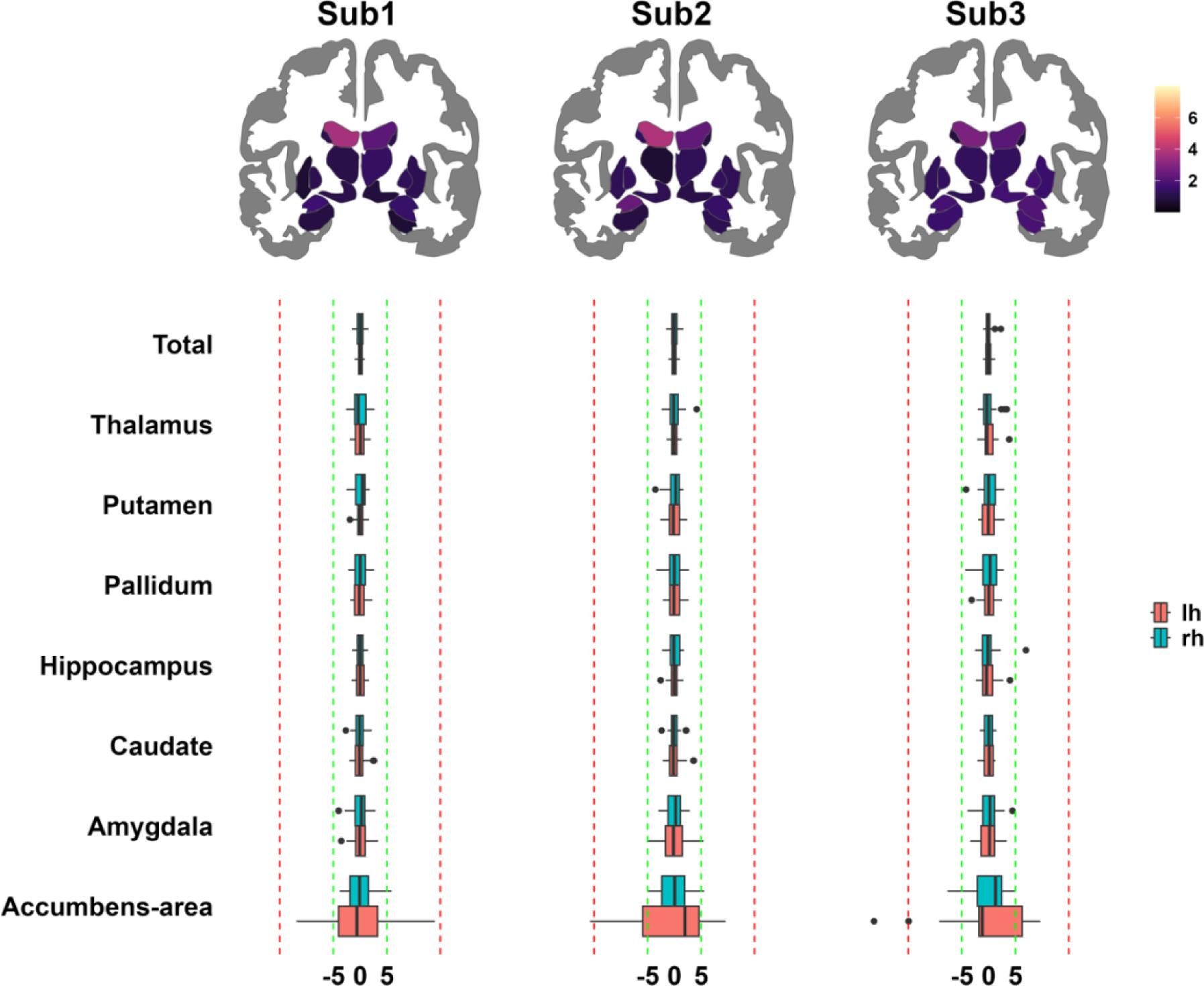
CVs and percentage change in brain volume. The upper panel depicts the coefficient of variation (CV) for each subject. The lower boxplots show the percentage change for each brain region and each subject. The green lines are in the range of ±5 % while the red lines depict the range of ±15%. All values in the boxplots are reported as percentages.

**Table 6.**
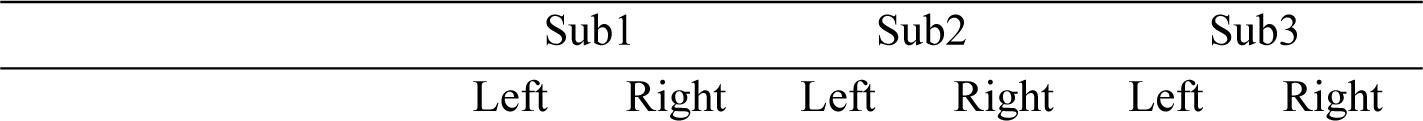
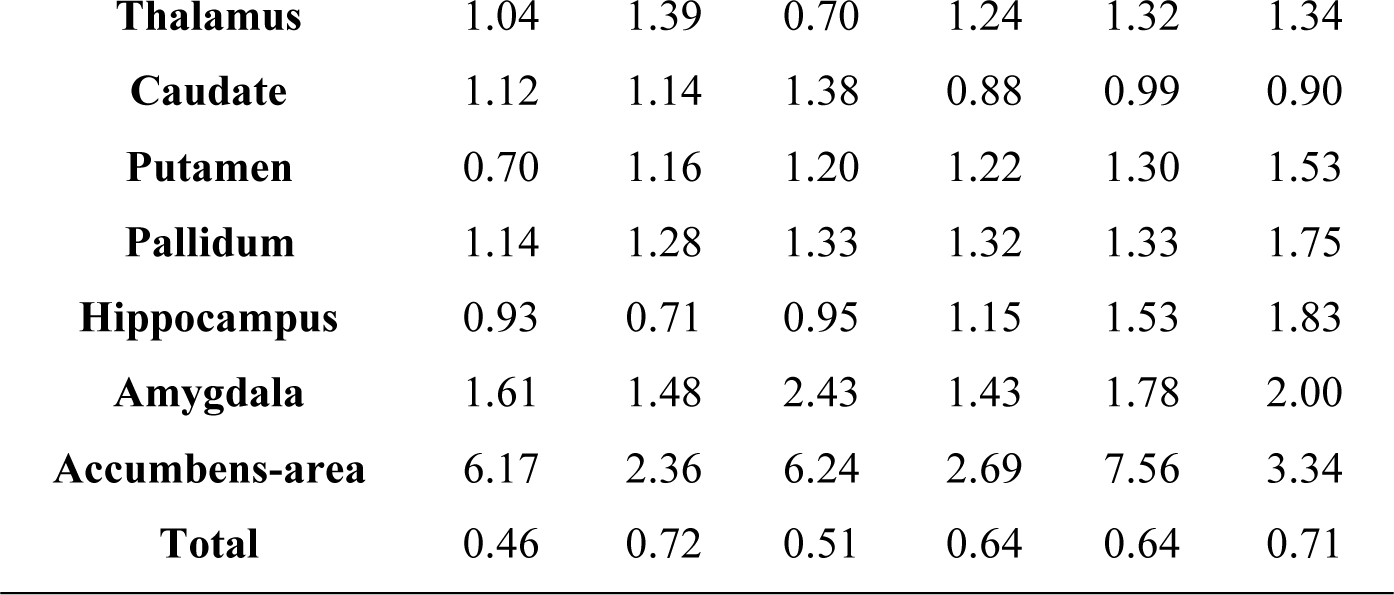
The CVs of the non-cortical brain volume based on the DK atlas.

#### 3.2.5 Correlation of percentage changes within and between different phenotypes

Within phenotypes, the average correlation coefficients of thickness (**Fig. 6A**), area (**Fig. 6B**), and volume (**Fig. 6C**) were 0.25, 0.23, and 0.27, respectively. Moreover, between phenotypes, the average correlation coefficients (without the diagonal lines) of thickness-volume (**Fig. 6D**), thickness-area (**Fig. 6E**), and area-volume (**Fig. 6F**) were 0.26, -0.09, and -3.4 × 10^-4^, respectively. More importantly, the correlation coefficients between brain regions (the diagonal lines in **Fig. 6DEF**) were positive for thickness-volume, negative for thickness-area, and negligible for area-volume (**Table 7**). The distribution of correlation coefficients of each phenotype association was depicted in **Fig. S13**.

**Fig. 6.**
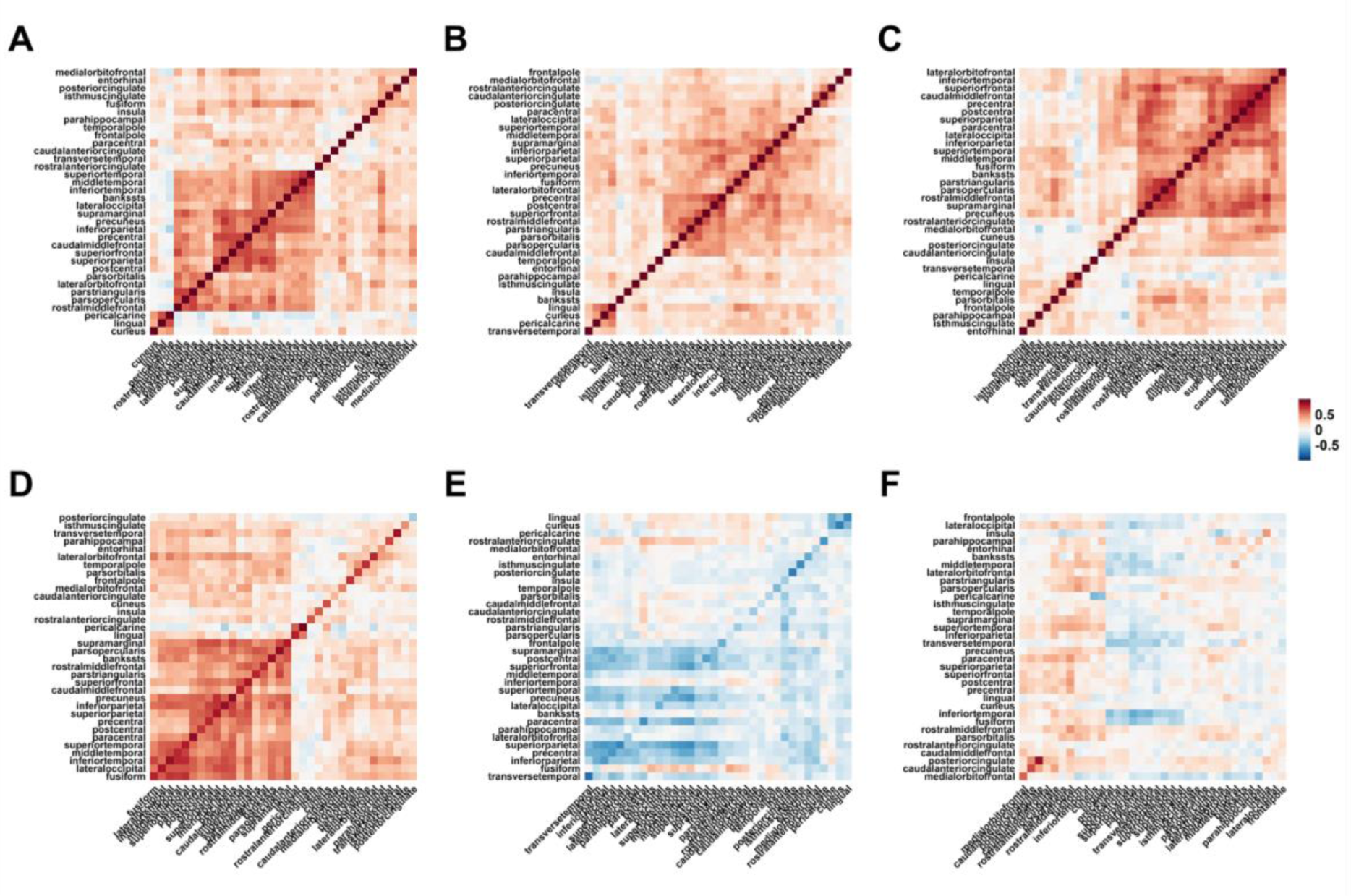
Correlation matrices within and between different phenotypes. Correlation matrices for **(A)** Thickness, **(B)** Surface area, **(C)** Brain volume, **(D)** Thickness and volume, **(E)** Thickness and area, and **(F)** Area and volume.

**Table 7.**
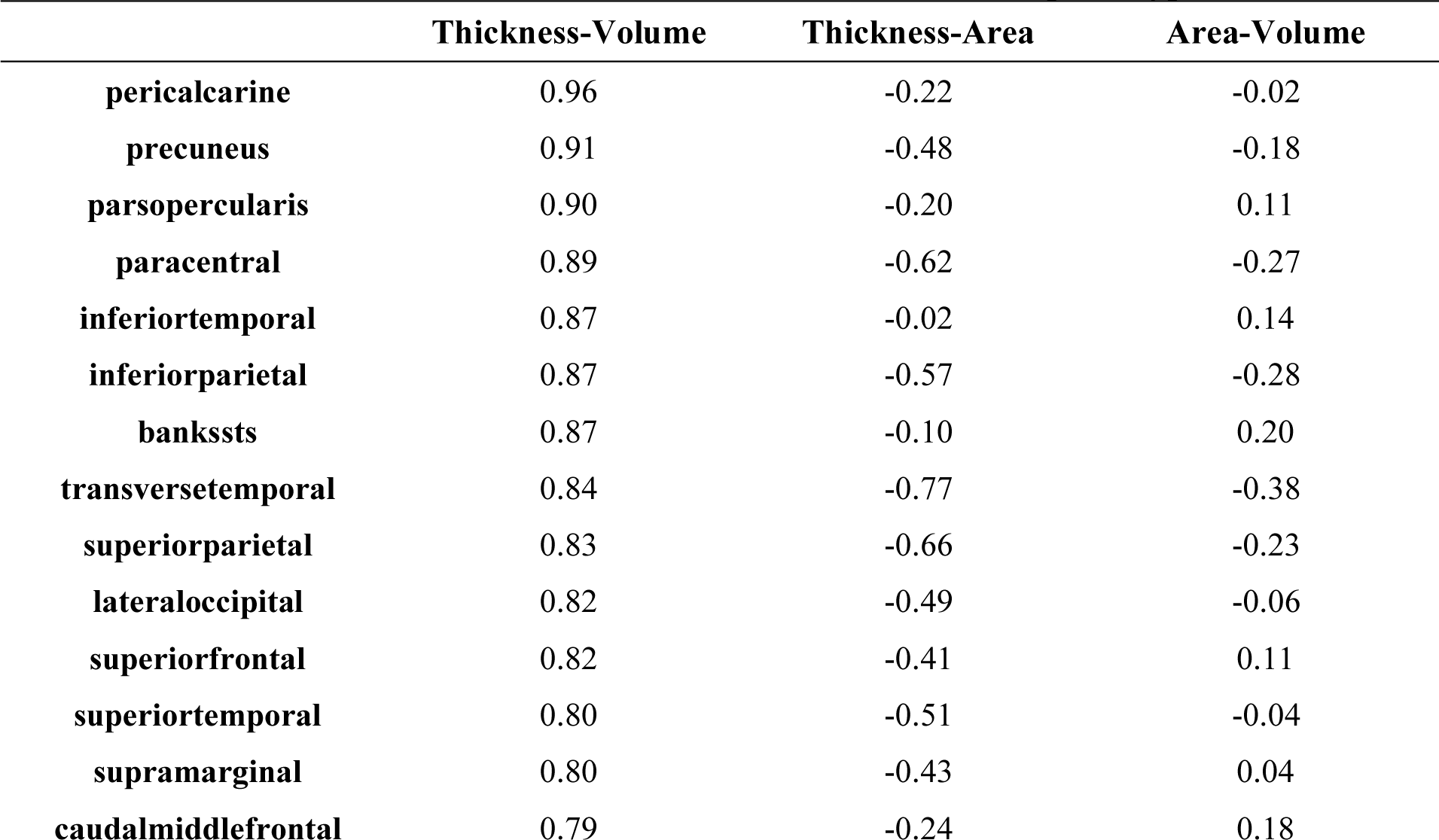
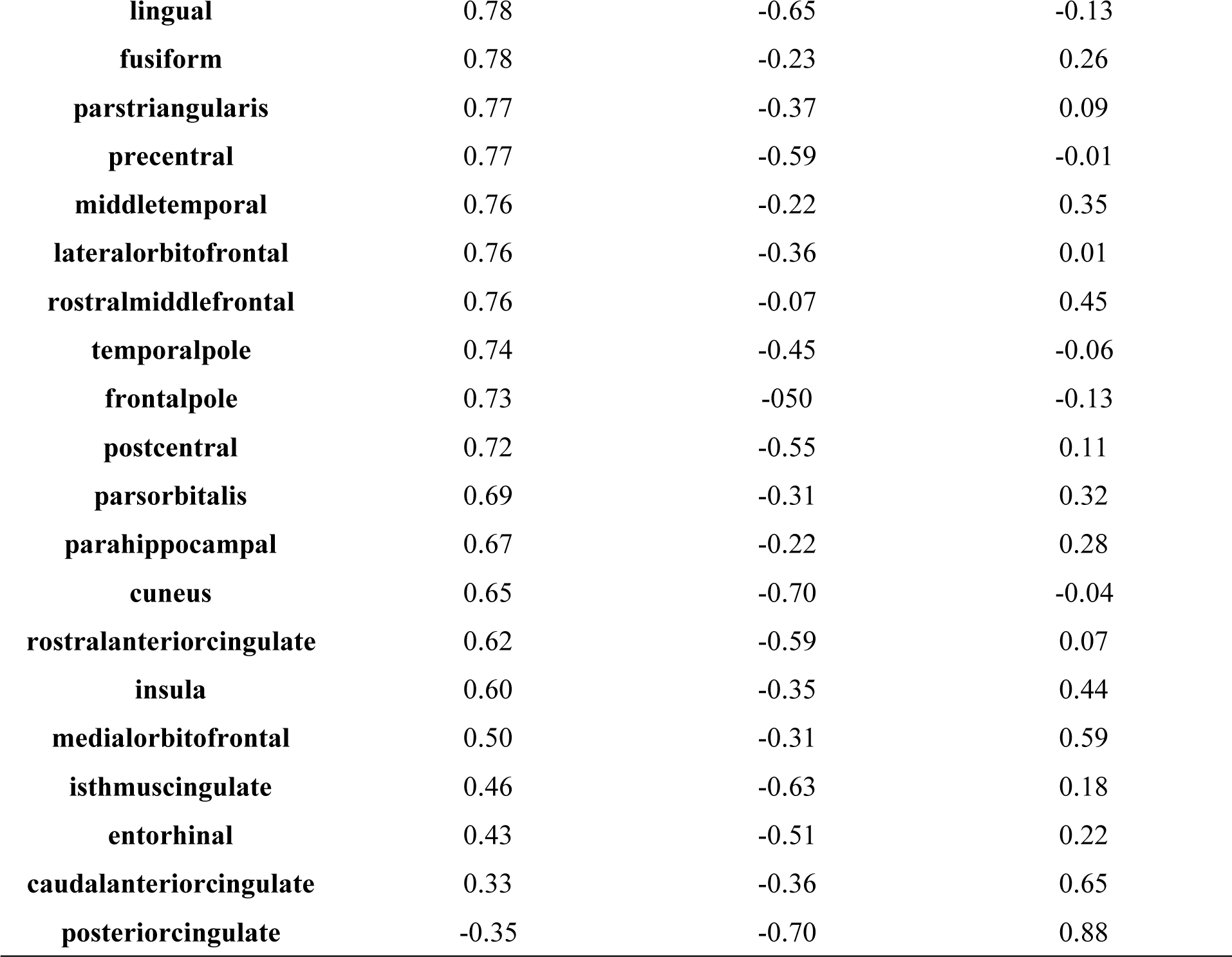
The correlation coefficients across different phenotypes.

|  | Thickness-Volume | Thickness-Area | Area-Volume |
| --- | --- | --- | --- |
| pericalcarine | 0.96 | -0.22 | -0.02 |
| precuneus | 0.91 | -0.48 | -0.18 |
| parsopercularis | 0.90 | -0.20 | 0.11 |
| paracentral | 0.89 | -0.62 | -0.27 |
| inferiortemporal | 0.87 | -0.02 | 0.14 |
| inferiorparietal | 0.87 | -0.57 | -0.28 |
| bankssts | 0.87 | -0.10 | 0.20 |
| transversetemporal | 0.84 | -0.77 | -0.38 |
| superiorparietal | 0.83 | -0.66 | -0.23 |
| lateraloccipital | 0.82 | -0.49 | -0.06 |
| superiorfrontal | 0.82 | -0.41 | 0.11 |
| superiortemporal | 0.80 | -0.51 | -0.04 |
| supramarginal | 0.80 | -0.43 | 0.04 |
| caudalmiddlefrontal | 0.79 | -0.24 | 0.18 |
| lingual | 0.78 | -0.65 | -0.13 |
| fusiform | 0.78 | -0.23 | 0.26 |
| parstriangularis | 0.77 | -0.37 | 0.09 |
| precentral | 0.77 | -0.59 | -0.01 |
| middletemporal | 0.76 | -0.22 | 0.35 |
| lateralorbitofrontal | 0.76 | -0.36 | 0.01 |
| rostralmiddlefrontal | 0.76 | -0.07 | 0.45 |
| temporalpole | 0.74 | -0.45 | -0.06 |
| frontalpole | 0.73 | -0.50 | -0.13 |
| postcentral | 0.72 | -0.55 | 0.11 |
| parsorbitalis | 0.69 | -0.31 | 0.32 |
| parahippocampal | 0.67 | -0.22 | 0.28 |
| cuneus | 0.65 | -0.70 | -0.04 |
| rostralanteriorcingulate | 0.62 | -0.59 | 0.07 |
| insula | 0.60 | -0.35 | 0.44 |
| medialorbitofrontal | 0.50 | -0.31 | 0.59 |
| isthmuscingulate | 0.46 | -0.63 | 0.18 |
| entorhinal | 0.43 | -0.51 | 0.22 |
| caudalanteriorcingulate | 0.33 | -0.36 | 0.65 |
| posteriorcingulate | -0.35 | -0.70 | 0.88 |

### 3.3 The influence of the stability

Regarding the time-of-day and daylight-hour effects, the meta-analysis can tell us about the significance characters of this specific dataset while the equivalent test can tell us about the generalization of our results by indicating whether the observed effect size is comparable to a predefined effect size (i.e., 0.5) (Lakens, 2017; Lakens et al., 2018).

The meta-analysis and equivalence test results of the time-of-day and daylight-hour effects were depicted in **Fig. 7**. The square dots represented the effect size estimated from this study where the extended lines illustrated the 90% (bold lines) and 95 % (narrow lines) confidence intervals (CI). If the CIs did not cross the zero lines, it means the meta-analysis results are significant. Moreover, if the CIs did not cross the ± 0.5 lines, it means the equivalence analysis results are significant meaning the results are trivial or can be practically ignored (Lakens, 2017; Lakens et al., 2018).

**Fig. 7.**
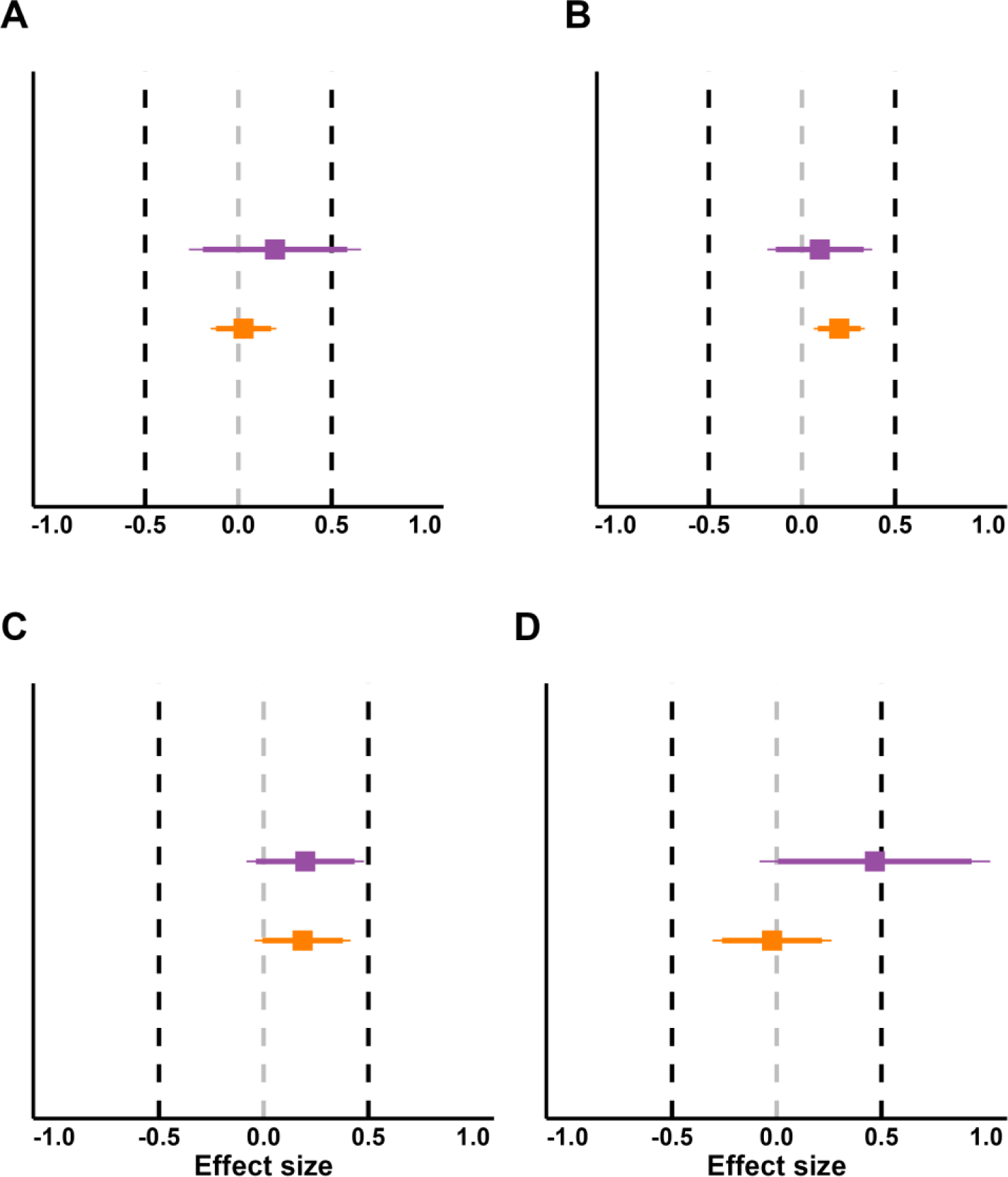
The meta-analysis and equivalence test results of the time-of-day and daylight-hour effects. **(A)** Average brain thickness. **(B)** Total surface area. **(C)** Total cortical brain volume. **(D)** Total subcortical brain volume (summary of the thalamus, pallidum, amygdala, hippocampus, putamen, accumbens area, and caudate nucleus). The square dots represented the effect size estimated from this study where the extended lines illustrated the 90% (bold lines) and 95 % (narrow lines) CI. Purple and yellow colors represent the time-of-day and daylight-hour effects, respectively.

#### 3.3.1 Time-of-day effect

The distributions of percentage change for morning and afternoon sessions of each brain region for each IDP were depicted in **Figs S3, S5, S7, S8**, and **S10.**

*Thickness (average)*: the significant time-of-day (ToD) effect can be observed in sub1 (*β* = 0.63*, p* = 0.006*, SE* = 0.22), but not in sub2 (*β* = 0.001*, p* = 0.996*, SE* = 0.23), and sub3 (*β* = - 0.11*, p* = 0.72*, SE* = 0.31). Collectively, meta-analysis suggested that there was no significant ToD effect (*β* = 0.196*, p* = 0.40*, SE* = 0.24, CI 95% = [-0.26 0.66]). However, we cannot rule out there was no ToD effect considering medium effect size (d = 0.5) based on the equivalence test (*Z* = -1.29, *p* = 0.099).

*Surface area (total)*: no ToD effects were shown in sub1 (*β* = 0.04*, p* = 0.87*, SE* = 0.23), sub2 (*β* = -0.06*, p* = 0.80*, SE* = 0.23), and sub3 (*β* = 0.45*, p* = 0.14*, SE* = 0.30). Jointly, the meta-analysis (*β* = 0.096*, p* = 0.51*, SE* = 0.14, CI 95% = [-0.19 0.38]) and equivalence test (*Z* = - 2.82, *p* = 2.43 × 10^-3^) also illustrated that there was no ToD effect.

*Cortical volume (total)*: nonsignificant ToD effects were observed for sub1 (*β* = 0.39*, p* = 0.09*, SE* = 0.23), sub2 (*β* = 0.06*, p* = 0.80*, SE* = 0.23), and sub3 (*β* = 0.11*, p* = 0.71*, SE* = 0.31). Furthermore, the meta-analysis (*β* = 0.199*, p* = 0.17*, SE* = 0.14, CI 95% = [-0.08 0.48]) and the equivalence test (*Z* = -2.10, *p* = 0.02) corroborated the none ToD effect.

Subcortical volume(total): sub2 showed significant ToD effect (*β* = 0.93*, p* = 0.003, *SE* = 0.29), but not for sub1 (*β* = 0.09*, p* = 0.79*, SE* = 0.33) and sub3 (*β* = 0.28*, p* = 0.52*, SE* = 0.43). Although the meta-analysis demonstrated that there was no ToD effect (*β* = 0.47*, p* = 0.10*, SE* = 0.28, CI 95% = [-0.08 0.48]), the equivalence test suggested that the ToD effect could not be ruled out (*Z* = -0.11, *p* = 0.46).

#### 3.3.2 Daylight-hour effect

The distributions of percentage change along the daylight hour of each phenotype were depicted in **Figs S2D, S4D,** and **S6D.**

*Thickness (average)*: no significant effects were found in sub1 (*β* = -0.003*, p* = 0.77*, SE* = 0.12), sub2 (*β* = -0.007*, p* = 0.53*, SE* = 0.11), and sub3 (*β* = 0.25*, p* = 0.10*, SE* = 0.14). The meta-analysis (*β* = 0.03*, p* = 0.82*, SE* = 0.09, CI 95% = [-0.15 0.20]) and the equivalence test (*Z* = -5.26, *p* = 7.39 × 10^-8^) reconfirmed the non-significant effect.

*Surface area (total):* a significant effect was found in sub3 (*β* = 0.30*, p* = 0.04*, SE* = 0.14), but not in sub1 (*β* = 0.15*, p* = 0.21*, SE* = 0.11), and sub2 (*β* = 0.19*, p* = 0.097*, SE* = 0.11). The meta-analysis demonstrated that there was a significant effect (*β* = 0.20*, p* = 0.004*, SE* = 0.07, CI 95% = 0.06 0.34]), however, the equivalence test showed that the effect could be ignored since it was too small (*Z* = -4.31, *p* = 8.31 × 10^-6^)

*Cortical volume (total)*: the significant effect was found in sub3 (*β* = 0.44*, p* = 0.002*, SE* = 0.13) but not for sub1 (*β* = 0.12*, p* = 0.30, *SE* = 0.12), for sub2 (*β* = 0.003*, p* = 0.77*, SE* = 0.11). The meta-analysis (*β* = 0.19*, p* = 0.11*, SE* = 0.12, CI 95% = [-0.04 0.42]) and the equivalence test (*Z* = -2.69, p = 3.62 × 10^-3^) suggested that the effect was nonsignificant and trivial compared to effect size 0.5.

*Subcortical volume (total)*: no significant effects were in sub1 (*β* = -0.04*, p* = 0.80*, SE* = 0.17), sub2 (*β* = -0.24*, p* = 0.13*, SE* = 0.16), and sub3 (*β* = 0.28*, p* = 0.18*, SE* = 0.20). The meta-analysis demonstrated a non-significant effect (*β* = -0.02*, p* = 0.88*, SE* = 0.15, CI 95% = [-0.31 0.26]), and the daylight-hour effect was practically trivial based on the equivalence test result (*Z* = 3.29, *p* = 4.96 × 10^-4^).

#### 3.3.3 Head movement effect

In sub3, two sessions with excessive head motion indicated by the CJV (sessions 1 and 7) were excluded in the previous analyses. To investigate the head movement effect, CVs were recalculated when considering the two excluded sessions, which were then compared to the previously reported CVs. Paired t-tests were then used to compare those two CVs.

The CVs (**Table 8**) and percentage changes (**Fig. 8**) were significantly improved after excluding those two sessions for the cortical thickness (left hemisphere: *t_33_* = -7.6, *p* = 4.8 × 10^-9^, *Cohen’s d* = 1.3; right hemisphere: *t_33_* = -7.13, *p* = 1.8 × 10^-8^, *Cohen’s d* = 1.22), surface area (left: *t_33_* = -6.03, *p* = 4.4 × 10^-7^, *Cohen’s d* = 1.03; right: *t_33_* = -2.27, *p* = 7.9 × 10^-5^, *Cohen’s d* = 0.73), and cortical brain volumes (left: *t_33_* = -5.06, *p* = 7.6 × 10^-6^, *Cohen’s d* = 0.87; right: *t_33_* = -6.21, *p* = 2.6 × 10^-7^, *Cohen’s d* = 1.06).

**Fig. 8.**
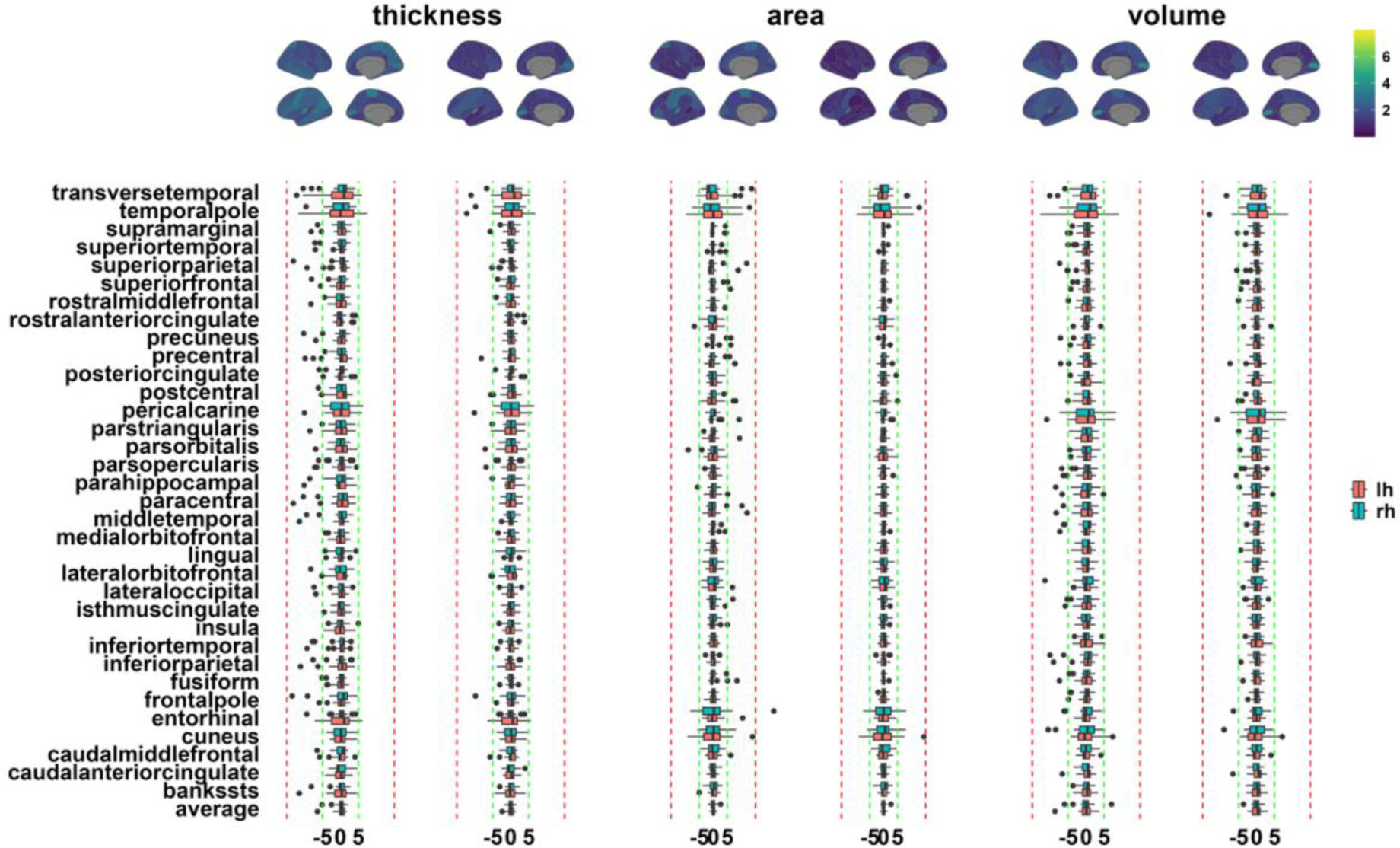
The CVs and percentage changes before and after excluding two sessions with excessive head movement in sub3. The upper panels depicted the CVs for cortical thickness, surface area, and cortical brain volume. The bottom panels illustrate the percentage changes in each brain region for cortical thickness, surface area, and cortical brain volume. Under each IDP, the left panels were results from all sessions whereas the right ones were results from sessions excluding the two sessions.

**Table 8.**
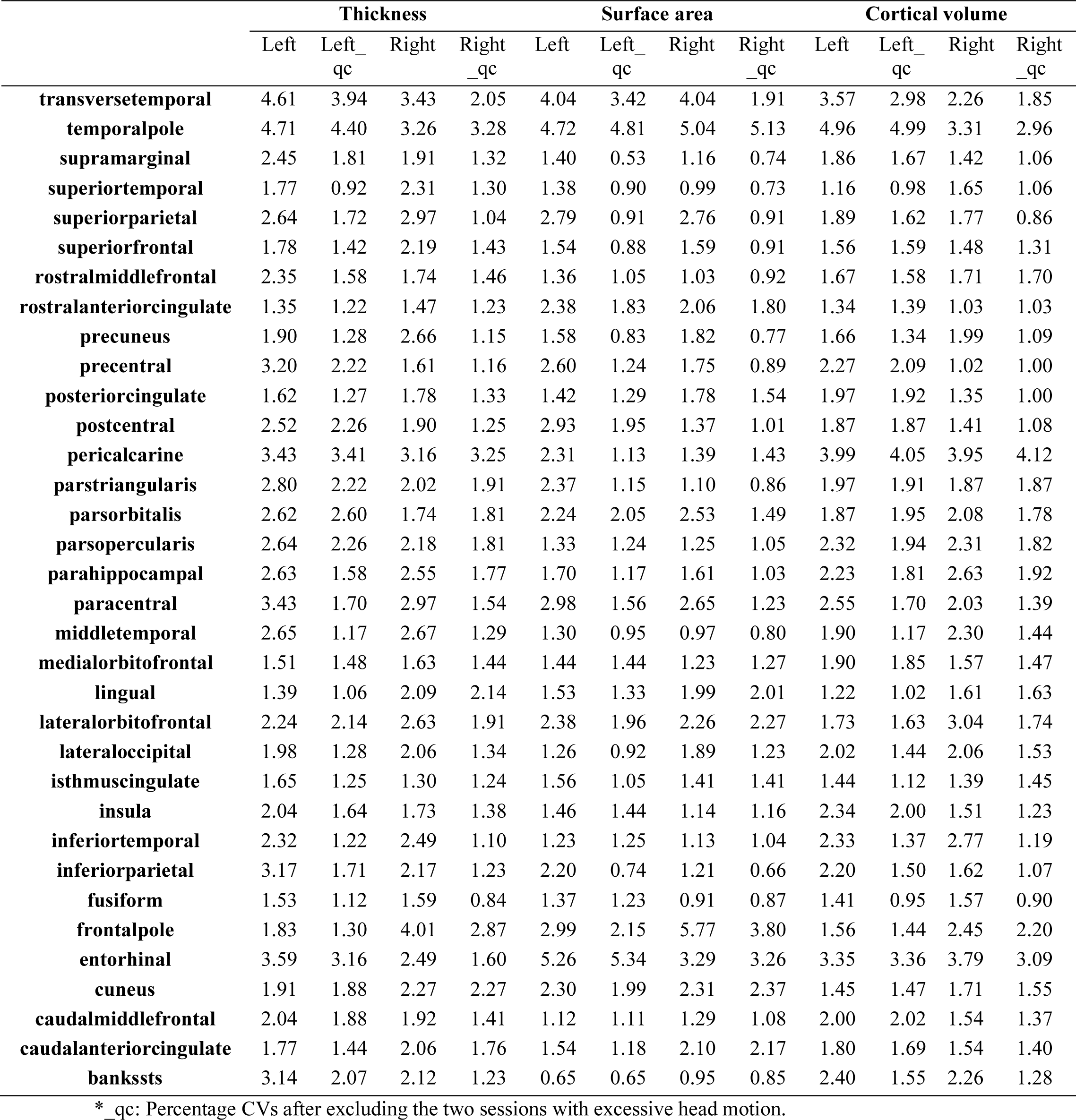
The CVs of the cortical thickness based on the DK atlas.

#### 3.3.4 Brain region size

As shown in **Fig. 9**, all IDPs CVs were significantly negatively correlated with their brain region size: thickness (*r* = -0.48, *t_202_* = -7.84, *p* = 2.62 × 10^-13^, CI 95% = [-0.58 -0.37]); surface area (*r* = -0.45, *t_202_* = -7.26, *p* = 8.13 × 10^-12^, CI 95% = [-0.56 -0.34]); cortical volume (*r* = -0.37, *t_202_* = -5.68, *p* = 4.71 × 10^-8^, CI 95% = [-0.48 -0.25]); subcortical volume (*r* = - 0.52, *t_40_* = -3.83, *p* = 4.40 × 10^-4^, CI 95% = [-0.71 -0.25]).

**Fig. 9.**
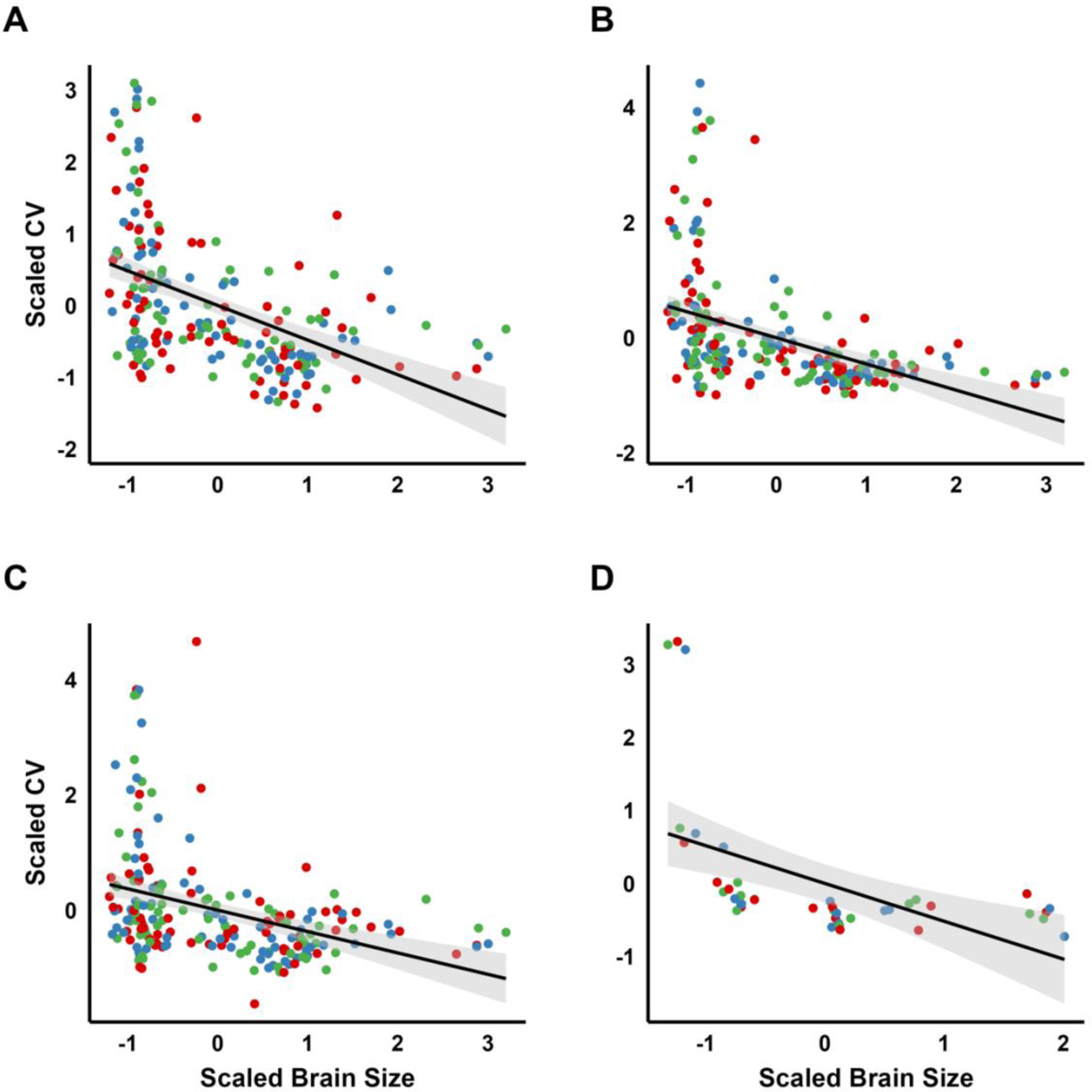
The brain region size negatively correlated with CVs. The x-axis represents the scaled brain region size for the 68 (left and right) brain regions. The y-axis depicts the scaled CVs for the 68 (left and right) brain regions. The red color represents subject 1, the blue depicts subject 2, and the green represents subject 3.

## 4 Discussion

This study examined the within-subject stability and the influence factors of the IDPs related to cortical thickness, surface area, and brain volume. Our findings demonstrated that the longitudinal stream of FreeSurfer produced more stable results compared to the cross-sectional stream. The stability of the IDPs was high, with most coefficients of variation (CVs) and percentage changes falling within 2% and 5%, respectively. Notably, CVs were negatively correlated with brain region size. Furthermore, percentage changes in cortical thickness were strongly correlated with cortical volume and negatively correlated with surface area. The effects of ToD and daylight hours on these measures were complex and will be further elaborated. Additionally, apparent head motion resulted in the underestimation of cortical thickness and volume and the overestimation of surface area. The broader implications of these findings, along with the specific characteristics of each IDP, are discussed in the following sections.

The longitudinal stream generates more reliable within-subject results, as suggested by much smaller CVs (**Table 2**) and smaller percentage changes (**Fig. 1**) compared to the cross-sectional stream. The longitudinal stream was proposed in 2012 by the FreeSurfer team, and since then it has been recommended for analyzing longitudinal datasets (Reuter et al., 2012). Specifically, the longitudinal stream creates and utilizes an unbiased within-subject template to generate a robust, inverse consistent registration, which boosts the stability and statistical power (Reuter et al., 2012; Vidal-Piñeiro et al., 2024). Indeed, our results are well in line with this argument. Accordingly, we also advocate the longitudinal stream should be used when analyzing longitudinal datasets.

### 4.1 The characteristics of IDPs

Cortical thickness is one of the IDPs of greatest interest in human brain research, given its role in defining normal cortical maturation (Bethlehem et al., 2022) and alterations observed in various neurological and mental disorders (Bethlehem et al., 2022; Frisoni, Fox, Jack, Scheltens, & Thompson, 2010; Lemaitre et al., 2012; Thompson et al., 2020). In our dataset, cortical thickness ranged from 1.50 to 4.03 mm (**Table S2**) which falls well within the known bounds of 1 and 4.5 mm (Fischl & Dale, 2000) indicating the validity of our dataset. What’s more, we observed that the thickest brain regions possess the largest standard deviations which agrees well with previous studies (Fischl & Dale, 2000; Hutton, De Vita, Ashburner, Deichmann, & Turner, 2008). Previous studies have also shown that the intersubjective standard deviations can be around 0.5 mm (Fischl & Dale, 2000; Hutton et al., 2008). Complementing that, we found the within-subject standard deviations are much smaller, where most of them lie well below 0.05 mm with the thickest brain regions possessing the largest standard deviation of around 0.10 mm (**Table S2**). These results indicate that the cortical thickness measurement is very stable across a short time (about a year) in adults.

The surface area and brain volumes are the other two IDPs that can be constructed from the T1w images which can be used as a proxy index of the brain size (Genon et al., 2022). As expected, we saw large differences between subjects regarding brain sizes (**Table S3-S6**) where sub2 has the largest brain size followed by sub3 and sub1. Even though the brain size varies, based on the DK atlas, the biggest and smallest surface areas (**Fig. S4B**) and brain volumes (**Figs. S6B and S9**) are the same within the three subjects. The DK atlas was developed based on curvature-based information such as the sulcal representations and the anatomic curvature (Desikan et al., 2006), therefore, the different size of the surface area and brain volume in different brain regions represent their own territories constrained by the curvature boundaries. The impact of the brain region size on stability was discussed in the next section.

### 4.2 IDPs are stable across one year

We found that the IDPs including the cortical thickness, surface area, and brain volumes in most brain regions are stable, where CVs are well constrained at 2% and the percentage change values are well within 5% over one year. CV and percentage change values have been used to evaluate the stability given that they are unitless and suitable for comparing variables with different sizes (Borga et al., 2020; Carbonell et al., 2022; M. Y. Wang et al., 2024; Y. Wang et al., 2021). CV is the division between standard deviation and average, where the smaller value denotes a much more stable measurement. In the same vein, percentage change is the division between the discrepancy and the average, where smaller values denote high stability and less variation. Generally, a measurement is considered reliable or stable when CVs are within 5 % or percentage changes are within 10% (Borga et al., 2020; Carbonell et al., 2022; M. Y. Wang et al., 2024; Y. Wang et al., 2021). For example, using CVs, it is found that the stability of brain metabolites such as N-acetyl-aspartate was reasonably high with CVs around 4% (M. Y. Wang et al., 2024). Therefore, the 2% of CVs in this study indicate that IDPs are fairly stable. In addition, the percentage change of 5% could encompass the annual 0.5 to 1 % decrease, for example in cortical brain volume, during adulthood (Lemaitre et al., 2012; Sele, Liem, Merillat, & Jancke, 2021; Storsve et al., 2014).

We further found that CVs were negatively correlated with brain region size (**Fig. 9**). For example, smaller brain regions such as the temporal pole, frontal pole, pericalcarine, and entorhinal cortex manifested larger fluctuations across all three IDPs. This result agrees well with the previous study, where the longitudinal (2-year interval) brain change stability showed a similar pattern (Parsons et al., 2024). In addition, small subcortical brain regions such as the accumbens area showed very high variations. Therefore, we infer that the greater variation in IDPs in these regions is likely due to their smaller size (**Fig. 9**) or proximity to air cavities, making them more prone to errors and more difficult to parcellate. Another speculation could be that these brain regions do vary that much, however, the underlying biological reason for it needs to be further investigated. Future studies are warranted to explore this issue. To be noticed, only subject 2 manifested high variability in the cingulate cortex especially the PCC compared to the other two participants indicating that individual differences can also influence the temporal stability of IDPs.

### 4.3 Cortical thickness changes together with cortical volume but not with surface area

We found that the overall association within each phenotype was quite similar (**Fig. 6A-C**) with an average strength of around 0.25. On the contrary, the relationships between each phenotype are quite divergent (**Fig. 6D-F**). Specifically, the percentage changes in cortical thickness and volume manifested a strong positive correlation while both of them illustrated a weak correlation with the percentage change of surface area. More importantly, the percentage changes in the cortical thickness showed a negative correlation with that of the surface area. These short-timescale results corroborate well with the long-timescale normal aging study (Storsve et al., 2014). Collectively, these results reinforce that cortical thickness and surface area play distinct roles in brain anatomy and aging (Lemaitre et al., 2012; Storsve et al., 2014; Vijayakumar et al., 2016), which is sensible since they have distinct genetic roots (Panizzon et al., 2009).

### 4.4 The time-of-day and daylight-hour effects

We found no time-of-day effect (meta-analysis results) on average cortical thickness, total surface area, and total brain volumes (**Fig. 7**). Previous studies, however, have reported significantly thinner cortical thickness (Trefler et al., 2016) and smaller brain volume (Karch et al., 2019; Nakamura et al., 2015; Trefler et al., 2016) in the afternoon compared to the morning. In this study, IDPs did show smaller values in the afternoon sessions but were not significantly different from the morning sessions. The discrepancy between our results and others could result from the following factors. First, this study is a within-subject design while two of the other studies were between-subject design (Nakamura et al., 2015; Trefler et al., 2016), from which the intersubjective variation could obstruct the reported time-of-day effect. Second, this study has recorded at least 10 sessions and up to 23 sessions for each subject in the morning or afternoon sessions whereas the other studies either had one data point (Nakamura et al., 2015) or two data points for morning and afternoon sessions in their primary and secondary datasets (Trefler et al., 2016). Fourth, although one is a within-subject design (Karch et al., 2019), the longitudinal stream, which will provide more robust results as also evidenced by this study and others (Reuter et al., 2012), was not implemented.

Although no time-of-day effects were uncovered in our study, we acknowledge that we could not rule out (equivalent test results) that there is no true time-of-day effect on average cortical thickness and total subcortical brain volume given the effect size of 0.5 (**Fig. 7)**. The results were largely driven by subject 1 on the average cortical thickness and by subject 2 on the total subcortical brain volume manifesting the inter-individual characteristics, from which studies with more subjects with within-subject design are needed to reconcile this effect. But before that, we recommend that when estimating the abovementioned two IDPs, the time-of-day effect should be considered since their effect sizes were not too trivial to be ignored. However, the time-of-day effect on the total cortical brain volume and total surface area was subtle with which the factor could be ignored.

Similarly, we found no daylight-hour effect (meta-analysis) on average cortical thickness, total cortical as well as subcortical brain volumes, but not total surface area. Previous studies have focused on the seasonal effect (Zhang et al., 2023) on functional brain organization and stated that daylight could reconfigure the resting state brain networks (M. Y. Wang et al., 2023b). However, to the best of our knowledge, this study could be the first study directly investigating the daylight effect on T1w-derived phenotypes. Although there is a daylight-hour effect on the total surface area, the effect size is too small to be considered significant. Taken together, we could suggest that the daylight hours could be left out when estimating the IDPs.

Regarding the equivalent test to the time-of-day and daylight-hour effects, to be noticed, deciding whether the effect size is too trivial is heavily dependent on the predefined effect size (Lakens, 2017; Lakens et al., 2018). In this study, an effect size of 0.5 was used, one could argue that the effect size could be set at 0.3 or 0.8 depending on the objectives of one’s studies (Lakens, 2017; Lakens et al., 2018).

### 4.5 Head motion underestimates thickness and volume whereas it overestimates surface area

Head motion can be indicated by the CJV values, with higher CJV values denoting apparent head motion, which could deteriorate the SNR and CNR, and blur the image (Ganzetti et al., 2016). After excluding the two sessions with excessive head motion in subject 3, the stability of all the T1w-derived phenotypes has been significantly improved evidenced by smaller CVs (**Table 8**) and percentage changes (**Fig. 8**). However, the pattern is different. On tne hand, the cortical thickness and cortical volume estimation were increased indicating the underestimation induced by head motion, which is in concordance with the previous study (Reuter et al., 2015). On the other hand, the surface area measurement is decreased indicating that head motion causes the surface area to be underestimated. This result indicates that the head motion evaluation should be controlled especially when investigating participants who move a lot such as autistic children or Parkinson’s patients (Reuter et al., 2015).

### 4.6 Limitations

Several limitations should be articulated before any conclusions. First, lacking female subjects hampers the generalization of our results. It is known that major events such as pregnancy (Hoekzema et al., 2017) and hormone levels can alter brain structure (Rizor et al., 2024). Second, although using the same MRI scanner and the same data processing stream can alleviate the interference from heterogeneous factors, it could impede the application to other datasets that were collected with other MRI scanners or data processing streams. Third, the timing of data collection was not fixed for each subject, for example, sub1 was not scanned at 10 AM on Mondays, instead, each subject was scanned based on the availability of the scanning time and their available time. Lastly, we acknowledge that the results are a mixture of the measurement (T1w) and the measure (MRI machine). Although the results indicate the IDPs are stable, it can be informative if the two sources could be separated, for example, using a phantom to evaluate the stability of the MRI machine and deduct from the current results which could lead to the measurement (T1w) stability.

## 5. Conclusion

In summary, the stability of T1w-derived phenotypes across one year during adulthood is fairly high with CVs within 2% while percentage changes within 5%. However, several small brain regions including the temporal pole, frontal pole, pericalcarine, entorhinal cortex, and accumbens area did manifest larger fluctuations. Moreover, time of day might be a confound when evaluating the average cortical thickness and total subcortical brain volumes. Furthermore, daylight length could be left out when evaluating the IDPs. Lastly, head motion leads to the underestimation of the cortical thickness and volume while overestimation of the surface area.

## Data and Code Availability

Data and code used in this manuscript can be found here on GitHub https://github.com/MengYunWang/BBSC/tree/main/fMRI/T1_analysis.

## Author Contributions

Meng-Yun Wang: Conceptualization, Data collection & curation, Formal analysis, Investigation, Methodology, Project administration, Visualization, Writing—original draft, and Writing—review & editing. Max Korbmacher: Conceptualization, Data collection, and Writing—review & editing. Rune Eikeland: Conceptualization, Data collection, Writing— review & editing. Stener Nerland: Methodology, Writing—review & editing. Didac Vidal-Pineiro: Methodology, Writing—review & editing. Karsten Specht: Conceptualization, Writing—review & editing, Funding acquisition.

## Ethics Statement

All participants have understood and signed written informed consent. The project was conducted according to the principles expressed in the Declaration of Helsinki. Ethics approval was granted by the Regional Committees for Medical and Health Research Ethics (REK-vest).

## Acknowledgments

We appreciate the technical support and data collection support from our radiologists (Christel Jansen, Eva Øksnes, Roger Barndon, Trond Øvreaas, Tor Fjørtoft, and Turid Randa) at the Haukeland University Hospital.

---

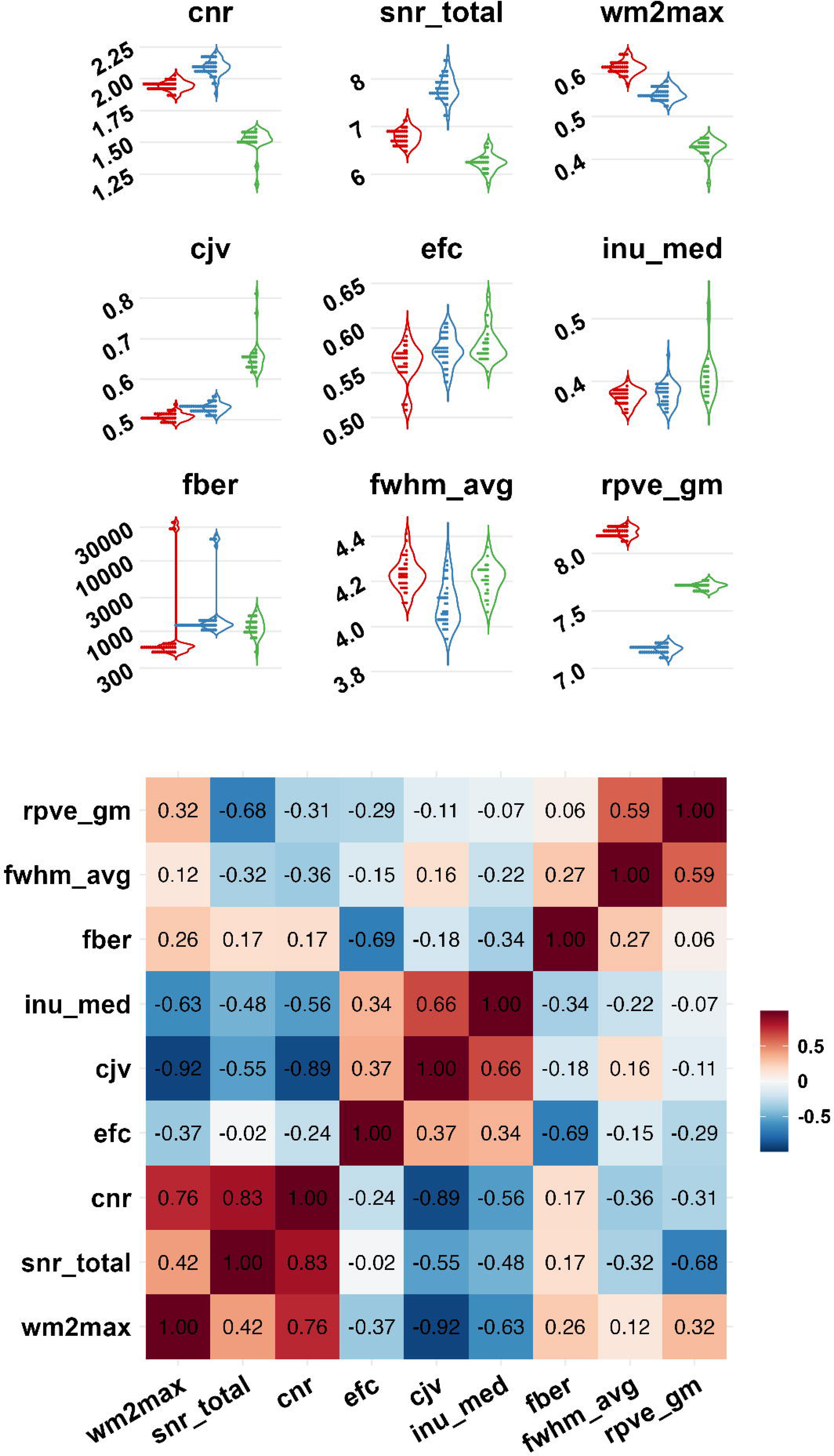

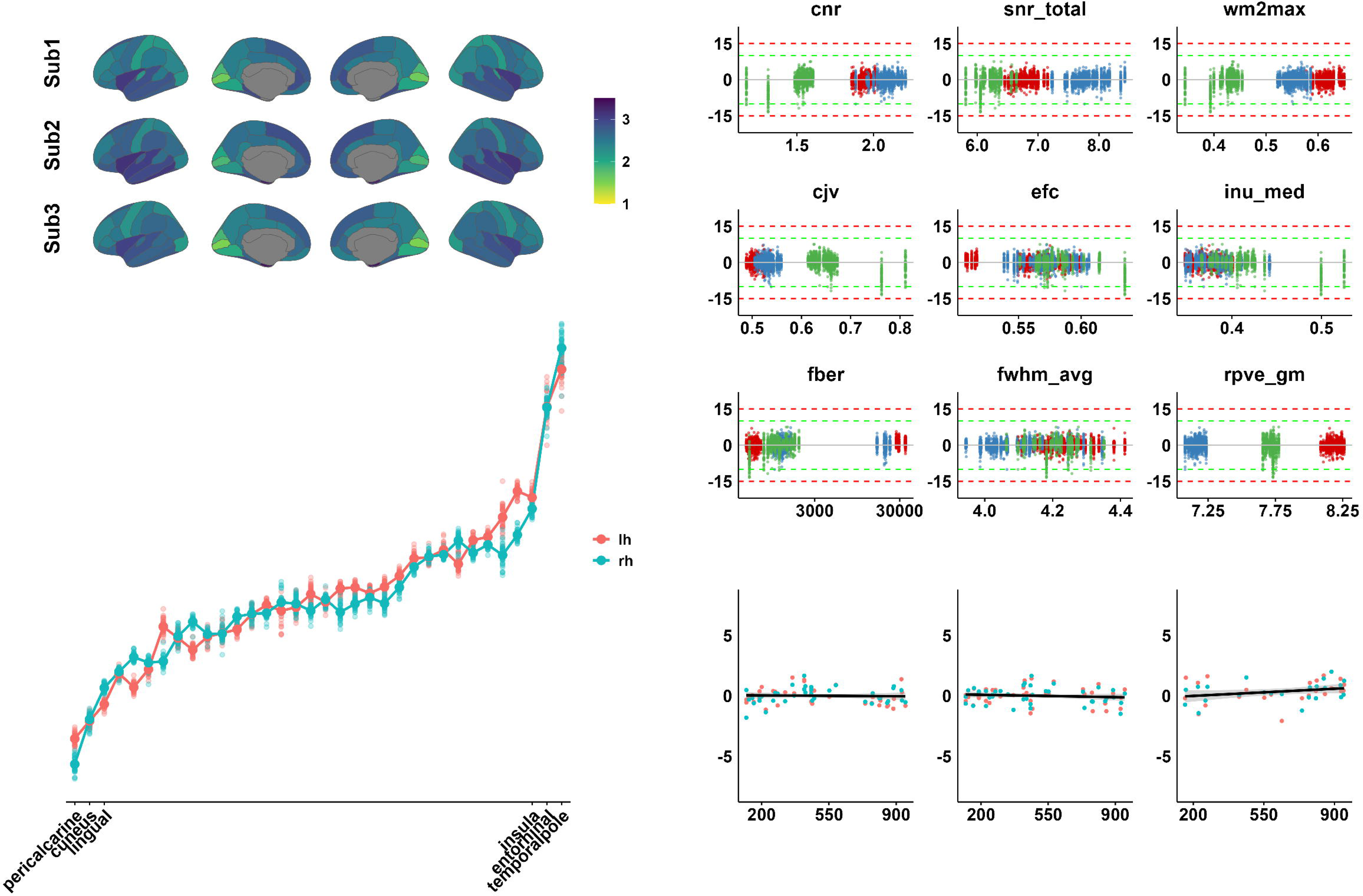

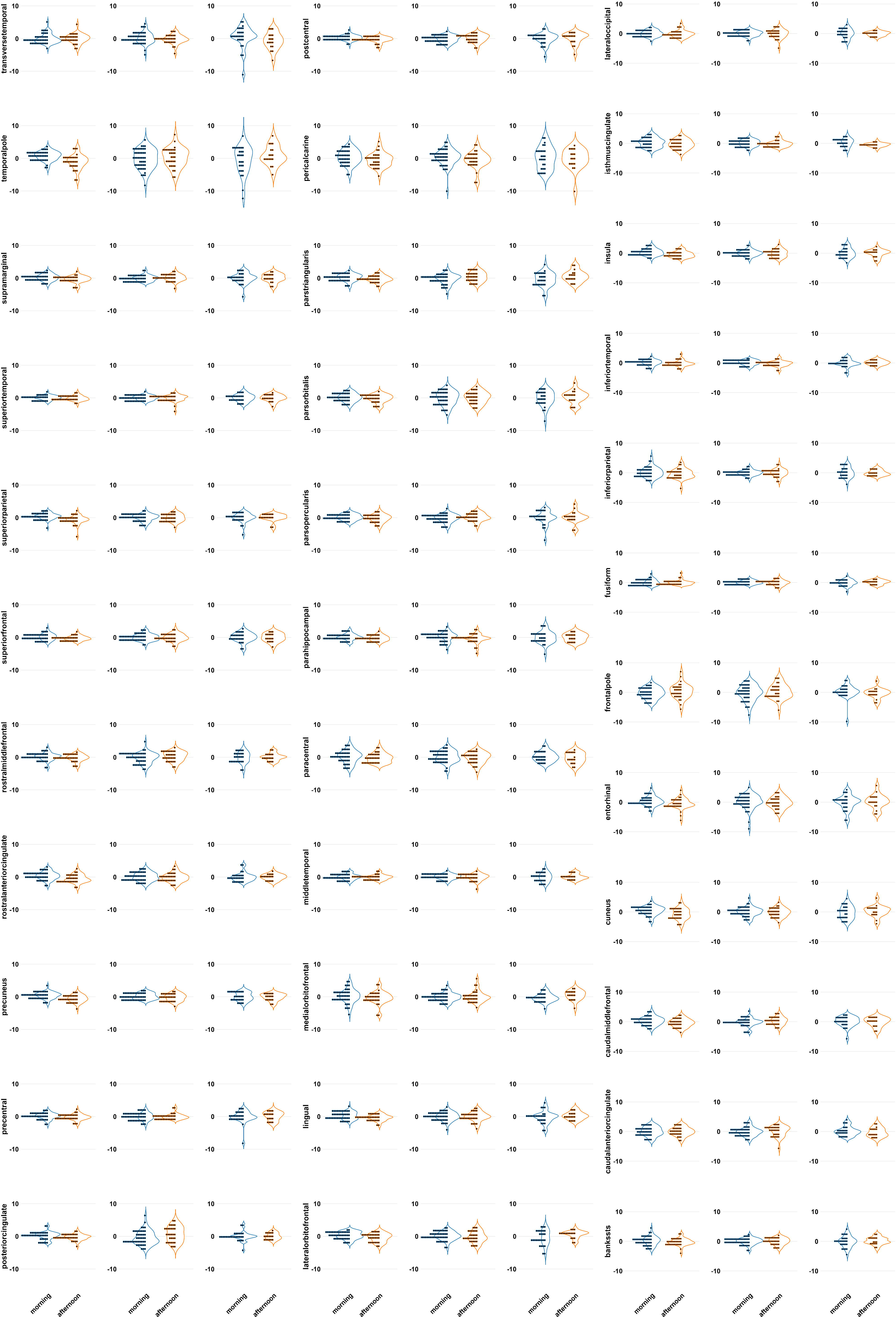

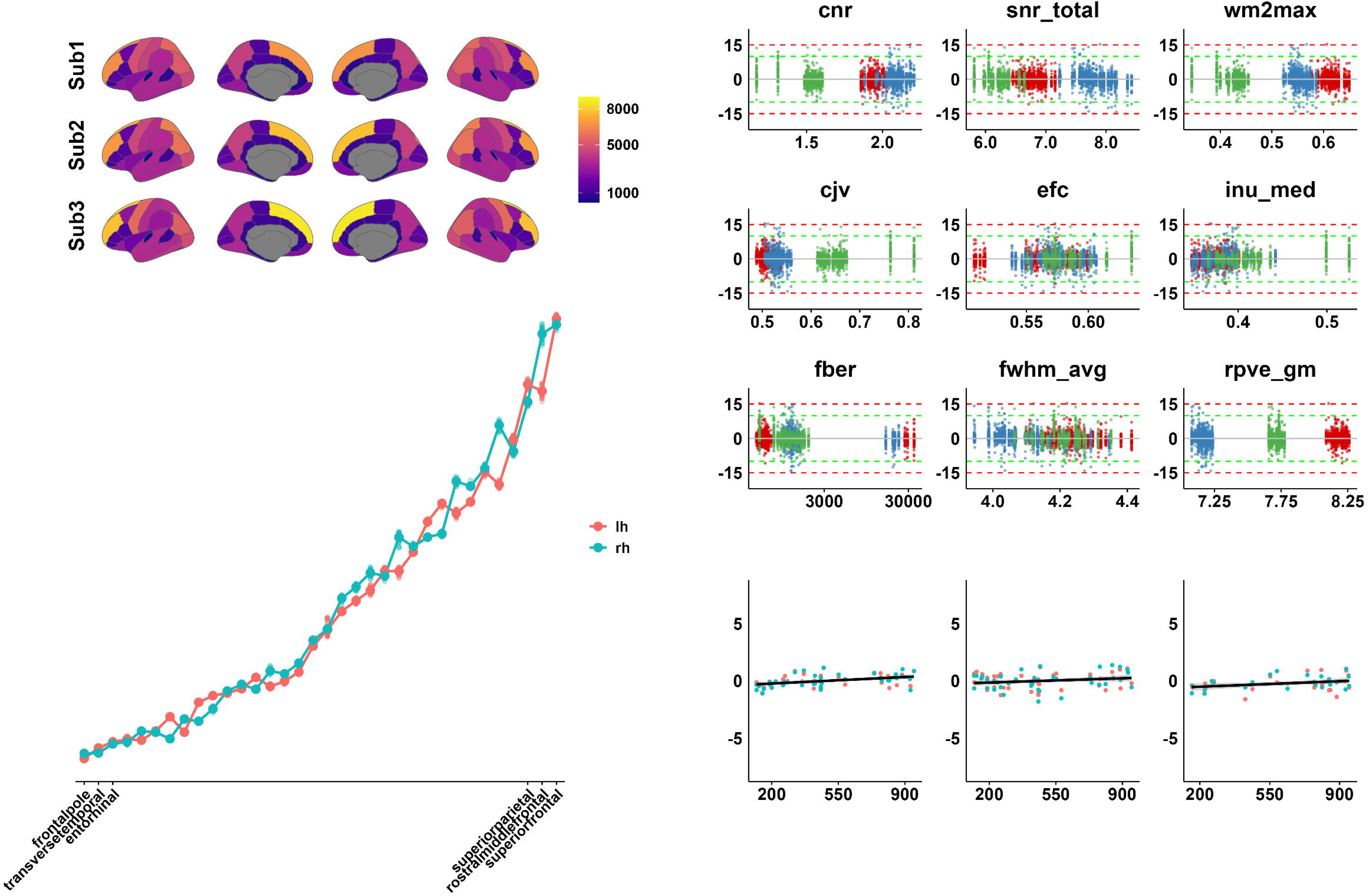

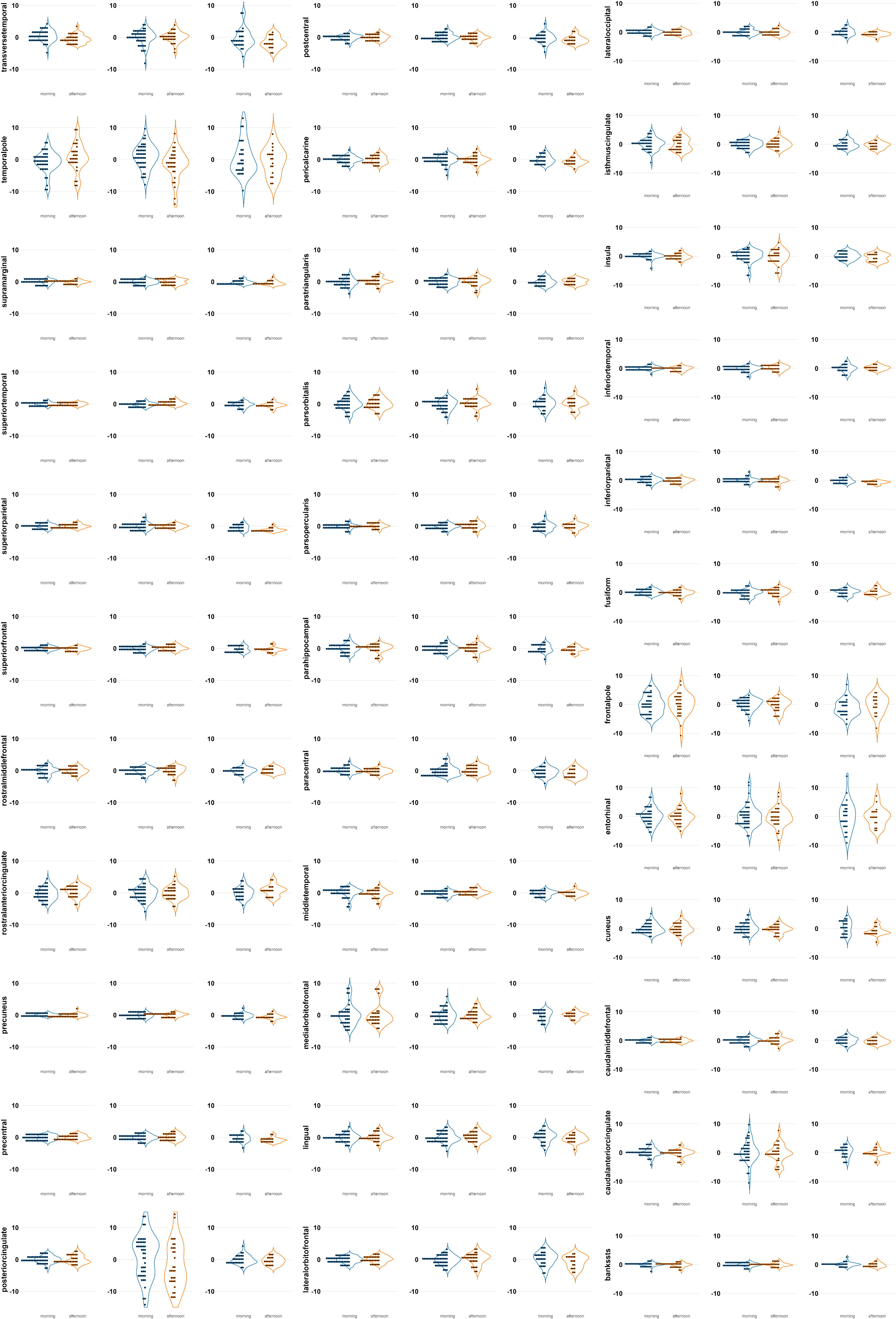

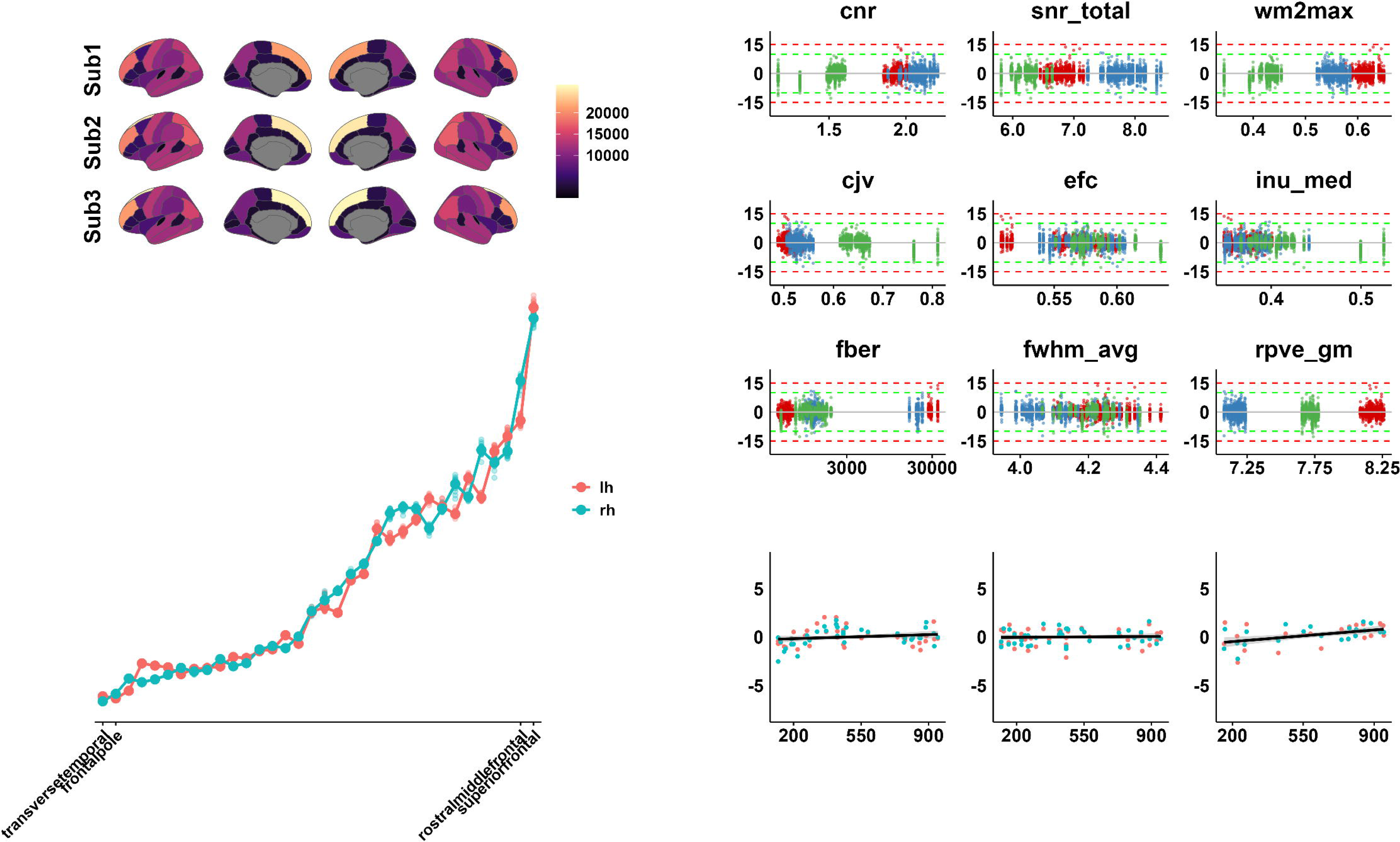

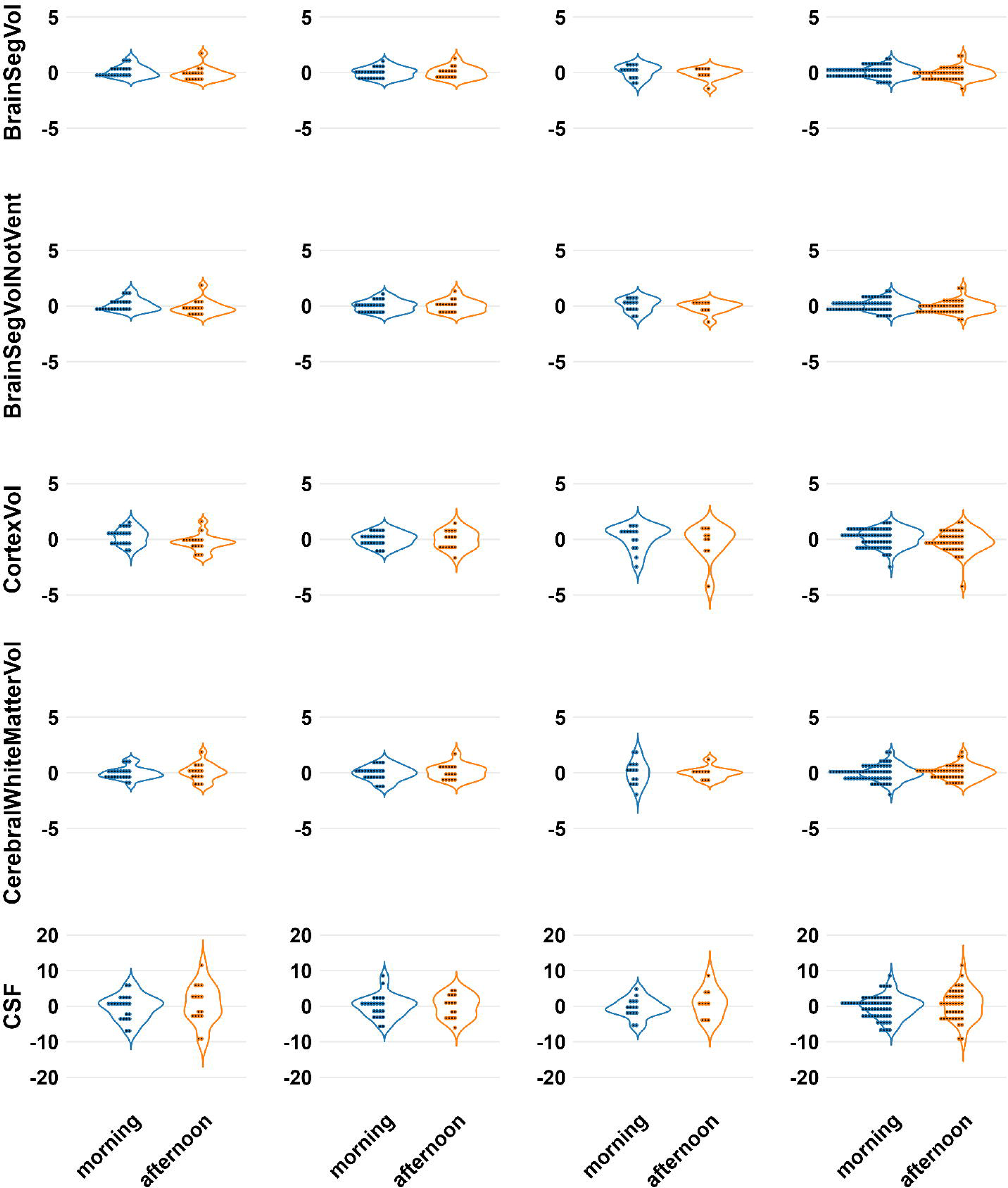

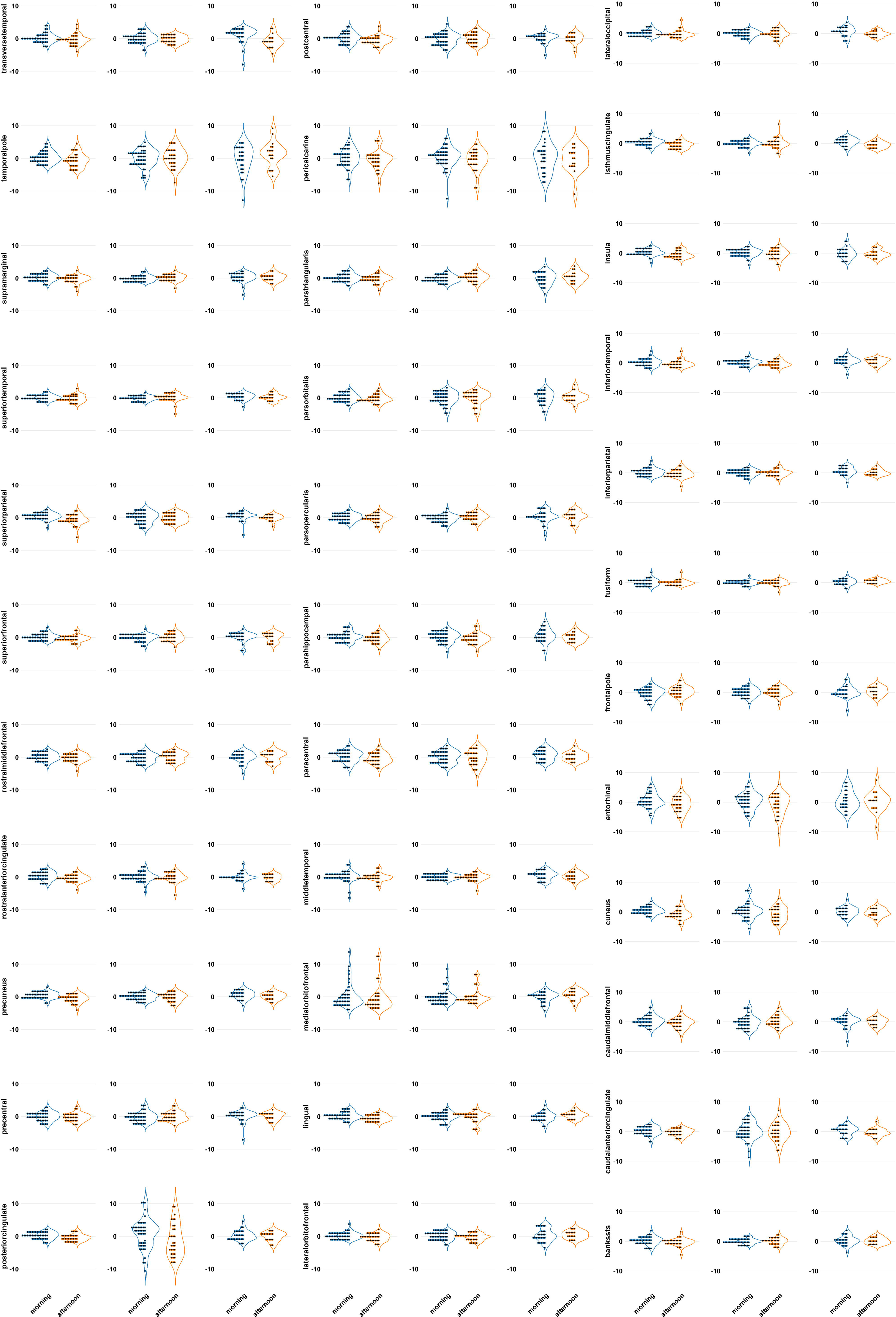

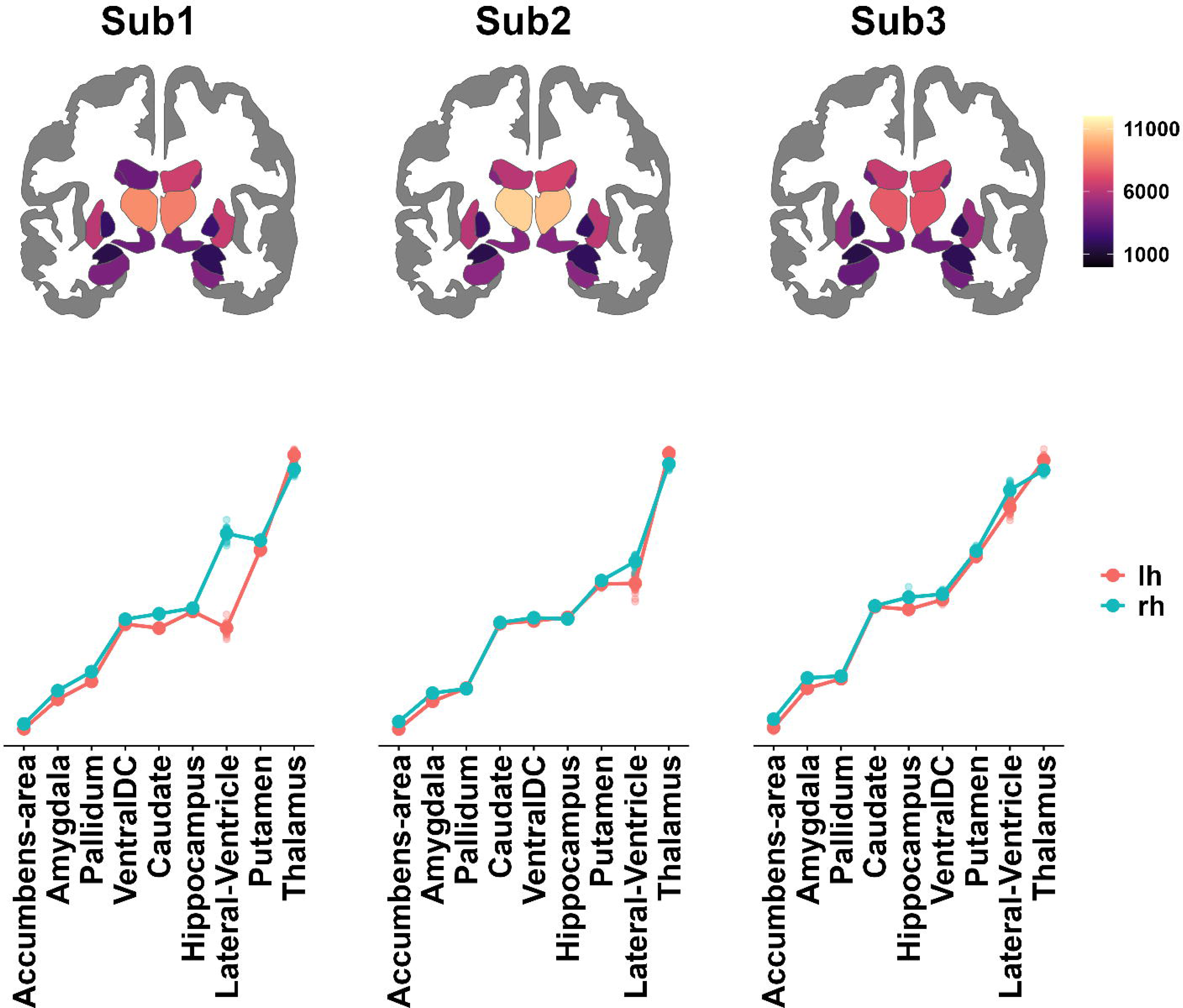

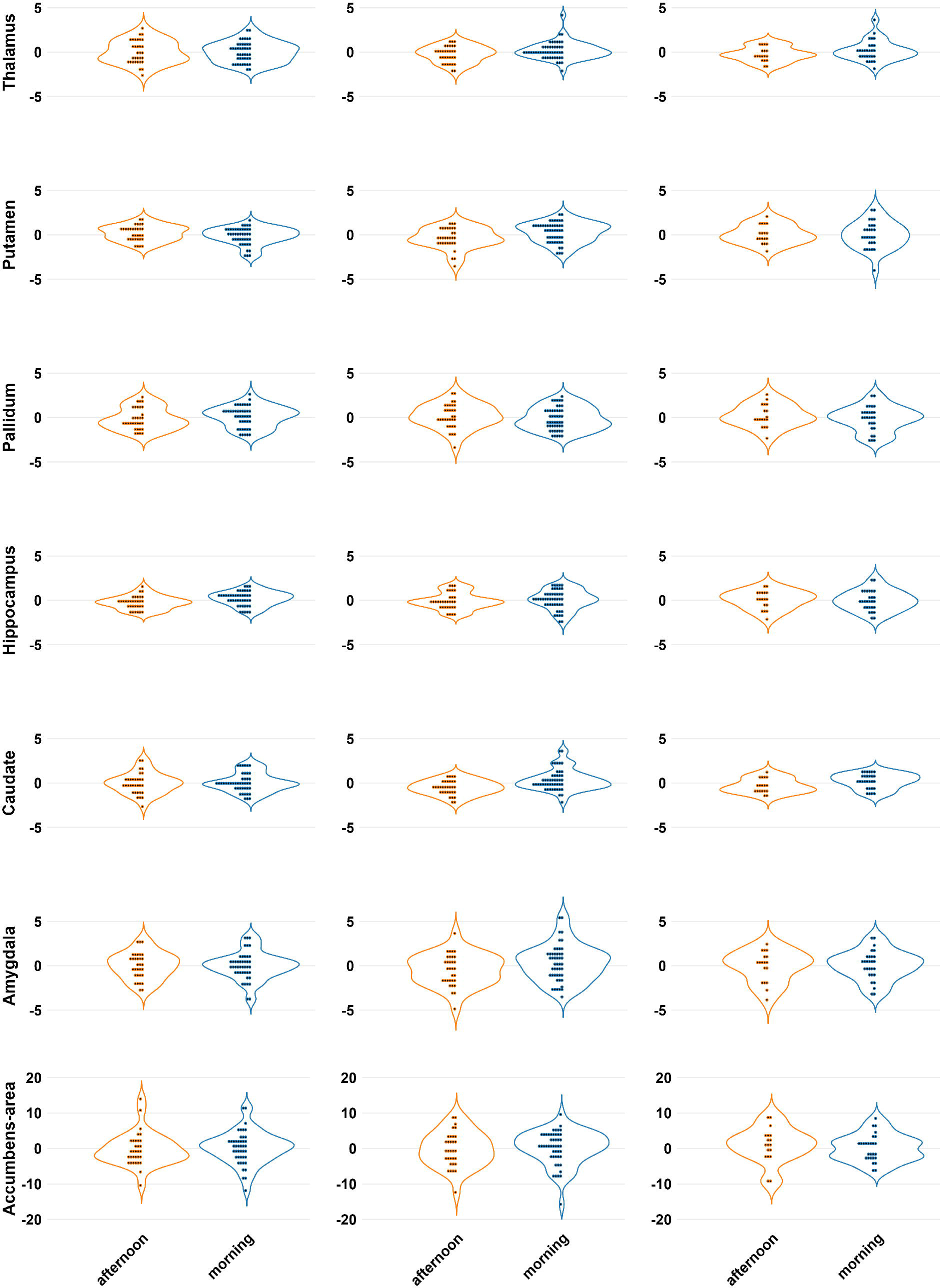

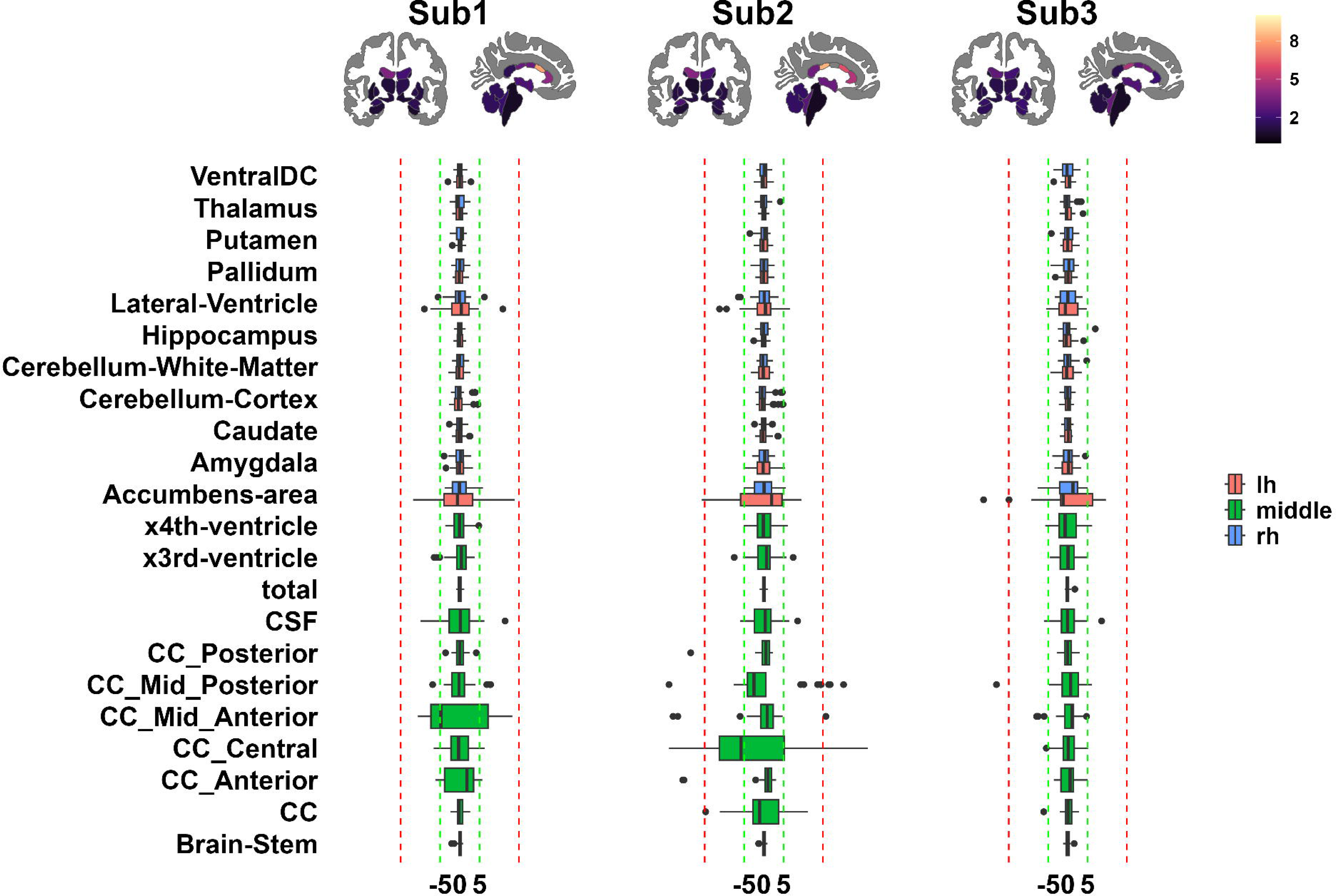

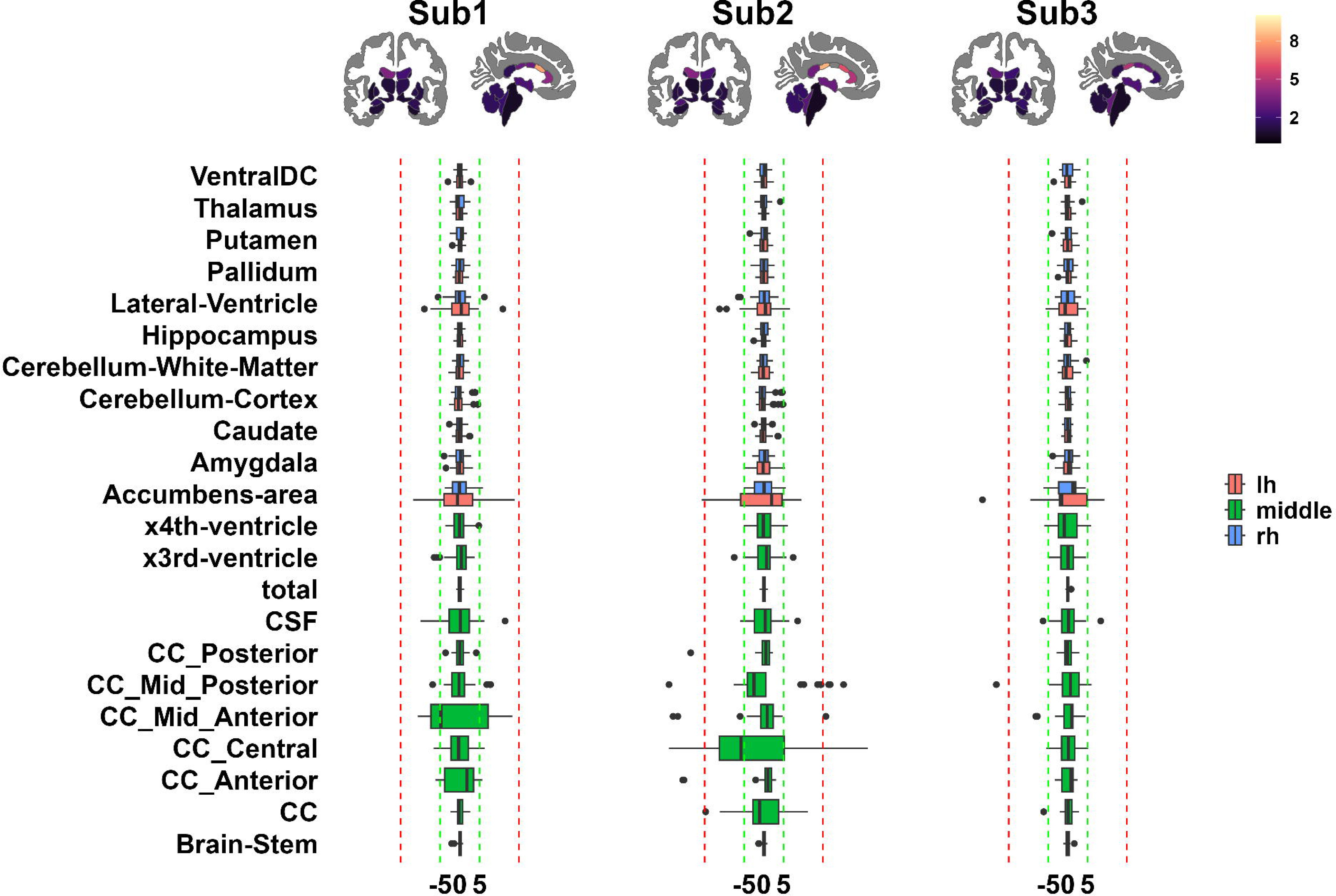

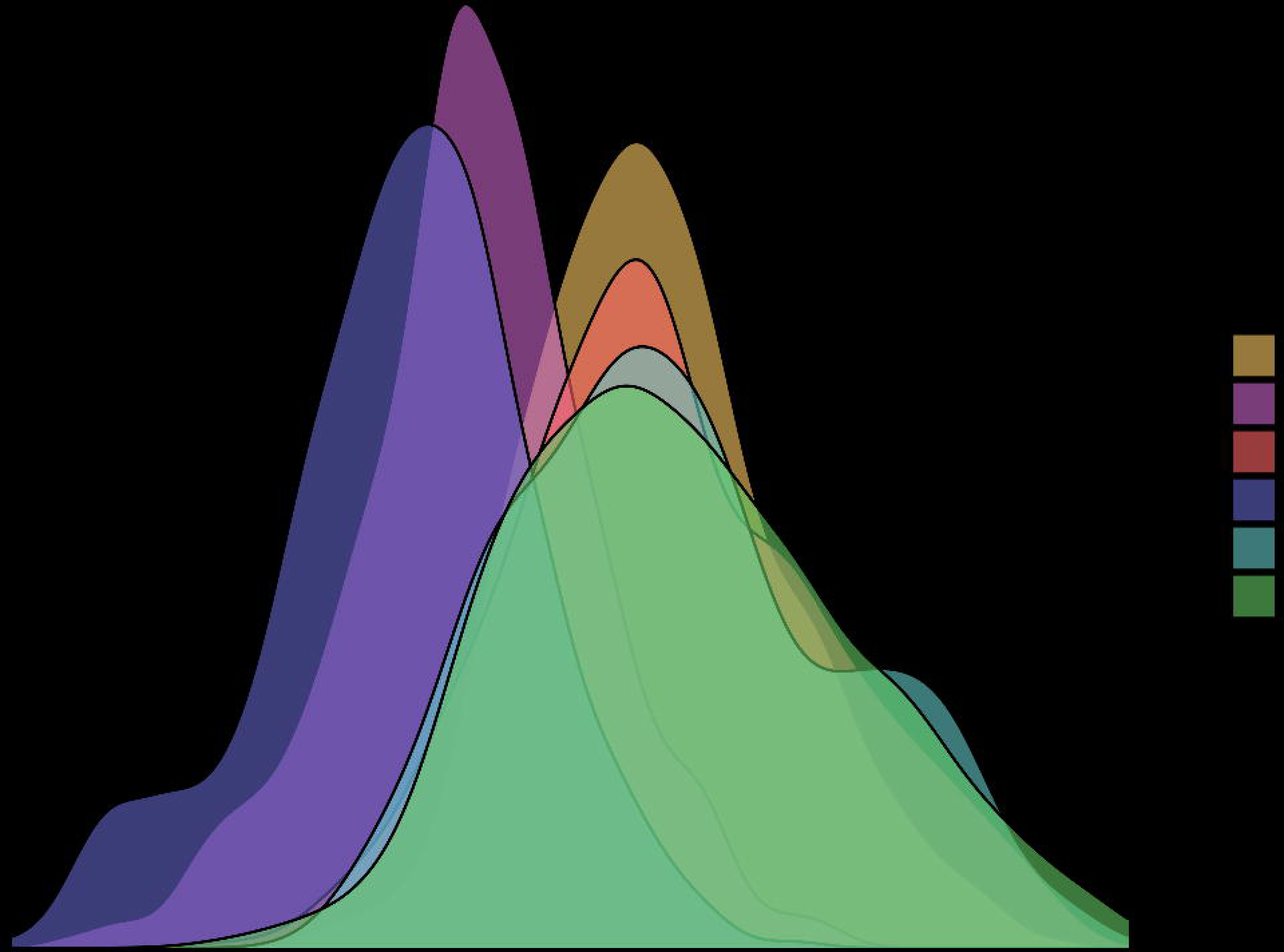

