## Supplementary Material for "The within-subject stability of cortical thickness, surface area, and brain volumes across one year"

**Data processing**

Cortical reconstruction and volumetric segmentation was performed with the Freesurfer image analysis suite, which is documented and freely available for download online (<http://surfer.nmr.mgh.harvard.edu/>). The technical details of these procedures are described in prior publications (Dale et al., 1999; Dale and Sereno, 1993; Fischl and Dale, 2000; Fischl et al., 2001; Fischl et al., 2002; Fischl et al., 2004a; Fischl et al., 1999a; Fischl et al., 1999b; Fischl et al., 2004b; Han et al., 2006; Jovicich et al., 2006; Segonne et al., 2004, Reuter et al. 2010, Reuter et al. 2012). Briefly, this processing includes motion correction and averaging (Reuter et al. 2010) of multiple volumetric T1 weighted images (when more than one is available), removal of non-brain tissue using a hybrid watershed/surface deformation procedure (Segonne et al., 2004), automated Talairach transformation, segmentation of the subcortical white matter and deep gray matter volumetric structures (including hippocampus, amygdala, caudate, putamen, ventricles) (Fischl et al., 2002; Fischl et al., 2004a) intensity normalization (Sled et al., 1998), tessellation of the gray matter white matter boundary, automated topology correction (Fischl et al., 2001; Segonne et al., 2007), and surface deformation following intensity gradients to optimally place the gray/white and gray/cerebrospinal fluid borders at the location where the greatest shift in intensity defines the transition to the other tissue class (Dale et al., 1999; Dale and Sereno, 1993; Fischl and Dale, 2000). Once the cortical models are complete, a number of deformable procedures can be performed for further data processing and analysis including surface inflation (Fischl et al., 1999a), registration to a spherical atlas which is based on individual cortical folding patterns to match cortical geometry across subjects (Fischl et al., 1999b), parcellation of the cerebral cortex into units with respect to gyral and sulcal structure (Desikan et al., 2006; Fischl et al., 2004b), and creation of a variety of surface based data including maps of curvature and sulcal depth. This method uses both intensity and continuity information from the entire three-dimensional MR volume in segmentation and deformation procedures to produce representations of cortical thickness, calculated as the closest distance from the gray/white boundary to the gray/CSF boundary at each vertex on the tessellated surface (Fischl and Dale, 2000). The maps are created using spatial intensity gradients across tissue classes and are therefore not simply reliant on absolute signal intensity. The maps produced are not restricted to the voxel resolution of the original data and thus are capable of detecting submillimeter differences between groups. Procedures for the measurement of cortical thickness have been validated against histological analysis (Rosas et al., 2002) and manual measurements (Kuperberg et al., 2003; Salat et al., 2004). Freesurfer morphometric procedures have been demonstrated to show good test-retest reliability across scanner manufacturers and field strengths (Han et al., 2006; Reuter et al., 2012).

**Aseg Atlas Information**
The aseg atlas is built from 40 subjects acquired using the same mp-rage sequence (by people at Wash U ages ago in collaboration with Randy Buckner). The subjects that make up the atlas are distributed in 4 groups of 10 subjects each: (1) young, (2) middle aged, (3) healthy older adults, (4) older adults with AD.

**Longitudinal Processing**
To extract reliable volume and thickness estimates, images were automatically processed with the longitudinal stream (Reuter et al., 2012) in FreeSurfer. Specifically, an unbiased within-subject template space and image is created using robust, inverse consistent registration (Reuter et al., 2010). Several processing steps, such as skull stripping, Talairach transforms, atlas registration as well as spherical surface maps and parcellations are then initialized with common information from the within-subject template, significantly increasing reliability and statistical power (Reuter et al., 2012).

**Table S1.** The scanning time and daylight length of each session

| Session | Sub1 | | | Sub2 | | | Sub3 | | |
| --- | --- | --- | --- | --- | --- | --- | --- | --- | --- |
|  | Date | Morning = 0 | Daylight | Date | Morning = 0 | Daylight | Date | Morning = 0 | Daylight |
| 1 | 20/01/2021 | 0 | 200 | 29/01/2021 | 0 | 244 | 03/02/2021 | 0 | 283 |
| 2 | 03/02/2021 | 0 | 283 | 02/02/2021 | 0 | 275 | 17/02/2021 | 0 | 427 |
| 3 | 12/02/2021 | 0 | 369 | 05/02/2021 | 0 | 301 | 24/02/2021 | 0 | 463 |
| 4 | 17/02/2021 | 0 | 427 | 12/02/2021 | 0 | 369 | 12/03/2021 | 0 | 552 |
| 5 | 17/02/2021 | 1 | 427 | 17/02/2021 | 1 | 427 | 17/03/2021 | 0 | 586 |
| 6 | 23/02/2021 | 0 | 458 | 19/02/2021 | 0 | 437 | 26/03/2021 | 0 | 639 |
| 7 | 24/02/2021 | 1 | 463 | 23/02/2021 | 0 | 458 | 08/04/2021 | 1 | 739 |
| 8 | 26/02/2021 | 0 | 473 | 24/02/2021 | 1 | 463 | 08/04/2021 | 1 | 739 |
| 9 | 12/03/2021 | 0 | 552 | 25/02/2021 | 1 | 468 | 14/04/2021 | 0 | 773 |
| 10 | 17/03/2021 | 0 | 586 | 10/03/2021 | 0 | 538 | 15/04/2021 | 1 | 778 |
| 11 | 08/04/2021 | 1 | 739 | 12/03/2021 | 0 | 552 | 21/04/2021 | 0 | 808 |
| 12 | 14/04/2021 | 0 | 773 | 16/03/2021 | 0 | 579 | 22/04/2021 | 1 | 812 |
| 13 | 15/04/2021 | 1 | 778 | 07/04/2021 | 0 | 731 | 28/04/2021 | 0 | 847 |
| 14 | 22/04/2021 | 1 | 812 | 08/04/2021 | 1 | 739 | 29/04/2021 | 1 | 853 |
| 15 | 23/04/2021 | 0 | 818 | 15/04/2021 | 1 | 778 | 05/05/2021 | 0 | 885 |
| 16 | 29/04/2021 | 1 | 853 | 16/04/2021 | 0 | 782 | 20/05/2021 | 1 | 930 |
| 17 | 30/04/2021 | 0 | 858 | 22/04/2021 | 1 | 812 | 21/05/2021 | 0 | 933 |
| 18 | 05/05/2021 | 0 | 885 | 28/04/2021 | 0 | 847 | 26/05/2021 | 0 | 945 |
| 19 | 06/05/2021 | 1 | 888 | 29/04/2021 | 1 | 853 | 27/05/2021 | 1 | 948 |
| 20 | 19/05/2021 | 0 | 927 | 06/05/2021 | 1 | 888 | 09/11/2021 | 0 | 270 |
| 21 | 20/05/2021 | 1 | 930 | 07/05/2021 | 0 | 891 | 10/11/2021 | 1 | 260 |
| 22 | 27/05/2021 | 1 | 948 | 19/05/2021 | 0 | 927 | 16/11/2021 | 0 | 226 |
| 23 | 03/11/2021 | 0 | 322 | 20/05/2021 | 1 | 930 | 17/11/2021 | 1 | 223 |
| 24 | 09/11/2021 | 0 | 270 | 28/05/2021 | 0 | 949 | 30/11/2021 | 0 | 163 |
| 25 | 10/11/2021 | 1 | 260 | 10/11/2021 | 0 | 260 | 01/12/2021 | 1 | 159 |
| 26 | 17/11/2021 | 1 | 221 | 10/11/2021 | 1 | 260 |  |  |  |
| 27 | 24/11/2021 | 0 | 189 | 16/11/2021 | 0 | 226 |  |  |  |
| 28 | 24/11/2021 | 1 | 189 | 16/11/2021 | 1 | 226 |  |  |  |
| 29 | 01/12/2021 | 1 | 159 | 24/11/2021 | 0 | 189 |  |  |  |
| 30 | 03/12/2021 | 0 | 151 | 24/11/2021 | 1 | 189 |  |  |  |
| 31 | 08/12/2021 | 0 | 133 | 30/11/2021 | 0 | 163 |  |  |  |
| 32 | 15/12/2021 | 0 | 120 | 01/12/2021 | 1 | 159 |  |  |  |
| 33 | 15/12/2021 | 1 | 120 | 08/12/2021 | 0 | 133 |  |  |  |
| 34 | 11/02/2022 | 0 | 360 | 15/12/2021 | 0 | 120 |  |  |  |
| 35 | 16/02/2022 | 0 | 421 | 15/12/2021 | 1 | 120 |  |  |  |
| 36 | 16/02/2022 | 1 | 421 | 09/02/2022 | 0 | 340 |  |  |  |
| 37 | 23/02/2022 | 0 | 458 | 16/02/2022 | 0 | 421 |  |  |  |
| 38 | 23/02/2022 | 1 | 458 | 16/02/2022 | 1 | 421 |  |  |  |
| 39 |  |  |  | 23/02/2022 | 0 | 458 |  |  |  |
| 40 |  |  |  | 23/02/2022 | 1 | 458 |  |  |  |

The unit of daylight is minute. Sessions 1 and 7 in red of sub3 were excluded for analysis of the cortical IDPs.

**Image quality checking**

The T1w images were quality assessed with MRIQC 23.1.0 (Esteban et al., 2017), where the following image quality metrics (IQMs) were used (**Fig. S1**): contrast-to-noise ratio (CNR); signal-to-noise ratio (SNR); white-matter to maximum intensity ratio (WM2MAX); coefficient of joint variation (CJV); entropy focus criterion (EFC); intensity non-uniformity (INU); foreground-to-background energy ratio (FBER); full-width half maximum (FWHM); residual partial volume effect feature (rPVE).

CNR measures the difference in mean intensities between gray and white matter, normalized by the standard deviation of non-brain areas, with higher values indicating better contrast between gray and white matter (Magnotta, Friedman, & First, 2006). SNR quantifies the mean intensity within gray matter relative to the standard deviation outside the brain, where higher values denote higher quality (Magnotta et al., 2006). WM2MAX estimates the median white matter intensity against the 95th percentile of the entire image intensity, pinpointing hyperintensities such as those in carotid vessels and fat with ideal values falling between 0.6 and 0.8 (Esteban et al., 2017).

CJV was proposed as a measure for INU correction algorithms, higher CJV indicates head motion or INU artifacts, whereas lower values imply better quality (Ganzetti, Wenderoth, & Mantini, 2016). EFC uses Shannon entropy to measure image blurring due to head motion with lower values reflecting a superior image clarity (Atkinson, Hill, Stoyle, Summers, & Keevil, 1997). INU summary statistics (max, min and median) of the INU field (bias field) as estimated with the N4ITK algorithm (Tustison et al., 2010). Values closer to 1.0 are better, values further from zero indicate greater field inhomogeneity.

FBER is defined as the mean energy of image values within the head relative to that outside the head with higher values indicating better images. The FWHM of the spatial distribution represents the image intensity values in voxel units where lower values are better with higher values indicating a blurrier image (Forman et al., 1995). The rPVE is a tissue-wise sum of partial volumes that fall in the range [5-95%] of the total volume of a pixel, computed on the partial volume maps, in which a lower rPVE is better (Esteban et al., 2017).

Atkinson, D., Hill, D. L., Stoyle, P. N., Summers, P. E., & Keevil, S. F. (1997). Automatic correction of motion artifacts in magnetic resonance images using an entropy focus criterion. *IEEE Trans Med Imaging, 16*(6), 903-910. doi:10.1109/42.650886

Esteban, O., Birman, D., Schaer, M., Koyejo, O. O., Poldrack, R. A., & Gorgolewski, K. J. (2017). MRIQC: Advancing the automatic prediction of image quality in MRI from unseen sites. *Plos One, 12*(9), e0184661. doi:10.1371/journal.pone.0184661

Forman, S. D., Cohen, J. D., Fitzgerald, M., Eddy, W. F., Mintun, M. A., & Noll, D. C. (1995). Improved assessment of significant activation in functional magnetic resonance imaging (fMRI): use of a cluster-size threshold. *Magn Reson Med, 33*(5), 636-647. doi:10.1002/mrm.1910330508

Ganzetti, M., Wenderoth, N., & Mantini, D. (2016). Intensity Inhomogeneity Correction of Structural MR Images: A Data-Driven Approach to Define Input Algorithm Parameters. *Frontiers in Neuroinformatics, 10*, 10. doi:10.3389/fninf.2016.00010

Magnotta, V. A., Friedman, L., & First, B. (2006). Measurement of Signal-to-Noise and Contrast-to-Noise in the fBIRN Multicenter Imaging Study. *Journal of Digital Imaging, 19*(2), 140-147. doi:10.1007/s10278-006-0264-x

Tustison, N. J., Avants, B. B., Cook, P. A., Zheng, Y., Egan, A., Yushkevich, P. A., & Gee, J. C. (2010). N4ITK: improved N3 bias correction. *IEEE Trans Med Imaging, 29*(6), 1310-1320. doi:10.1109/TMI.2010.2046908

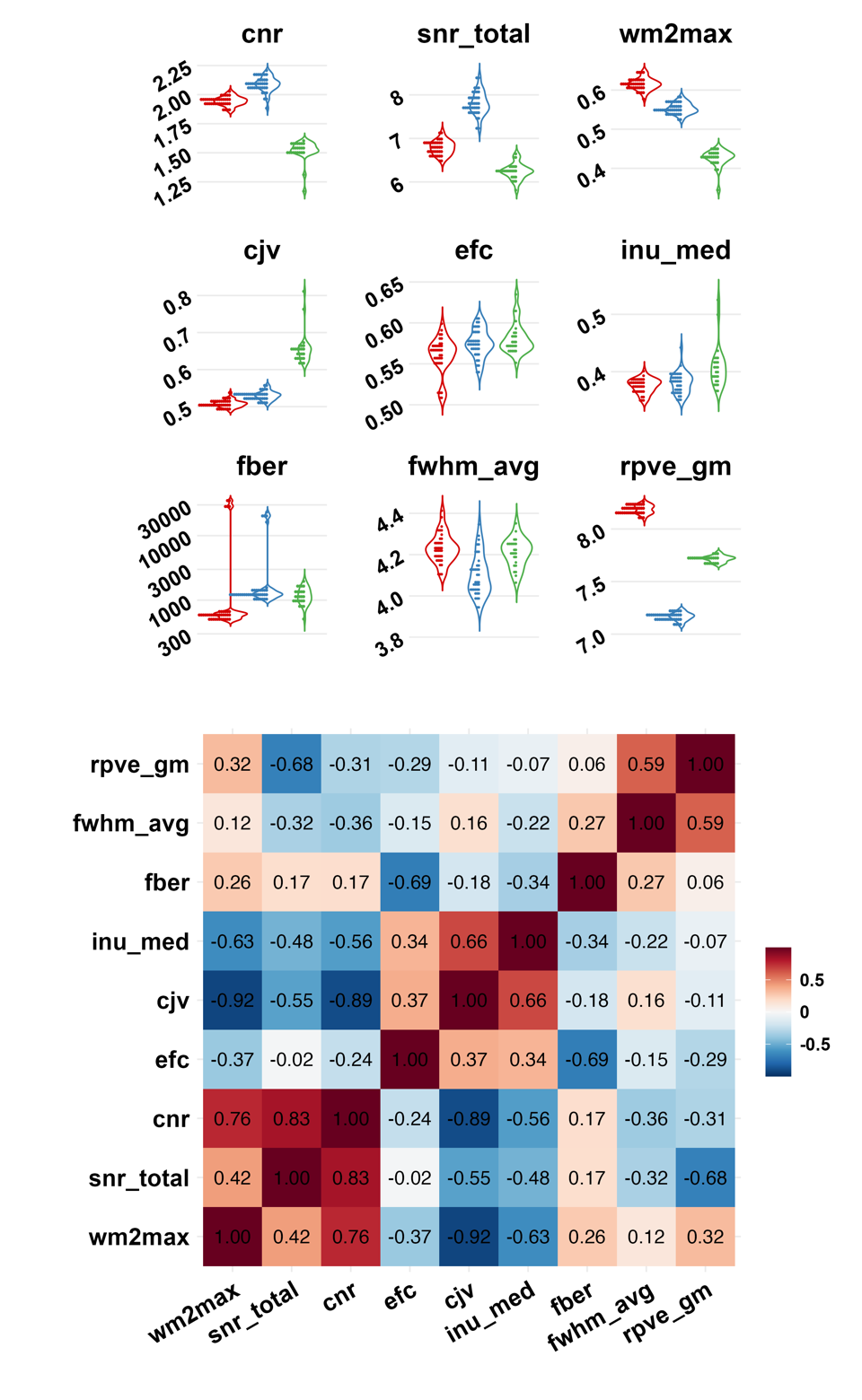

**Fig. S1. Distributions and correlations of IQMs and percentage changes of different brain tissues from different streams.** The upper panel of the violin plot of the IQM distributions for each subject where the red color represents subject 1, the blue depicts subject 2, and the green represents subject 3. The bottom panel of the correlation matrix between the IQMs, where the cnr, snr_total, and wm2max form one cluster representing the image intensity, the cjv, efc, and inu_med constructs another cluster representing head motion or related artifacts, and the fber, fwhm_avg, and rpve_gm form the last cluster representing the technique qualities. Contrast-to-noise ratio (CNR); signal-to-noise ratio (SNR); white-matter to maximum intensity ratio (wm2max); coefficient of joint variation (CJV); entropy-focus criterion (EFC); intensity non-uniformity (INU); foreground-to-background energy ratio (FBER); full-width half-maximum (FWHM); residual partial volume effect (rPVE).

**Cortical thickness**

The mean cortical thicknesses of the three subjects were 2.59 mm, 2.70 mm, and 2.58 mm, respectively with a shared standard deviation of 0.02 mm (**Table S2)**. In addition, subject 1 showed a range of 1.59 to 4.04 mm, subject 2 with a range of 1.83 to 3.75 mm, and subject 3 with a range of 1.50 to 4.03 mm (**Table S2)**. Moreover, analysis of the cortical thickness distribution across various brain regions revealed a consistent ranking of regions within subjects, identifying the insula, entorhinal cortex, and temporal pole as the thickest regions for all subjects (**Fig. S2AB** and **Table S2**). Conversely, regions such as the lingual gyrus, cuneus, and pericalcarine cortex were consistently ranked as the thinnest across the subjects, as also shown in **Fig. S2AB** and **Table S2**. Furthermore, standard deviations ranged from 0.02 to 0.15 mm, where the thickest brain regions had the largest standard deviations (**Table S2**).

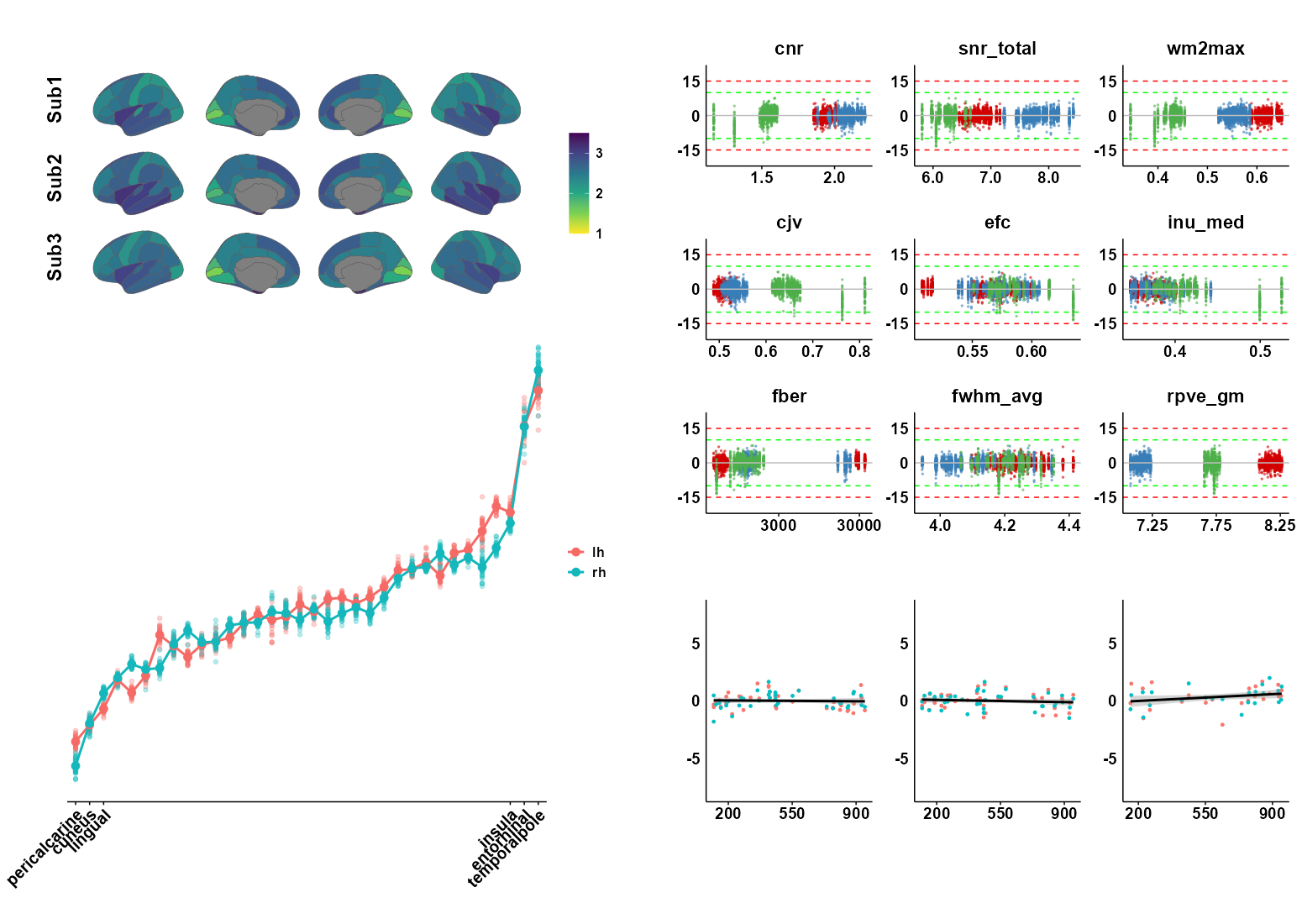

**Fig. S2. Mean cortical thickness for each brain region, percentage change distributions along with IQMs, and percentage changes of average thickness along the daylight length.** **(A)** Average cortical thickness across all sessions of different brain regions for each subject where the unit is a millimeter. **(B)** Ranked brain regions are illustrated from the thinnest to the thickest (example data from sub1). All three subjects have the same the most three thinnest and thickest brain regions although the brain regions ranking in the middle vary. **(C)** The distribution of the percentage changes along with different IQMs, where the red, blue, and green dots represent subjects 1, 2, and 3, respectively. Each vertical line represents one session with which the dots denote the different regions. Sub3 showed two sessions with outliers, which were sessions 1 and 7. **(D)** Percentage change (y-axis) of average thickness along different daylight lengths (x-axis with a unit of minute) for sub1, sub2, and sub3, respectively.

**Table S2.** Mean cortical thickness across sessions for each subject based on DK atlas

|  | Sub1 | | Sub2 | | Sub3 | | | | Ranking | | | | | |
| --- | --- | --- | --- | --- | --- | --- | --- | --- | --- | --- | --- | --- | --- | --- |
|  | L | R | L | R | L | L_qc | R | R_qc |  |  |  |  |  |  |
| temporalpole | 3.91±0.07 | 4.04±0.09 | 3.75±0.10 | 3.71±0.12 | 3.41±0.16 | 3.43±0.15 | 4.02±0.13 | 4.03±0.13 | 34 | 34 | 34 | 34 | 34 | 34 |
| entorhinal | 3.69±0.07 | 3.69±0.06 | 3.56±0.08 | 3.29±0.08 | 3.31±0.12 | 3.33±0.11 | 3.39±0.08 | 3.40±0.05 | 33 | 33 | 33 | 33 | 33 | 33 |
| insula | 3.16±0.04 | 3.09±0.04 | 3.20±0.03 | 3.16±0.04 | 2.96±0.06 | 2.96±0.05 | 3.03±0.05 | 3.02±0.04 | 31 | 32 | 32 | 32 | 32 | 32 |
| parahippocampal | 3.20±0.03 | 2.94±0.03 | 2.97±0.03 | 2.78±0.05 | 2.61±0.07 | 2.62±0.04 | 2.66±0.07 | 2.68±0.05 | 32 | 31 | 28 | 24 | 19 | 25 |
| frontalpole | 3.04±0.07 | 2.82±0.06 | 2.73±0.08 | 2.65±0.06 | 2.92±0.05 | 2.93±0.04 | 2.94±0.12 | 2.96±0.09 | 30 | 27 | 17 | 16 | 30 | 31 |
| superiortemporal | 2.93±0.02 | 2.88±0.02 | 3.10±0.03 | 3.03±0.03 | 2.93±0.05 | 2.94±0.03 | 2.84±0.07 | 2.86±0.04 | 29 | 29 | 30 | 30 | 31 | 29 |
| middletemporal | 2.91±0.02 | 2.83±0.02 | 3.15±0.03 | 2.93±0.03 | 2.75±0.07 | 2.76±0.03 | 2.68±0.07 | 2.70±0.03 | 28 | 28 | 31 | 29 | 24 | 26 |
| fusiform | 2.85±0.03 | 2.82±0.02 | 2.98±0.03 | 2.92±0.02 | 2.80±0.04 | 2.80±0.03 | 2.91±0.05 | 2.92±0.02 | 27 | 26 | 29 | 28 | 28 | 30 |
| superiorfrontal | 2.81±0.03 | 2.81±0.03 | 2.86±0.03 | 2.89±0.03 | 2.69±0.05 | 2.70±0.04 | 2.75±0.06 | 2.76±0.04 | 26 | 25 | 25 | 27 | 23 | 28 |
| inferiortemporal | 2.80±0.03 | 2.75±0.02 | 2.95±0.03 | 3.09±0.02 | 2.80±0.06 | 2.81±0.03 | 2.73±0.07 | 2.74±0.03 | 25 | 24 | 27 | 31 | 29 | 27 |
| rostralanteriorcingulate | 2.77±0.04 | 2.90±0.04 | 2.81±0.03 | 2.68±0.04 | 2.78±0.04 | 2.78±0.03 | 2.65±0.04 | 2.64±0.03 | 24 | 30 | 21 | 17 | 27 | 22 |
| precentral | 2.70±0.02 | 2.63±0.03 | 2.83±0.03 | 2.72±0.03 | 2.76±0.09 | 2.77±0.06 | 2.21±0.04 | 2.22±0.03 | 23 | 23 | 24 | 19 | 26 | 5 |
| parsorbitalis | 2.63±0.03 | 2.53±0.03 | 2.83±0.06 | 2.77±0.05 | 2.67±0.07 | 2.67±0.07 | 2.60±0.05 | 2.60±0.05 | 22 | 19 | 23 | 23 | 21 | 19 |
| parsopercularis | 2.63±0.03 | 2.53±0.03 | 2.76±0.04 | 2.74±0.03 | 2.65±0.07 | 2.66±0.06 | 2.50±0.05 | 2.51±0.05 | 21 | 18 | 19 | 21 | 20 | 16 |
| bankssts | 2.62±0.02 | 2.48±0.05 | 2.76±0.03 | 2.59±0.03 | 2.53±0.08 | 2.55±0.05 | 2.62±0.06 | 2.63±0.03 | 20 | 14 | 18 | 12 | 13 | 21 |
| lateralorbitofrontal | 2.59±0.03 | 2.57±0.03 | 2.71±0.05 | 2.79±0.04 | 2.58±0.06 | 2.59±0.06 | 2.66±0.07 | 2.68±0.05 | 19 | 22 | 16 | 25 | 16 | 24 |
| caudalmiddlefrontal | 2.59±0.04 | 2.49±0.03 | 2.90±0.04 | 2.69±0.04 | 2.61±0.05 | 2.61±0.05 | 2.53±0.05 | 2.54±0.04 | 18 | 16 | 26 | 18 | 18 | 17 |
| posteriorcingulate | 2.54±0.03 | 2.56±0.03 | 2.45±0.07 | 2.57±0.05 | 2.60±0.04 | 2.60±0.03 | 2.40±0.04 | 2.41±0.03 | 17 | 21 | 8 | 11 | 17 | 9 |
| supramarginal | 2.52±0.03 | 2.48±0.03 | 2.77±0.03 | 2.76±0.03 | 2.53±0.06 | 2.54±0.05 | 2.64±0.05 | 2.65±0.04 | 16 | 15 | 20 | 22 | 12 | 23 |
| paracentral | 2.51±0.04 | 2.53±0.05 | 2.50±0.05 | 2.53±0.04 | 2.53±0.09 | 2.55±0.04 | 2.44±0.07 | 2.46±0.04 | 15 | 17 | 10 | 10 | 14 | 13 |
| medialorbitofrontal | 2.49±0.06 | 2.54±0.05 | 2.68±0.05 | 2.80±0.04 | 2.41±0.04 | 2.42±0.04 | 2.43±0.04 | 2.43±0.04 | 14 | 20 | 15 | 26 | 9 | 11 |
| caudalanteriorcingulate | 2.47±0.04 | 2.47±0.04 | 2.45±0.05 | 2.46±0.03 | 2.77±0.05 | 2.77±0.04 | 2.42±0.05 | 2.42±0.04 | 13 | 13 | 7 | 9 | 25 | 10 |
| transversetemporal | 2.40±0.04 | 2.19±0.03 | 2.81±0.05 | 2.72±0.04 | 2.67±0.12 | 2.68±0.11 | 2.59±0.09 | 2.61±0.05 | 12 | 6 | 22 | 20 | 22 | 20 |
| parstriangularis | 2.38±0.03 | 2.46±0.03 | 2.65±0.04 | 2.65±0.04 | 2.57±0.07 | 2.58±0.06 | 2.43±0.05 | 2.43±0.05 | 11 | 12 | 14 | 15 | 15 | 12 |
| inferiorparietal | 2.36±0.04 | 2.36±0.05 | 2.50±0.03 | 2.62±0.03 | 2.40±0.08 | 2.42±0.04 | 2.59±0.06 | 2.60±0.03 | 10 | 10 | 9 | 14 | 8 | 18 |
| rostralmiddlefrontal | 2.34±0.03 | 2.35±0.02 | 2.63±0.04 | 2.45±0.04 | 2.43±0.06 | 2.44±0.04 | 2.38±0.04 | 2.38±0.03 | 9 | 9 | 13 | 8 | 10 | 7 |
| isthmuscingulate | 2.33±0.03 | 2.34±0.04 | 2.52±0.03 | 2.39±0.03 | 2.52±0.04 | 2.52±0.03 | 2.47±0.03 | 2.47±0.03 | 8 | 8 | 12 | 5 | 11 | 14 |
| precuneus | 2.26±0.03 | 2.43±0.03 | 2.50±0.03 | 2.60±0.03 | 2.38±0.05 | 2.39±0.03 | 2.37±0.06 | 2.39±0.03 | 7 | 11 | 11 | 13 | 7 | 8 |
| superiorparietal | 2.15±0.03 | 2.19±0.03 | 2.34±0.03 | 2.39±0.03 | 2.28±0.06 | 2.30±0.04 | 2.29±0.07 | 2.31±0.02 | 6 | 5 | 5 | 6 | 6 | 6 |
| postcentral | 2.12±0.02 | 2.13±0.02 | 2.30±0.03 | 2.31±0.03 | 2.17±0.05 | 2.18±0.05 | 2.46±0.05 | 2.47±0.03 | 5 | 4 | 4 | 4 | 5 | 15 |
| lateraloccipital | 2.04±0.03 | 2.22±0.02 | 2.35±0.03 | 2.40±0.02 | 2.06±0.04 | 2.06±0.03 | 2.14±0.04 | 2.15±0.03 | 4 | 7 | 6 | 7 | 3 | 4 |
| lingual | 1.94±0.02 | 2.04±0.02 | 2.26±0.03 | 2.21±0.04 | 2.18±0.03 | 2.18±0.02 | 1.97±0.04 | 1.98±0.04 | 3 | 3 | 3 | 3 | 4 | 3 |
| cuneus | 1.84±0.03 | 1.85±0.04 | 2.10±0.03 | 2.02±0.03 | 1.76±0.03 | 1.76±0.03 | 1.84±0.04 | 1.85±0.04 | 2 | 2 | 2 | 2 | 2 | 2 |
| pericalcarine | 1.74±0.03 | 1.59±0.04 | 1.90±0.04 | 1.83±0.06 | 1.70±0.06 | 1.70±0.06 | 1.51±0.05 | 1.50±0.05 | 1 | 1 | 1 | 1 | 1 | 1 |
| average | 2.60±0.02 | 2.57±0.02 | 2.72±0.02 | 2.68±0.02 | 2.58±0.04 | 2.59±0.02 | 2.56±0.04 | 2.57±0.02 |  |  |  |  |  |  |

Unit: millimeter; Red color: the three thickest brain regions; Blue color: the three thinnest brain regions; Green color: brain thickness after removing two sessions with excessive head movement

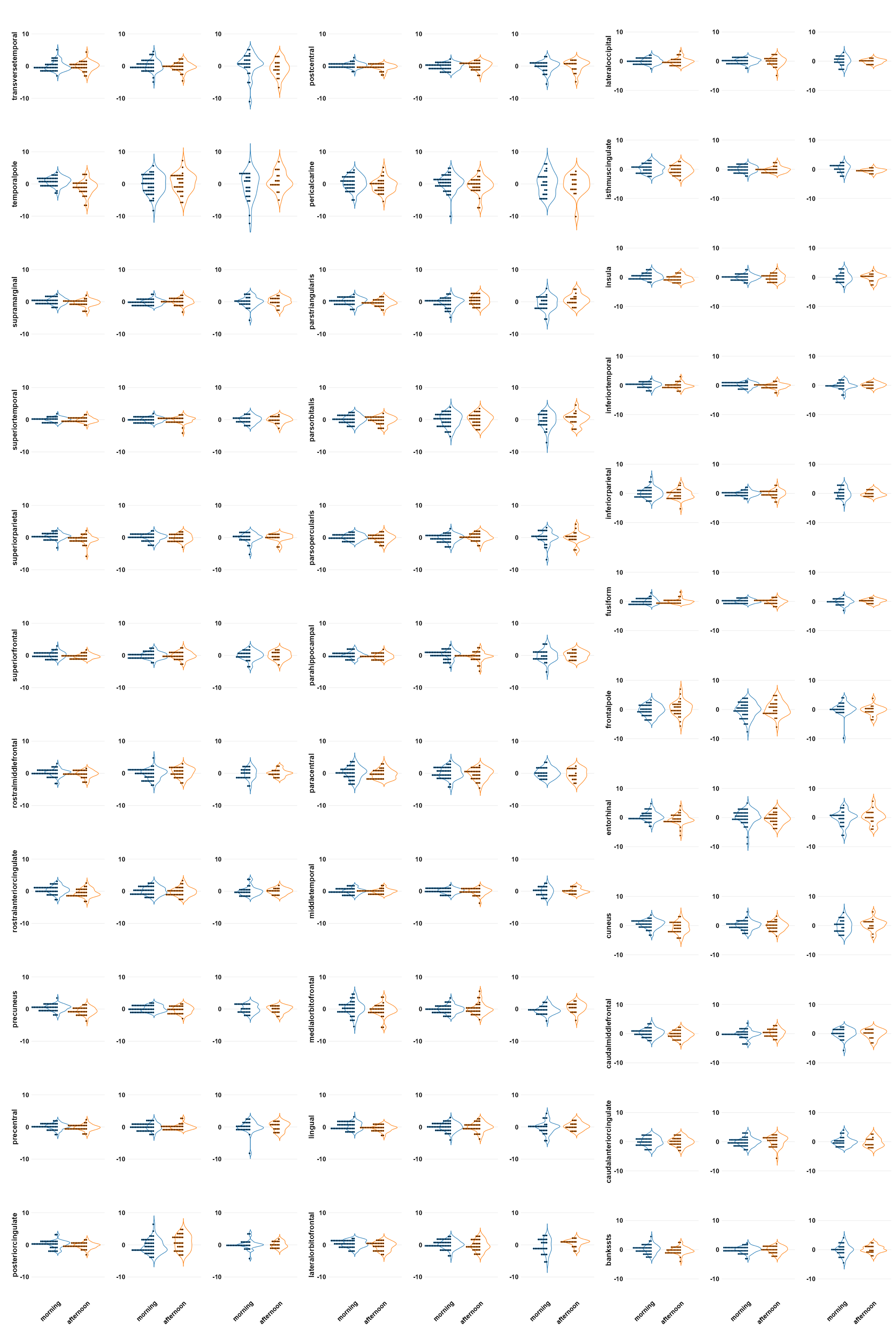

**Fig. S3. The percentage change distribution of time-of-day in the cortical thickness of DK atlas-defined brain regions.** From left to right, sub1, sub2, and sub3.

**Surface area**

The total surface areas of one brain hemisphere are around 850, 912, and 908 cm^2^ for sub1, sub2, and sub3, respectively. Additionally, the range of the surface area of one brain region in each brain hemisphere is from 219.82 to 6815.53 mm^2^ for sub1, 298.30 to 8046.82 mm^2^ for sub2, and 248.83 to 8594.52 mm^2^ for sub3 (**Table S3**). The superior parietal, rostral middle frontal, and superior frontal cortices emerged as the consistently largest surface areas in all subjects (**Fig. S4AB**). In contrast, the entorhinal, transverse temporal, and frontal pole regions were invariably the smallest (**Fig. S4AB** and **Table S3**).

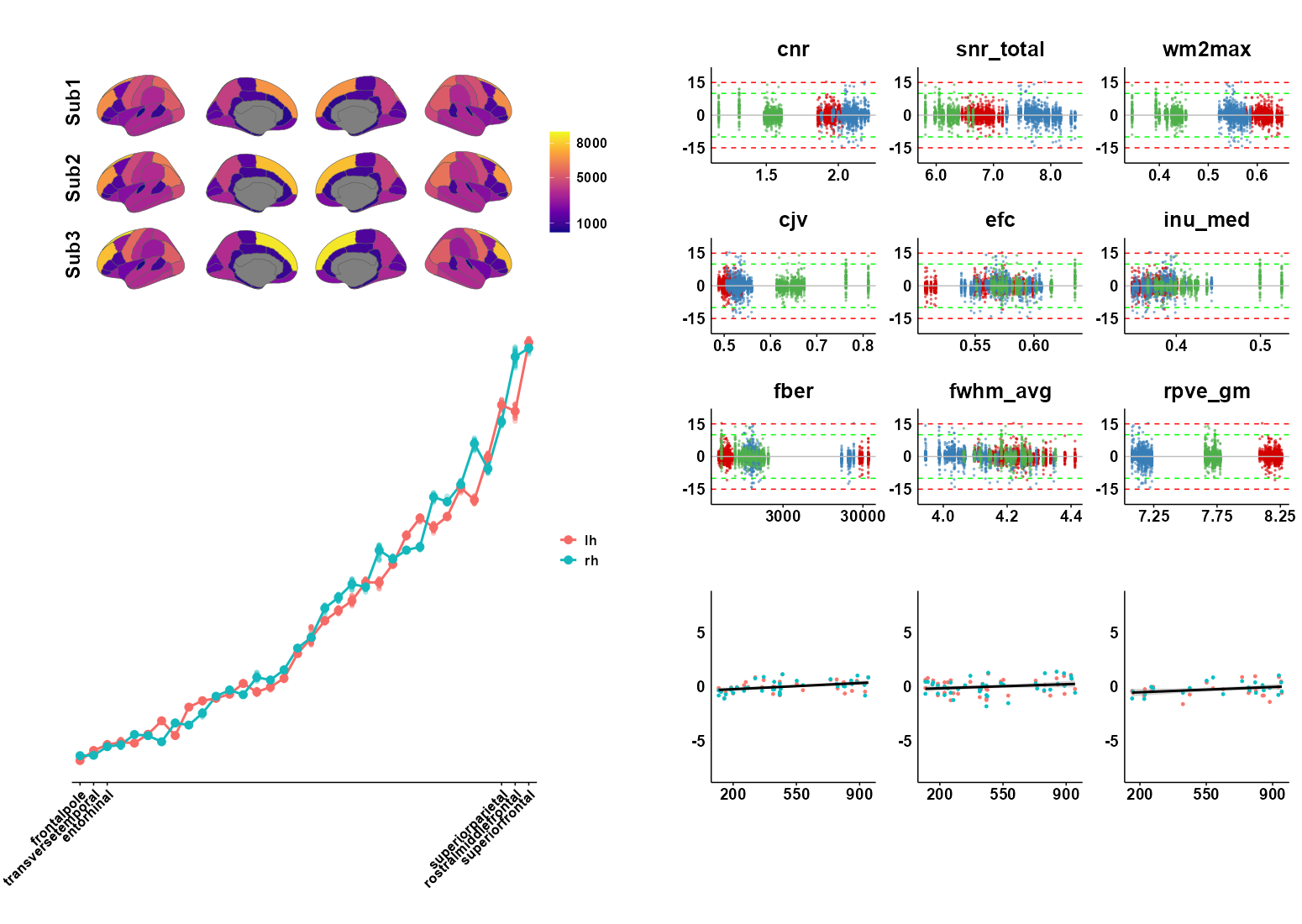

**Fig. S4. Mean surface area for each brain region, percentage change distribution along with IQMs, and percentage change of total surface area along the daylight length.** **(A)** The average surface area of different brain regions for each subject and the unit is millimeters per square. **(B)** Surface areas from the smallest to the largest (data from sub1). The three smallest and largest surface areas were the same in all three subjects, but the surface area ranking in the middle varied. **(C)** The distributions of surface area percentage changes along with the IQMs, where the red, blue, and green dots represent subjects 1, 2, and 3, respectively. Each vertical line represents one session with which the dots denote the different regions. Sub3 showed two sessions (1 and 7) with outliers. **(D)** The distribution of percentage changes (y-axis) of the total surface area along the daylight length (x-axis with a unit of minute) for each subject.

**Table S3.** Mean area size across sessions for each subject based on DK atlas

|  | Sub1 | | Sub2 | | Sub3 | | | | Ranking | | | | | |
| --- | --- | --- | --- | --- | --- | --- | --- | --- | --- | --- | --- | --- | --- | --- |
|  | L | R | L | R | L | L_qc | R | R_qc |  |  |  |  |  |  |
| superiorfrontal | 6815.53±38.39 | 6730.39±36.79 | 8046.82±61.05 | 7739.12±59.13 | 8070.00±124.33 | 8047.35±71.22 | 8627.32±137.21 | 8594.52±77.88 | 34 | 34 | 34 | 34 | 34 | 34 |
| rostralmiddlefrontal | 5732.58±62.88 | 6591.13±81.41 | 6475.52±65.21 | 6849.00±82.68 | 5935.68±81.00 | 5921.57±62.43 | 7738.92±79.95 | 7749.52±71.64 | 32 | 33 | 33 | 33 | 33 | 33 |
| superiorparietal | 5832.13±42.09 | 5566.18±32.70 | 5374.10±44.24 | 6185.10±57.14 | 5491.52±153.46 | 5449.35±49.49 | 3910.24±107.99 | 3883.91±35.36 | 33 | 32 | 31 | 32 | 31 | 28 |
| inferiorparietal | 4330.68±34.35 | 5220.21±46.77 | 5002.48±45.45 | 5851.90±48.07 | 5147.76±113.32 | 5117.57±37.7 | 5341.16±64.73 | 5326.83±35.05 | 29 | 31 | 30 | 31 | 30 | 31 |
| precentral | 5006.71±40.56 | 4825.58±36.76 | 4593.68±48.60 | 4571.15±36.63 | 4975.96±129.40 | 4942.65±61.38 | 5605.96±98.35 | 5581.3±49.51 | 31 | 30 | 29 | 29 | 29 | 32 |
| lateraloccipital | 4525.66±39.08 | 4574.71±36.50 | 5690.80±60.15 | 4705.55±40.30 | 5584.76±70.21 | 5573±51.05 | 4743.88±89.56 | 4732.78±58.41 | 30 | 29 | 32 | 30 | 32 | 30 |
| postcentral | 3900.16±37.66 | 4378.82±33.58 | 4148.85±46.11 | 3789.88±39.40 | 4486.16±131.57 | 4456.74±86.89 | 3770.56±51.47 | 3760.17±37.9 | 26 | 28 | 27 | 27 | 28 | 26 |
| precuneus | 4069.92±18.88 | 4304.18±22.53 | 4313.45±29.12 | 4375.82±29.49 | 3786.04±59.79 | 3779.43±31.2 | 3773.48±68.60 | 3755.52±28.94 | 28 | 27 | 28 | 28 | 23 | 25 |
| supramarginal | 4047.05±24.61 | 3591.97±23.36 | 3670.00±23.09 | 3701.28±30.40 | 4322.20±60.50 | 4305.91±22.63 | 3016.36±34.99 | 3011.91±22.43 | 27 | 26 | 25 | 26 | 27 | 20 |
| superiortemporal | 3772.39±22.34 | 3540.61±14.32 | 4082.55±29.72 | 3585.57±22.81 | 4266.44±58.75 | 4253.65±38.36 | 3770.12±37.51 | 3762.78±27.53 | 25 | 25 | 26 | 24 | 25 | 27 |
| middletemporal | 3031.05±36.96 | 3538.39±58.76 | 3475.28±29.98 | 3496.85±27.16 | 3895.64±50.60 | 3886.3±36.86 | 3677.72±35.63 | 3673.91±29.37 | 22 | 24 | 24 | 23 | 24 | 24 |
| inferiortemporal | 3317.53±17.46 | 3398.55±28.94 | 3374.12±37.44 | 3685.78±28.36 | 4273.60±52.72 | 4276.13±53.51 | 4018.44±45.26 | 4016.83±41.89 | 24 | 23 | 21 | 25 | 26 | 29 |
| lingual | 2741.79±38.12 | 3005.34±39.05 | 3400.68±50.86 | 3459.50±57.01 | 3048.32±46.57 | 3045.39±40.52 | 3603.40±71.72 | 3604.91±72.33 | 21 | 22 | 22 | 22 | 21 | 23 |
| fusiform | 3031.55±28.75 | 2960.08±26.21 | 3460.75±37.52 | 3295.20±44.02 | 3702.28±50.85 | 3708±45.51 | 3436.92±31.22 | 3440.48±29.95 | 23 | 21 | 23 | 21 | 22 | 22 |
| lateralorbitofrontal | 2587.50±25.15 | 2794.61±31.59 | 3160.75±49.60 | 2977.07±52.43 | 2828.96±67.41 | 2823.17±55.36 | 3031.16±68.57 | 3032.87±68.82 | 20 | 20 | 20 | 20 | 20 | 21 |
| insula | 2426.79±18.56 | 2622.97±29.94 | 2457.20±38.29 | 2565.20±72.94 | 2337.64±34.16 | 2334.52±33.61 | 2112.44±24.16 | 2112±24.41 | 19 | 19 | 19 | 19 | 18 | 17 |
| medialorbitofrontal | 2151.37±97.26 | 2162.76±21.33 | 2185.62±45.79 | 2401.15±38.65 | 2024.00±29.08 | 2023.04±29.05 | 2513.16±31.03 | 2513.26±32 | 18 | 18 | 17 | 18 | 17 | 18 |
| caudalmiddlefrontal | 1907.79±11.14 | 1993.68±10.96 | 2151.05±16.52 | 1927.15±24.87 | 2443.20±27.39 | 2445.26±27.11 | 2805.92±36.27 | 2799.87±30.31 | 17 | 17 | 16 | 17 | 19 | 19 |
| pericalcarine | 1517.58±20.93 | 1650.18±15.59 | 1614.97±27.40 | 1683.00±18.72 | 1391.84±32.12 | 1384.48±15.65 | 1669.72±23.19 | 1669.74±23.82 | 16 | 16 | 14 | 15 | 14 | 15 |
| cuneus | 1301.42±21.28 | 1533.66±29.39 | 1645.67±23.02 | 1710.25±33.19 | 1635.40±37.58 | 1630.96±32.41 | 1775.84±40.98 | 1773.83±42.1 | 13 | 15 | 15 | 16 | 16 | 16 |
| parstriangularis | 1379.13±21.71 | 1490.53±11.81 | 1576.80±21.98 | 1429.88±16.49 | 1083.48±25.68 | 1081±12.44 | 1390.52±15.33 | 1393±11.97 | 14 | 14 | 13 | 12 | 11 | 14 |
| paracentral | 1266.71±9.89 | 1331.26±13.70 | 1363.80±20.77 | 1612.58±21.09 | 1494.92±44.58 | 1485.43±23.12 | 1314.28±34.85 | 1305.74±16.03 | 12 | 13 | 12 | 14 | 15 | 13 |
| parsopercularis | 1436.05±13.02 | 1259.08±4.96 | 2280.72±17.46 | 1441.83±14.17 | 1323.48±17.62 | 1321.22±16.33 | 1262.24±15.73 | 1262.83±13.23 | 15 | 12 | 18 | 13 | 13 | 12 |
| posteriorcingulate | 1201.55±14.16 | 1233.58±15.34 | 1047.58±75.93 | 1026.30±71.22 | 880.80±12.48 | 879.96±11.38 | 991.44±17.67 | 988.78±15.27 | 11 | 11 | 9 | 10 | 8 | 10 |
| isthmuscingulate | 1162.84±18.58 | 965.42±23.77 | 1163.28±14.27 | 880.08±13.45 | 908.40±14.19 | 905.39±9.51 | 811.92±11.48 | 811.17±11.4 | 10 | 10 | 11 | 8 | 9 | 6 |
| parsorbitalis | 614.84±12.54 | 812.89±12.29 | 814.00±15.14 | 999.42±14.25 | 722.72±16.16 | 720.74±14.81 | 1091.76±27.64 | 1097.96±16.37 | 6 | 9 | 7 | 9 | 7 | 11 |
| bankssts | 1062.55±4.37 | 779.39±6.86 | 1080.88±5.27 | 1053.72±8.13 | 1172.76±7.57 | 1172.87±7.57 | 986.96±9.41 | 987.13±8.36 | 9 | 8 | 10 | 11 | 12 | 9 |
| caudalanteriorcingulate | 494.95±3.16 | 630.63±12.14 | 483.98±21.19 | 867.88±19.80 | 634.80±9.77 | 636.13±7.5 | 886.04±18.60 | 886.78±19.24 | 4 | 7 | 4 | 7 | 5 | 8 |
| parahippocampal | 633.92±7.93 | 612.87±9.04 | 608.15±6.92 | 570.70±7.39 | 715.36±12.14 | 712.78±8.34 | 602.52±9.69 | 603.26±6.2 | 7 | 6 | 6 | 5 | 6 | 5 |
| rostralanteriorcingulate | 846.79±12.38 | 515.97±11.57 | 980.58±21.11 | 772.00±17.51 | 1006.76±23.92 | 1011.17±18.53 | 815.72±16.81 | 817.35±14.72 | 8 | 5 | 8 | 6 | 10 | 7 |
| temporalpole | 509.34±17.80 | 469.26±22.66 | 524.48±24.01 | 506.15±20.31 | 468.72±22.12 | 467.65±22.49 | 564.04±28.43 | 565.52±28.99 | 5 | 4 | 5 | 4 | 3 | 4 |
| entorhinal | 469.68±13.333 | 438.24±11.35 | 472.92±20.49 | 425.68±12.07 | 484.36±25.49 | 482.91±25.77 | 514.08±16.93 | 515.57±16.81 | 3 | 3 | 3 | 2 | 4 | 3 |
| transversetemporal | 378.05±5.91 | 302.63±5.34 | 427.55±8.52 | 298.30±6.73 | 406.60±16.42 | 403.87±13.82 | 308.80±12.48 | 305.61±5.83 | 2 | 2 | 2 | 1 | 2 | 1 |
| frontalpole | 219.82±7.04 | 296.03±11.25 | 320.10±6.33 | 490.45±9.54 | 249.92±7.46 | 248.83±5.34 | 356.32±20.55 | 352.83±13.41 | 1 | 1 | 1 | 3 | 1 | 2 |
| Total | 83723.42±433.67 | 86121.82±474.84 | 91469.15±611.62 | 90930.48±671.70 | 91200.48±1113.47 | 90934.43±609.31 | 90839.36±671.70 | 90690.48±527.53 |  |  |  |  |  |  |

Unit: millimeters per square; Red color: the three thickest brain regions; Blue color: the three thinnest brain regions; Green color: brain thickness after removing two sessions with excessive head movement

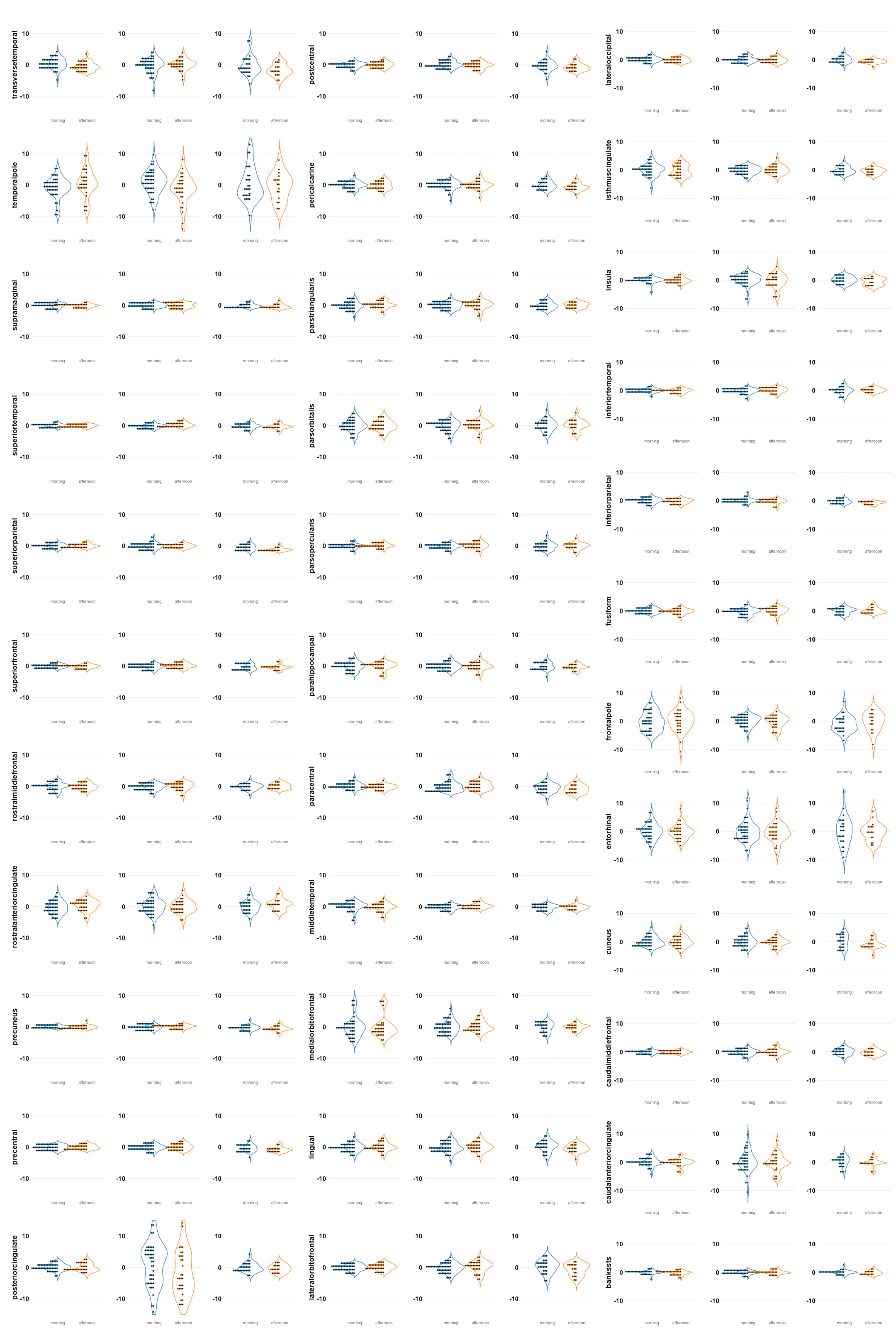

**Fig. S5. The percentage change distribution of time-of-day in the surface area of DK atlas-defined brain regions**. From left to right, sub1, sub2, and sub3.

**Cortical brain volume**

The average and standard deviation of the total gray matter volumes, total white matter volumes, and the CSF volumes including the left and right hemispheres are described in **Table S4**.

**Table S4.** The values (Mean ± SD) of volumes of the different brain apartments

|  | **Sub1** | **Sub2** | **Sub3** |
| --- | --- | --- | --- |
| **Gray Matter** | 477.87 ± 3.88 | 548.77 ± 3.89 | 507.50 ± 6.67 |
| **Left - GM** | 236.60 ± 2.10 | 275.81 ± 2.21 | 254.96 ± 3.57 |
| **Right - GM** | 241.27 ± 2.20 | 272.96 ± 2.03 | 252.54 ± 3.40 |
| **White Matter** | 453.74 ± 2.99 | 515.36 ± 3.32 | 445.01 ± 4.01 |
| **Left - WM** | 224.54 ± 1.55 | 258.50 ± 1.62 | 223.28 ± 2.68 |
| **Right - WM** | 229.20 ± 1.75 | 256.86 ± 1.90 | 221.73 ± 1.92 |
| **CSF** | 0.81 ± 0.04 | 1.22 ± 0.04 | 1.05 ± 0.04 |
| **IntraCranialVol (ICV)** | 1525.148 | 1658.298 | 1555.418 |

The unit of all values is cubic centimeters.

***Mean cortical brain volume across sessions***

For three subjects, brain volume of different regions varied from 731.53 to 21682.50 mm³ for sub1, 981.82 to 26338.03 mm³ for sub2, and 903.65 to 26170.17 mm³ for sub3 (**Table S5**). The intracranial volume (ICV) remained consistent for each subject in all sessions (sub1: 1,525,148 mm³; sub2: 1,658,298 mm³; sub3: 1,555,418 mm³) after running the longitudinal pipeline. Ranking the volumes revealed that the superior frontal and rostral middle frontal cortices were the largest in all subjects (**Fig. S6**). Conversely, the frontal pole and transverse temporal regions were consistently the smallest across all subjects (**Fig. S6B** and **Table S5**).

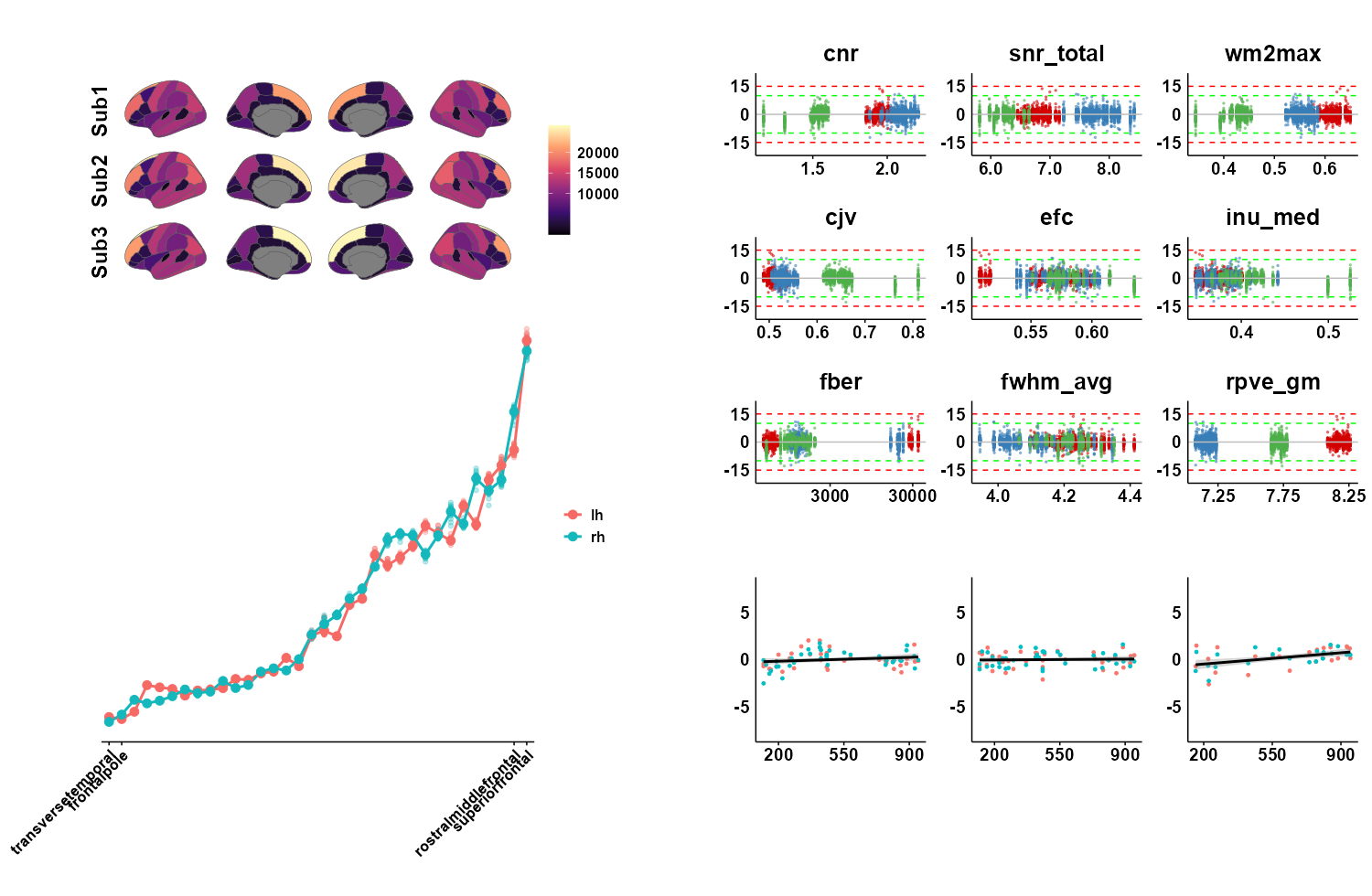

**Fig. S6. Mean cortical brain volume for each brain region, percentage change distributions along with IQMs, and percentage change of total cortical brain volume along the daylight length.** **(A)** The average brain volume size of different brain regions for each subject where the unit is cubic millimeters. **(B)** Brain regions from the smallest to the largest (data from sub1). All three subjects have the same the most two smallest and largest brain volumes although the brain regions ranking in the middle vary. **(C)** The distribution of the percentage changes along the IQMs, where the red, blue, and green dots represent subjects 1, 2, and 3, respectively. **(D)** Percentage changes of total cortical brain volumes along daylight length (x-axis with a unit of minute) for each subject.

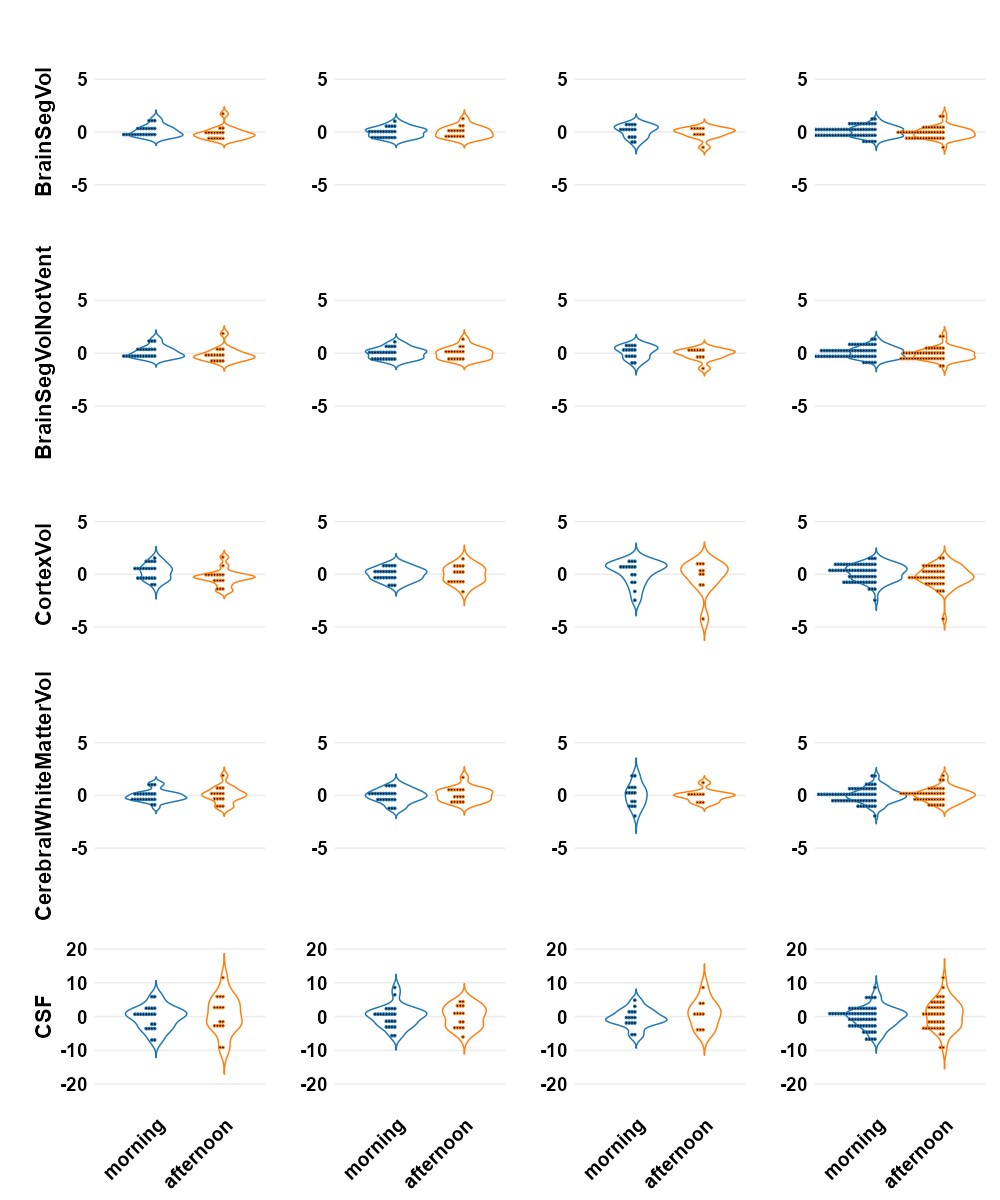

**Fig. S7. The percentage change distribution of time-of-day in different brain compartments.** From left to right, sub1, sub2, and sub3.

**Table S5.** Mean volume size across sessions for each subject based on DK atlas

|  | Sub1 | | Sub2 | | Sub3 | | | | Ranking | | | | | |
| --- | --- | --- | --- | --- | --- | --- | --- | --- | --- | --- | --- | --- | --- | --- |
|  | L | R | L | R | L | L_qc | R | R_qc |  |  |  |  |  |  |
| superiorfrontal | 21682.50±228.56 | 21111.29±232.77 | 26338.03±316.78 | 25311.47±272.31 | 24289.32±379.55 | 24305.43±385.33 | 26131.32±385.48 | 26170.17±342.58 | 34 | 34 | 34 | 34 | 34 | 34 |
| rostralmiddlefrontal | 15672.11±199.03 | 17776.50±233.63 | 19785.47±264.41 | 19406.60±282.46 | 16233.72±271.59 | 16232.78±257.08 | 20960.88±359.22 | 20981.52±357.34 | 33 | 33 | 33 | 33 | 33 | 33 |
| inferiorparietal | 11576.18±132.66 | 14100.29±213.56 | 14075.00±156.12 | 17178.70±193.70 | 13430.20±295.56 | 13488.7±202.35 | 15042.36±244.19 | 15096.7±160.82 | 29 | 32 | 27 | 32 | 28 | 32 |
| precentral | 14832.47±215.09 | 14033.61±184.83 | 14455.42±239.72 | 13568.90±185.98 | 14976.16±340.62 | 15018.09±313.63 | 13176.80±133.94 | 13188±132.42 | 32 | 31 | 31 | 30 | 32 | 31 |
| superiorparietal | 14024.03±190.38 | 13431.89±199.66 | 14154.15±219.49 | 16342.77±268.20 | 14016.40±264.44 | 14046.57±227.55 | 9767.40±172.63 | 9802.7±84.39 | 31 | 30 | 28 | 31 | 30 | 24 |
| middletemporal | 10697.45±107.79 | 12289.55±233.41 | 14154.40±146.34 | 12785.55±99.39 | 12931.72±246.24 | 12976.22±152.14 | 11739.64±269.91 | 11801.96±169.38 | 26 | 29 | 29 | 27 | 27 | 28 |
| superiortemporal | 12605.97±120.07 | 11611.61±117.03 | 14346.17±164.62 | 12513.42±104.23 | 14467.56±168.38 | 14494.17±142.02 | 12123.88±200.06 | 12168.7±128.92 | 30 | 28 | 30 | 26 | 31 | 29 |
| precuneus | 9764.34±130.43 | 11059.63±125.53 | 11735.23±149.76 | 11845.23±127.12 | 9621.24±159.93 | 9649.22±129.38 | 9271.88±184.19 | 9314.83±101.38 | 23 | 27 | 25 | 25 | 22 | 23 |
| inferiortemporal | 11082.16±157.90 | 10986.55±123.65 | 11140.50±121.95 | 13304.10±118.49 | 13764.96±320.79 | 13826.61±189.18 | 12969.64±359.81 | 13061.91±156.05 | 27 | 26 | 24 | 29 | 29 | 30 |
| lateraloccipital | 10406.61±143.45 | 10979.68±102.18 | 15233.73±179.73 | 12971.02±159.67 | 12835.80±259.75 | 12889.39±185.87 | 11083.12±227.78 | 11128.87±170.03 | 25 | 25 | 32 | 28 | 26 | 26 |
| postcentral | 9348.50±147.89 | 10742.92±155.66 | 10967.98±157.69 | 9967.92±197.43 | 10785.32±201.29 | 10790.04±202.23 | 10745.48±152.00 | 10775.26±116.18 | 22 | 24 | 22 | 22 | 23 | 25 |
| supramarginal | 11510.89±119.98 | 9937.08±129.61 | 10990.90±108.07 | 11396.92±109.85 | 11917.92±221.27 | 11935.35±199.08 | 8884.00±126.29 | 8909±94.55 | 28 | 23 | 23 | 23 | 25 | 22 |
| fusiform | 9908.03±129.52 | 9254.16±59.11 | 11832.27±105.34 | 11418.80±82.60 | 11475.36±161.36 | 11506.52±109.56 | 11302.12±177.69 | 11344.96±102.11 | 24 | 22 | 26 | 24 | 24 | 27 |
| insula | 7500.24±88.89 | 8040.53±106.68 | 7727.90±90.60 | 7891.65±129.37 | 6723.16±157.48 | 6719±134.71 | 6229.56±93.84 | 6219.48±76.67 | 21 | 21 | 19 | 19 | 18 | 17 |
| lateralorbitofrontal | 7170.05±75.09 | 7502.58±92.80 | 9204.55±120.87 | 8908.15±95.30 | 7950.60±137.20 | 7963.78±130.21 | 8184.24±249.14 | 8238.04±143.54 | 20 | 20 | 21 | 21 | 21 | 21 |
| lingual | 5439.37±58.64 | 6595.03±78.37 | 8258.75±104.77 | 8368.73±128.99 | 7172.48±87.60 | 7187.57±72.98 | 7315.76±118.09 | 7323.78±119.2 | 17 | 19 | 20 | 20 | 20 | 19 |
| medialorbitofrontal | 5741.00±271.68 | 6108.95±160.75 | 6214.08±168.57 | 7351.30±118.96 | 5260.40±99.83 | 5267.74±97.2 | 6818.00±106.89 | 6825.96±100.09 | 19 | 18 | 16 | 18 | 17 | 18 |
| caudalmiddlefrontal | 5478.87±106.81 | 5524.84±91.60 | 6873.95±136.37 | 5787.77±124.94 | 6883.72±137.67 | 6876.61±138.94 | 7954.16±122.51 | 7960.61±108.72 | 18 | 17 | 17 | 17 | 19 | 20 |
| parstriangularis | 3791.97±39.01 | 4177.34±59.73 | 4669.40±60.27 | 4503.95±49.24 | 3173.12±62.52 | 3174.35±60.64 | 3816.80±71.55 | 3820.65±71.49 | 15 | 16 | 15 | 15 | 14 | 16 |
| paracentral | 3495.71±61.89 | 3673.55±62.34 | 3731.32±74.63 | 4267.68±91.45 | 3910.32±99.76 | 3931.3±66.68 | 3498.16±71.18 | 3513.13±48.79 | 14 | 15 | 14 | 14 | 16 | 12 |
| parsopercularis | 4250.50±51.46 | 3559.08±45.70 | 7019.18±90.97 | 4616.52±51.46 | 3891.60±90.18 | 3898.83±75.69 | 3556.40±82.10 | 3570.3±64.98 | 16 | 14 | 18 | 16 | 15 | 14 |
| posteriorcingulate | 3397.16±35.80 | 3492.71±42.04 | 2781.82±134.48 | 2939.45±143.18 | 2353.36±46.37 | 2356.87±45.16 | 2573.00±34.69 | 2576.7±25.87 | 13 | 13 | 7 | 11 | 7 | 10 |
| cuneus | 2585.03±32.35 | 2976.34±56.50 | 3720.57±57.59 | 4042.20±131.89 | 2980.24±43.08 | 2982.7±43.88 | 3564.12±60.93 | 3560.52±55.09 | 8 | 12 | 13 | 13 | 12 | 13 |
| temporalpole | 3035.13±59.49 | 2760.03±54.80 | 2791.30±73.46 | 2667.32±78.98 | 2121.76±105.15 | 2129.17±106.34 | 3125.08±103.29 | 3138.78±92.87 | 11 | 11 | 8 | 9 | 4 | 11 |
| isthmuscingulate | 3089.97±34.62 | 2593.61±36.41 | 3291.43±37.19 | 2342.35±38.02 | 2404.12±34.66 | 2410.78±27.01 | 2141.32±29.69 | 2141.17±31 | 12 | 10 | 12 | 5 | 9 | 4 |
| parsorbitalis | 2174.68±31.89 | 2509.95±24.58 | 2813.12±60.02 | 3597.50±65.05 | 2415.80±45.18 | 2415.83±47.19 | 3617.24±75.25 | 3629.52±64.49 | 4 | 9 | 9 | 12 | 10 | 15 |
| entorhinal | 2497.79±71.94 | 2398.34±53.14 | 2517.88±60.81 | 2201.65±82.37 | 2359.68±79.15 | 2365.61±79.47 | 2491.72±94.46 | 2506.96±77.38 | 6 | 8 | 5 | 4 | 8 | 7 |
| pericalcarine | 2455.71±44.89 | 2320.18±82.11 | 2746.57±60.56 | 2610.25±102.94 | 2131.72±84.97 | 2130.96±86.2 | 2166.84±85.67 | 2165.22±89.28 | 5 | 7 | 6 | 7 | 5 | 5 |
| parahippocampal | 2541.18±34.97 | 2140.74±32.26 | 2249.78±37.96 | 1970.12±30.15 | 2170.92±48.32 | 2176.61±39.37 | 1958.12±51.49 | 1967.57±37.76 | 7 | 6 | 4 | 3 | 6 | 3 |
| caudalanteriorcingulate | 1288.61±19.20 | 1945.45±23.84 | 1195.42±45.00 | 2534.57±46.79 | 1850.16±33.26 | 1853.57±31.29 | 2560.84±39.48 | 2564.48±35.8 | 3 | 5 | 2 | 6 | 3 | 9 |
| bankssts | 2627.76±20.22 | 1895.66±34.94 | 2841.78±29.21 | 2762.02±27.09 | 2797.00±67.25 | 2810.22±43.56 | 2404.12±54.32 | 2403.04±30.81 | 9 | 4 | 10 | 10 | 11 | 6 |
| rostralanteriorcingulate | 2748.97±34.74 | 1748.97±22.87 | 3105.75±54.36 | 2626.68±28.21 | 3135.68±42.09 | 3135.78±43.52 | 2532.28±25.97 | 2530.22±26.08 | 10 | 3 | 11 | 8 | 13 | 8 |
| frontalpole | 886.21±16.55 | 1130.08±19.56 | 1146.25±19.14 | 1744.03±23.07 | 1029.68±16.10 | 1031.52±14.8 | 1482.00±36.26 | 1486.52±32.68 | 1 | 2 | 1 | 2 | 1 | 2 |
| transversetemporal | 996.84±15.02 | 731.53±11.96 | 1287.97±18.45 | 981.82±12.71 | 1130.76±40.34 | 1136.13±33.81 | 900.24±20.33 | 903.65±16.76 | 2 | 1 | 3 | 1 | 2 | 1 |

Unit: cubic millimeter; Red color: the three thickest brain regions; Blue color: the three thinnest brain regions; Green color: brain thickness after removing two sessions with excessive head movement

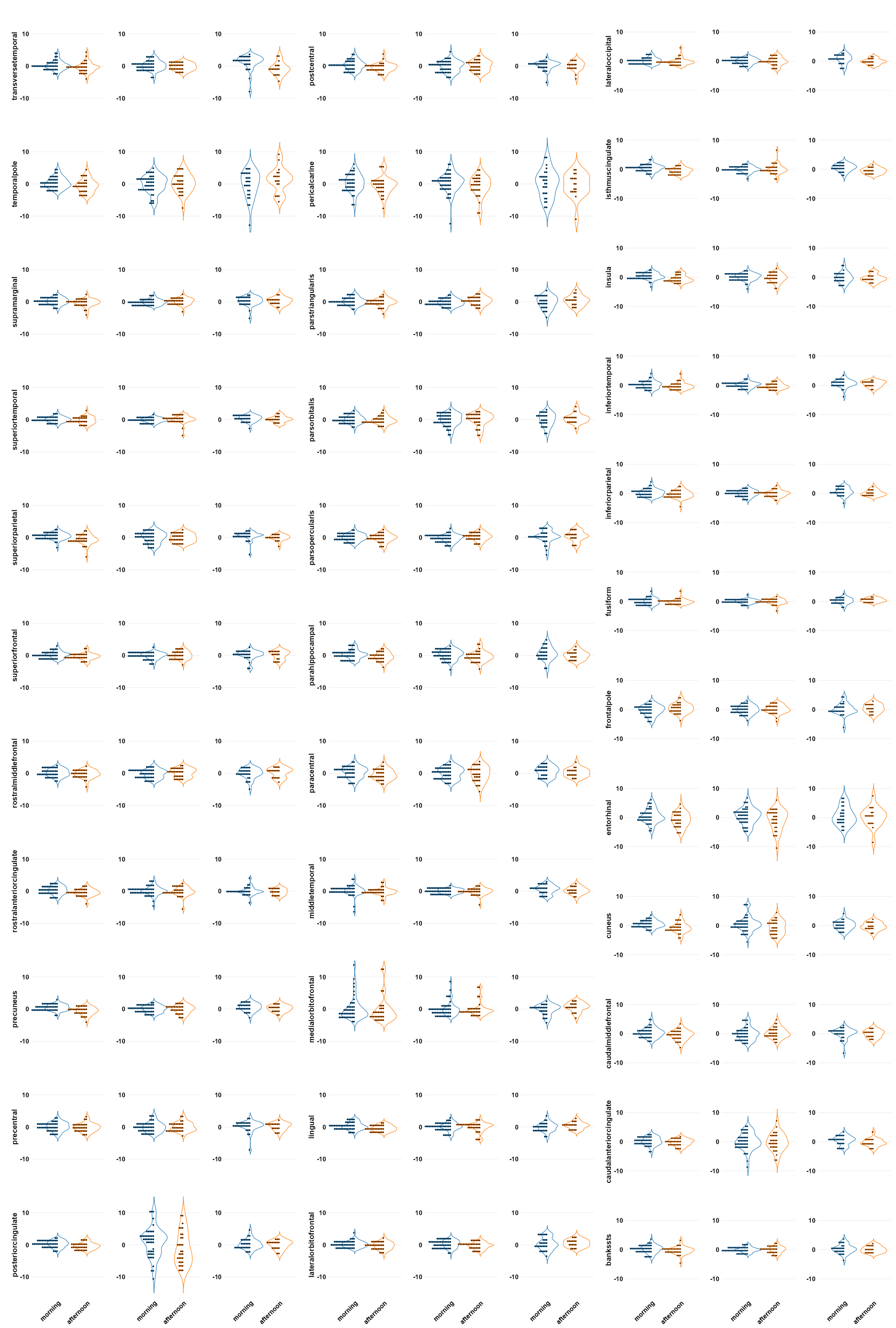

**Fig. S8. The percentage change distribution of time-of-day in the cortical brain volume of DK atlas-defined brain regions** From left to right, sub1, sub2, and sub3.

**Subcortical volumes**

Subcortical structures encompassed the thalamus, caudate nucleus, putamen, pallidum, hippocampus, amygdala, and accumbens area. Subcortical brain volumes ranged from 721.54 to 9063.16 mm^3^ for sub1, 607.53 to 10865.75 mm^3^ for sub2, and 497.60 to 7692.36 mm^3^ for sub3 (**Table S6**). The putamen and and thalamus consistently ranked as the largest subcortical structures, see **Fig. S9** and **Table S6**. Additionally, the pallidum, amygdala, and accumbens area constantly ranked as the smallest subcortical structures across all three subjects (**Fig. S9** and **Table S6**). **Table S6** depicts the average value and standard deviation of the subcortical structures along with the cerebellum, ventricles, brain stem, corpus callosum (CC), and CSF.

**Table S6.** The mean and standard deviation of the non-cortical brain volumes

|  |  | **Sub1** | **Sub2** | **Sub3** |
| --- | --- | --- | --- | --- |
| **Cerebellum Cortex** | L | 49513.90±902.94 | 69978.78±1144.13 | 54862.35±575.78 |
|  | R | 47662.64±787.88 | 69575.09±1216.57 | 54736.96±638.23 |
| **Cerebellum White Matter** | L | 12248.32±164.52 | 18075.81±289.03 | 13059.24±249.25 |
|  | R | 11890.80±144.38 | 17155.27±219.62 | 13061.13±219.39 |
| **Thalamus** | L | 9063.16±94.43 | 10865.75±76.00 | 7692.36±101.40 |
|  | R | 8628.22±120.11 | 10467.27±129.49 | 7426.36±99.28 |
| **Putamen** | L | 6174.55±43.24 | 5999.58±72.23 | 5111.26±66.52 |
|  | R | 6458.49±75.02 | 6127.76±74.73 | 5247.80±80.06 |
| **Hippocampus** | L | 4296.58±39.89 | 4758.70±45.44 | 3679.56±56.31 |
|  | R | 4398.27±31.41 | 4701.36±54.06 | 4006.50±73.39 |
| **VentralDC** | L | 3913.89±41.81 | 4626.59±54.44 | 3941.93±50.84 |
|  | R | 4060.38±33.84 | 4745.13±48.53 | 4088.62±71.07 |
| **Caudate** | L | 3789.69±42.54 | 4526.46±62.57 | 3750.78±37.03 |
|  | R | 4227.56±48.17 | 4568.34±40.35 | 3773.77±33.81 |
| **Pallidum** | L | 2162.79±24.57 | 2129.04±28.34 | 1811.80±24.17 |
|  | R | 2463.63±31.62 | 2106.66±27.91 | 1887.07±33.07 |
| **Amygdala** | L | 1618.33±26.01 | 1639.32±39.76 | 1554.44±27.71 |
|  | R | 1883.15±27.96 | 1946.16±27.77 | 1834.68±36.75 |
| **Accumbens area** | L | 721.54±44.54 | 607.53±37.92 | 497.60±37.60 |
|  | R | 872.33±20.61 | 884.08±23.81 | 725.29±24.23 |
| **Lateral Ventricle** | L | 3793.08±139.14 | 6024.30±229.73 | 6428.02±181.62 |
|  | R | 6671.21±146.38 | 6838.28±159.56 | 6891.03±148.69 |
| **Total subcortex** | L | 27826.64±126.86 | 30526.37±154.48 | 24097.79±155.04 |
|  | R | 28931.65±208.29 | 30801.63±196.29 | 24901.48±177.96 |
| **4th Ventricle** | M | 2084.01±41.13 | 2658.65±71.46 | 1990.08±59.46 |
| **3rd Ventricle** | M | 1042.81±24.89 | 944.19±29.42 | 1110.36±27.76 |
| **Brain Stem** | M | 20490.16±127.91 | 27060.06±146.57 | 21694.58±134.01 |
| **CC** | M | 3172.85±36.88 | 4148.34±238.93 | 3605.80±64.08 |
| **CSF** | M | 812.64±36.07 | 1224.99±40.01 | 1051.72±35.34 |
| **ICV** | T | 1,525,148 | 1,658,298 | 1,555,418 |

The unit of all values is the cubic millimeters.

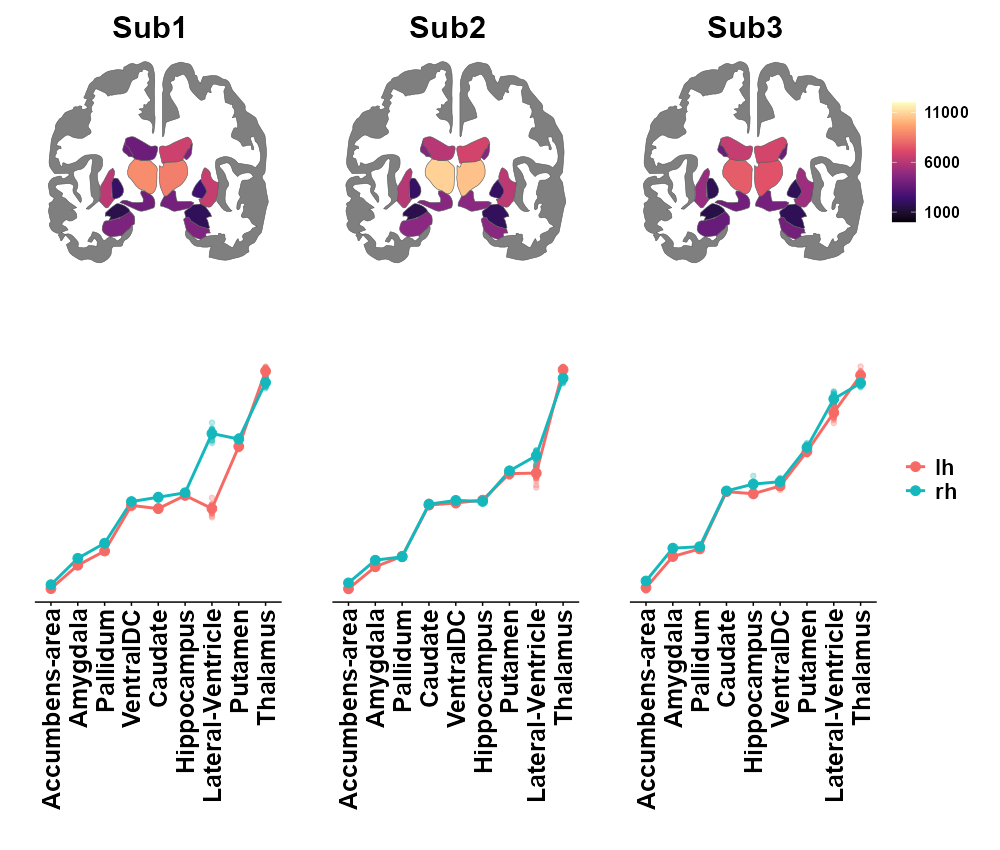

**Fig. S9. The rank of subcortical volumes based on the *aseg* atlas**.

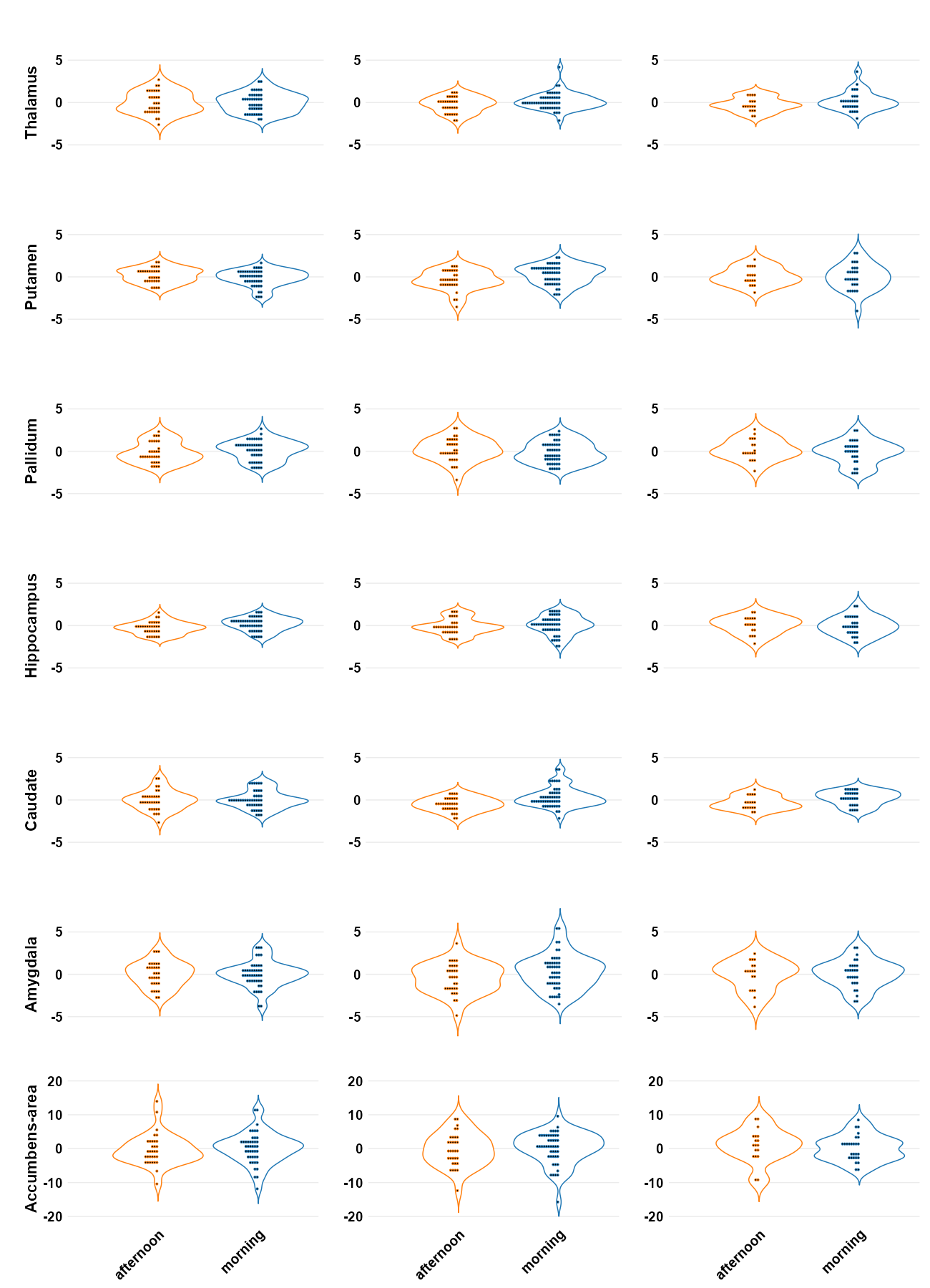

**Fig. S10. The percentage change distribution of time-of-day in the subcortical brain volumes.** From left to right, sub1, sub2, and sub3.

**Table S7.** The CV values of the non-cortical brain volume based on the DK atlas

|  | Sub1 | | Sub2 | | Sub3 | |
| --- | --- | --- | --- | --- | --- | --- |
|  | L | R | L | R | L | R |
| **Lateral-Ventricle** | 3.67 | 2.19 | 3.81 | 2.33 | 2.83 | 2.16 |
| **Cerebellum-White-Matter** | 1.34 | 1.21 | 1.60 | 1.28 | 1.91 | 1.68 |
| **Cerebellum-Cortex** | 1.82 | 1.65 | 1.63 | 1.75 | 1.05 | 1.17 |
| **Thalamus** | 1.04 | 1.39 | 0.70 | 1.24 | 1.32 | 1.34 |
| **Caudate** | 1.12 | 1.14 | 1.38 | 0.88 | 0.99 | 0.90 |
| **Putamen** | 0.70 | 1.16 | 1.20 | 1.22 | 1.30 | 1.53 |
| **Pallidum** | 1.14 | 1.28 | 1.33 | 1.32 | 1.33 | 1.75 |
| **Hippocampus** | 0.93 | 0.71 | 0.95 | 1.15 | 1.53 | 1.83 |
| **Amygdala** | 1.61 | 1.48 | 2.43 | 1.43 | 1.78 | 2.00 |
| **Accumbens-area** | 6.17 | 2.36 | 6.24 | 2.69 | 7.56 | 3.34 |
| **VentralDC** | 1.07 | 0.83 | 1.18 | 1.02 | 1.29 | 1.74 |
|  | middle | | middle | | middle | |
| **x3rd-ventricle** | 2.39 | | 3.12 | | 2.50 | |
| **x4th-ventricle** | 1.97 | | 2.69 | | 2.99 | |
| **Brain-Stem** | 0.62 | | 0.54 | | 0.62 | |
| **CSF** | 4.44 | | 3.27 | | 3.36 | |
| **CC** | 1.16 | | 5.76 | | 1.78 | |
| **CC_Posterior** | 1.44 | | 3.22 | | 1.36 | |
| **CC_Mid_Posterior** | 3.10 | | 8.39 | | 4.74 | |
| **CC_Central** | 3.10 | | 17.18 | | 2.85 | |
| **CC_Mid_Anterior** | 8.08 | | 6.11 | | 3.25 | |
| **CC_Anterior** | 4.02 | | 4.86 | | 2.02 | |

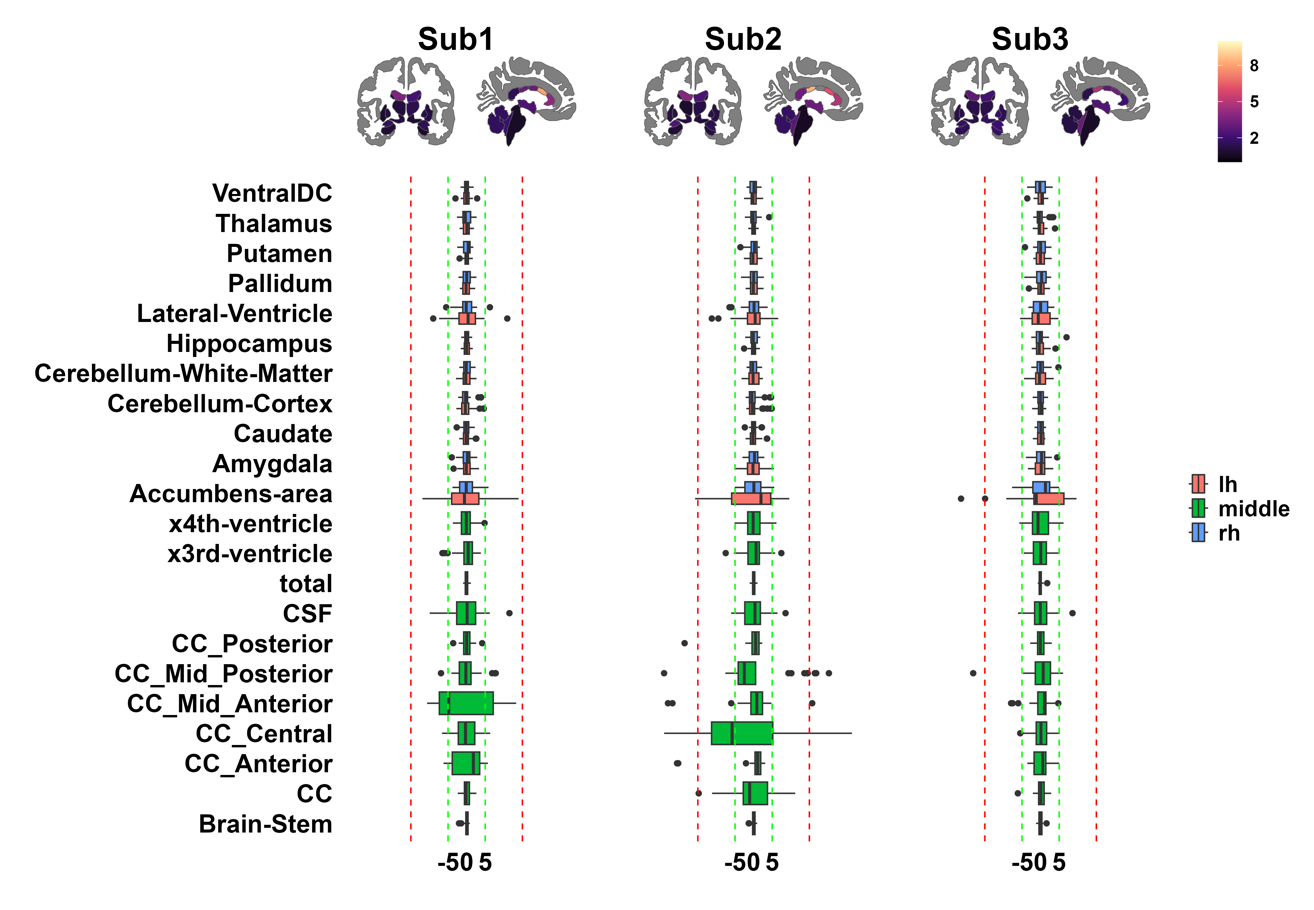

**Fig. S11. The CVs and percentage changes among non-cortical brain volumes with all sessions.**

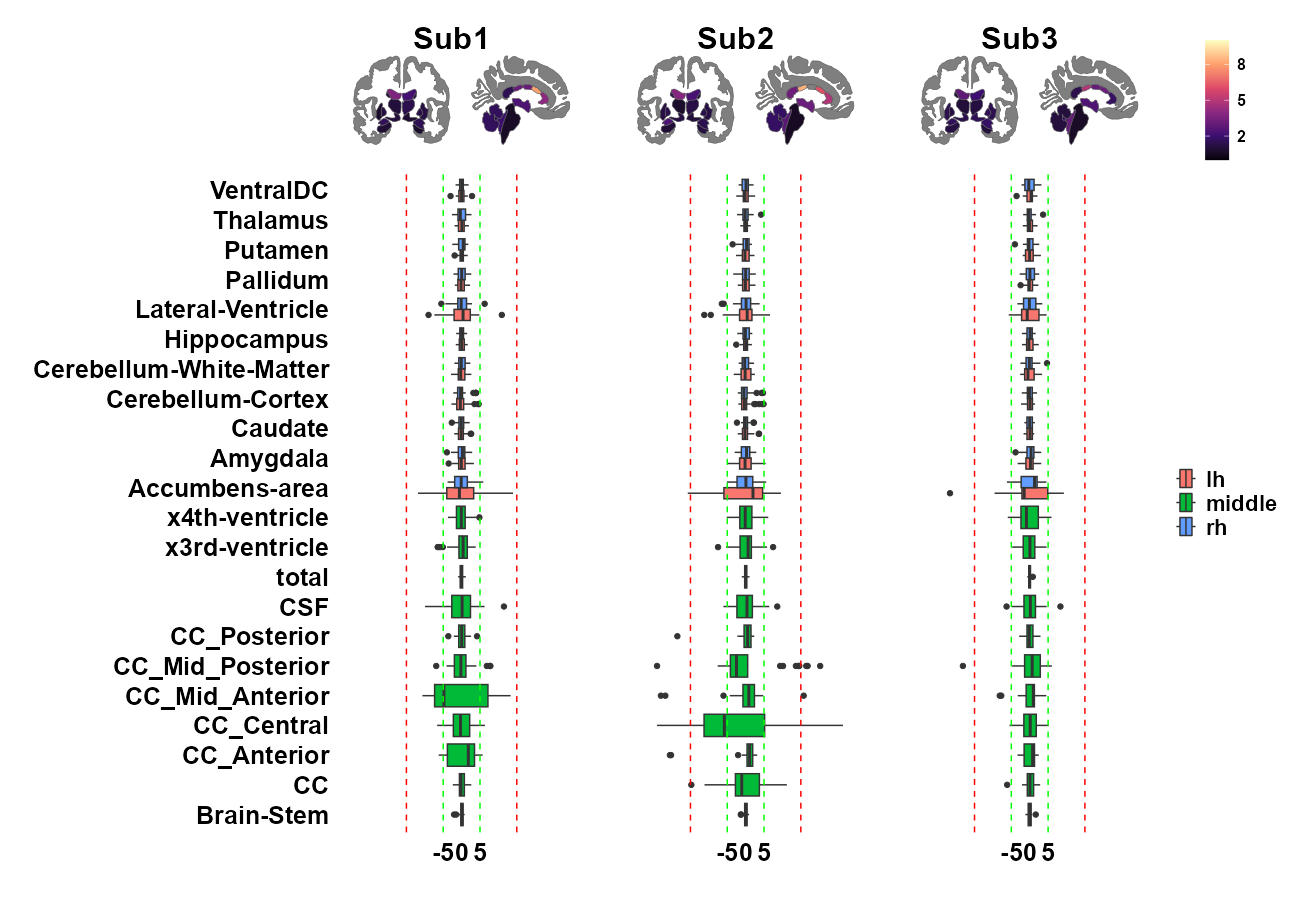

**Fig. S12. The CVs and percentage changes among non-cortical brain volumes excluding sessions 1 and 7 in sub3.**

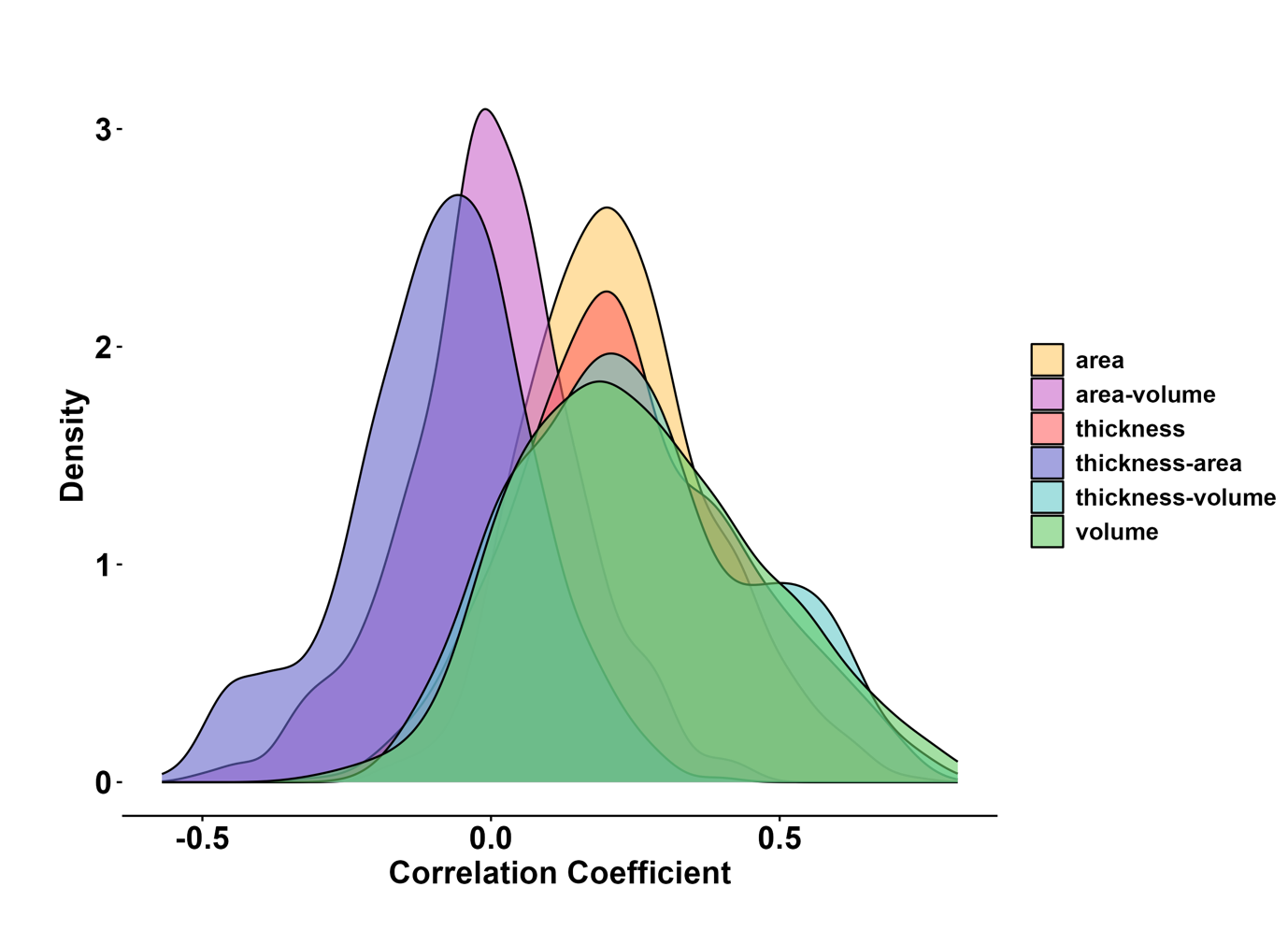

**Fig. S13.** The distribution of correlation coefficients of different phenotype associations.
